# A cold shock protein from a thermophile bacterium promotes the high-temperature growth of bacteria and fungi through binding to diverse RNA species

**DOI:** 10.1101/739763

**Authors:** Zikang Zhou, Hongzhi Tang, Weiwei Wang, Lige Zhang, Fei Su, Yuanting Wu, Linquan Bai, Sicong Li, Yuhui Sun, Fei Tao, Ping Xu

## Abstract

High temperatures deleteriously affect cells by damaging cellular structures and changing the behavior of diverse biomolecules, and extensive research about thermophilic microorganisms has elucidated some of the mechanisms that can overcome these effects and allow thriving in high-temperature ecological niches. We here used functional genomics methods to screen out a cold-shock protein (CspL) from a high-productivity lactate producing thermophile strain (*Bacillus coagulans* strain 2-6) grown at 37°C and 60°C. We subsequently made the highly striking finding that transgenic expression of CspL conferred massive increases in high temperature growth of other organisms including *E. coli* (2.4-fold biomass increase at 45°C) and the eukaryote *S. cerevisiae* (a 2.7-fold biomass increase at 34°C). Pursuing these findings, we used bio-layer interferometry assays to characterize the nucleotide-binding function of CspL *in vitro*, and used proteomics and RNA-Seq to characterize the global effects of CspL on mRNA transcript accumulation and used RIP-Seq to identify *in vivo* RNA targets of this nucleotide-binding protein (e.g. *rpoE*, and *rmf*, etc.). Finally, we confirmed that a nucleotide-binding-dead variant form of CspL does not have increased growth rates or biomass accumulation effects at high temperatures. Our study thus establishes that CspL can function as a global RNA chaperone.

## Introduction

High temperatures deleteriously affect many types of cells by damaging cellular structures such as lipid membranes and by disrupting the processing and/or functions of biomolecules such as proteins and RNA (Richter et al., 2010; Tyedmers et al., 2010; Zwirowski et al., 2017; Yost et al., 1990). Thermophilic microorganisms are of scientific and industrial interest because of the unique cellular and metabolic processes that enable them to remain viable and even thrive at high temperatures (Olson et al., 2015; Chen and Wan, 2017). While the nature of the mechanisms conferring high temperature tolerance are diverse, they are typically grouped into conceptual categories such as membrane metabolism (e.g., increased sterol content) (Daniel and Cowan, 2000), enzyme properties (e.g., increased numbers of disulfide bonds to improve protein stability) (Liu et al., 2016), post-translational processing of protein biosynthesis (e.g., phosphorylation for regulating the activity of enzymes) (Kobayashi et al., 2017) and post-transcriptional regulation of RNA metabolism (e.g., molecular chaperones including certain heat shock and cold shock proteins) (Schlesinger 1990; Treweek et al., 2015; Phadtare et al., 1998; Thieringer et al., 1998; Hudson and Ortlund 2014).

Several of the high temperature tolerance mechanisms discovered and characterized from thermophiles have been successfully exploited as novel strategies to facilitate the growth of mesophilic organisms (e.g., *Escherichia coli* and *Saccharomyces cerevisiae*) at elevated temperatures, but progress in this area has been slow (Liu et al., 2014; Vanbogelen et al., 1987). Increased tolerance for growth at elevated temperatures offers potential applied benefits such as decreasing the risk of contamination in non-sterile open fermentation, thereby substantially reducing production costs over traditional sterile fermentation (Olson et al., 2015; Wang et al., 2012; Sun et al., 2017; Zheng et al., 2017; Murata et al., 2015). Moreover, elevated fermentation temperatures can in some cases cause faster microbial growth and increased overall biomass production (Wallace-Salinas and Gorwa-Grausland 2013), making increased tolerance for high temperature growth an industrially attractive trait.

Here, building on earlier work with a high yield of L-lactate acid thermophile strain *Bacillus coagulans* 2-6 (DSM 21869). We used integrated multi-omics methods to identify candidate genes associated with high temperature response and tolerance. Heterologous expression in *Escherichia coli* of a gene encoding the cold shock protein CspL caused increased growth rates and biomass production at both 37 °C and 45 °C; cells expressing *cspL* had a 2.4-fold increase in biomass at 45 °C, while the cell morphology changed. We used bio-layer interferometry assays to confirm the nucleotide-binding function of CspL, and used RNA-Seq, RIP-Seq, and iTRAQ to characterize, respectively, the global effects of CspL on mRNA transcript and protein accumulation and the *in vivo* RNA targets of this nucleotide-binding protein. We also show that a nucleotide-binding-dead variant form of CspL does not confer increased growth or biomass production at high temperatures. Moreover, growth increases at elevated temperatures were also observed when we expressed *cspL* in *Saccharomyces cerevisiae* (2.7-fold biomass increase) and *Pseudomonas putida* (1.4-fold increase). The GFP-expressing assay shows that it has no any deleterious effects on green fluorescent protein expression or function. The fermentation validations demonstrate significant improvements in the growth and fermentation performance of two industrially relevant microorganisms. Further investigation reveals that CspL as an RNA chaperone may significantly contribute to the global transcriptional and post-transcriptional regulation and the establishment of a new proteostasis.

## Results

### Screening for high-temperature growth related genes from *Bacillus coagulans* 2-6

We previously isolated a *Bacillus coagulans* strain 2-6 (DSM 21869) from a milk processing plant in Beijing by culturing samples from soil at 55 °C (Qin et al., 2009). This thermophile can produce optically pure ʟ-lactic acid when cultured at 60 °C, but, despite having sequenced its genome, we to date know relatively little about the mechanisms that drive the high-temperature productivity of this strain (Su et al., 2014). We therefore observed the growth of *B. coagulans* 2-6 at 37 °C and 60 °C, then used RNA sequencing (RNA-Seq) and the iTRAQ proteomics method to investigate how exposure to high temperature affects, respectively, its transcriptome and proteome (Figure 1A-figure supplement 1A).

At the mRNA level, there were 170 differentially accumulated mRNA transcripts (*p* < 0.05, ≥ 2-fold change) between the cells grown at 37 °C and 60 °C (106 mRNAs increased and 64 decreased for the 60 °C samples) (Figure 1B, Supplementary file Tables 1 and 2). Prediction using the MEME program indicated that there were no significantly different transcriptional start site structures in the genes encoding the increased vs. decreased transcripts (Figure 1-figure supplement 1B). The iTRAQ experiment identified 275 proteins with differential accumulation (*p* < 0.05, ≥ 2-fold change) between *B. coagulans* 2-6 cells grown at 37 °C or 60 °C (122 proteins increased and 144 decreased for the 60 °C samples; Supplementary file Tables 3 and 4).

**Figure 1.**
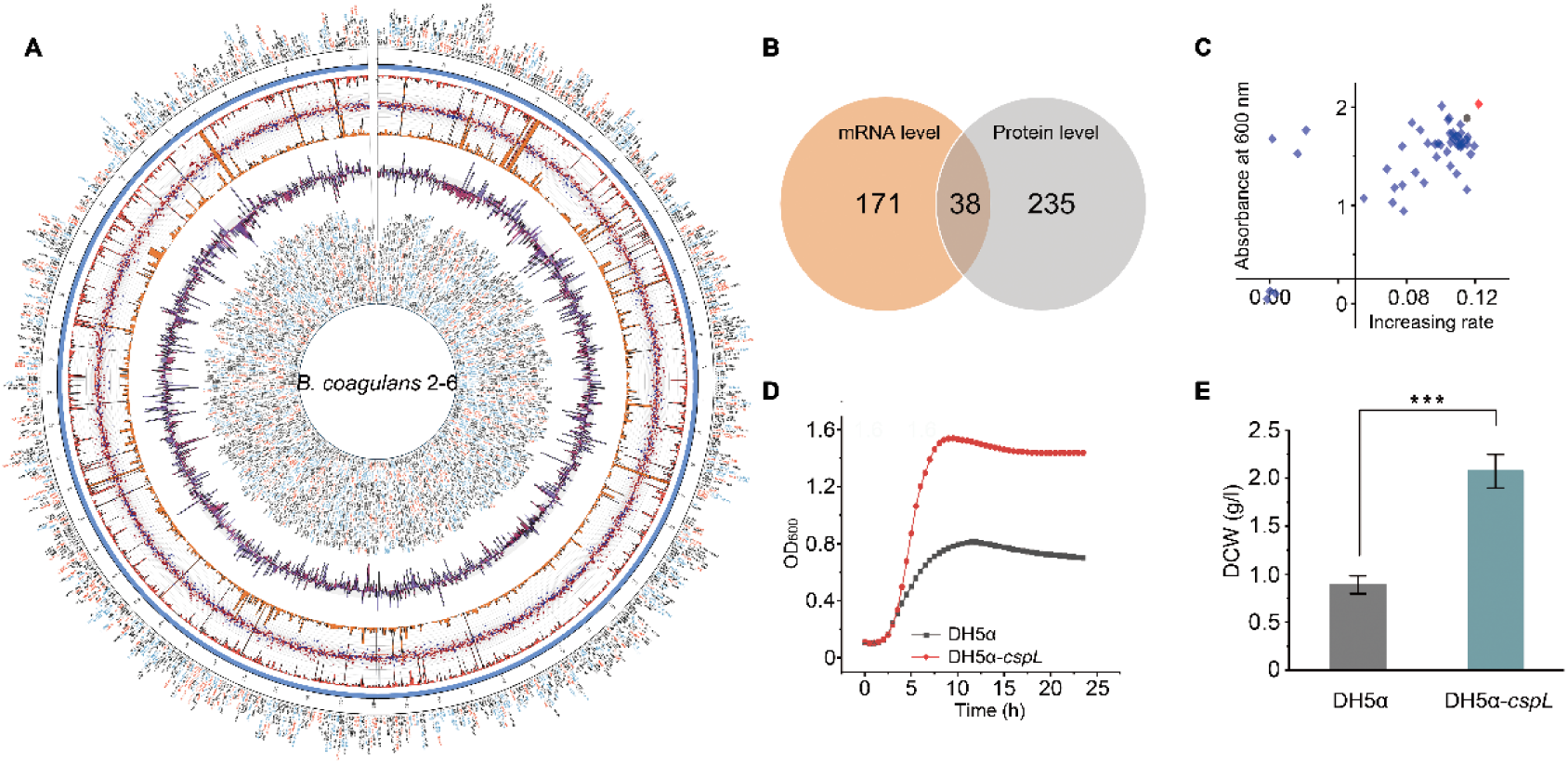
Screening and identifying candidate genes associated with high temperature growth. (A) Global view of *B. coagulans* 2-6 transcriptome and proteome analysis at 37 °C and 60 °C. Concentric circles from the periphery to the core represent the following: i) differential proteome analysis; the font color of the protein names indicates whether a protein is down-regulated (blue), up-regulated (red), or shows no significant change (gray) at 60 °C; ii) chromosomal location; iii) bar chart in red (inner orientation) representing gene expression levels at 60 °C; iv) scatter diagram representing protein expression at different conditions (red squares and blue triangles represent 60 °C and 37 °C, respectively); v) bar chart in orange (outer orientation) representing gene expression levels at the 37 °C condition; vi) bar chart representing differential expression of the same gene at the mRNA (purple) and protein (blue) levels; vii) transcriptome analysis; the font color of the gene names indicates whether a gene is down-regulated (blue), up-regulated (red), or shows no significant changes (gray) at 60 °C. (B) Venn diagram depicting the number of genes up-regulated at the mRNA and protein levels in *B. coagulans* 2-6. (C) Heterologous expression of candidate genes in *E. coli*. For 38 recombinants grown at 45 °C, the growth rate of the logarithmic phase is plotted against biomass at the stationary phase, which is represented by absorbance at 600 nm. Blue diamonds indicate different recombinants, the black circle indicates the control, and the red diamond indicates the strain heterologously expressing the *cspL* gene. (D) Functional analysis of the DH5α-*cspL* at 45 °C. Compared with the control group, the DH5α-*cspL* shows a growth advantage. (E) Differences in dry cell weight (DCW) between the DH5α-*cspL* and the control at the 45 °C culturing condition. *** *P* < 0.001 (two-tailed Student’s *t*-test).

There were 38 transcripts/proteins that were differentially expressed in both the RNA-Seq and iTRAQ datasets (24 increased and 14 decreased in the 60 °C samples; Figure 1B and Supplementary file Table 5). Both KEGG, GO, and protein interaction network analyses suggested enrichment among these candidate thermotolerant gene transcripts/proteins for functional roles related to stress responses and post-transcriptional modification processes (e.g., the global stress regulators RsbV and GsiB and the molecular chaperones Hsp20 and GroEL) (Figure 1-figure supplement 1C and D). We next used *E. coli* DH5α cells to heterologously express the 38 candidate thermotolerance-related genes monitored the growth of the cells at 45 °C (Figure 1 C-figure supplement 2). The expression of several of these candidate genes resulted in an obvious improvement in the growth of *E. coli* at 45 °C.

### Expression of a *B. coagulans* 2-6 cold shock protein improves the growth of *Escherichia coli*, *Saccharomyce cerevisiae*, and *Pseudomonas putida*

The strain heterologously expressing *cold shock protein L* (*BCO26_1317*; hereafter “*cspL*”) exhibited the most pronounced increases in growth (2.5-fold increase) and biomass production (2.4-fold increase in dry weight) when grown at 45 °C (Figure 1D and E). CspL shares 66% identity with its closest *E. coli* homolog CspA. Interestingly, we observed similar increases in growth and biomass production when we grew the strain heterologously expressing *cspL* at 37 °C (Figure 2A-C, Figure 3B). We also overexpressed *cspA* in *E. coli* and this caused significantly increased growth at both 37 °C and 45 °C compared to the empty vector control strain; however, the growth increases resulting from overexpression of *cspA* were not as pronounced as those resulting from *cspL* expression. Intriguingly, scanning electron microscopy analysis revealed a striking morphological phenotype: for samples grown at both 37 °C and 45 °C, log-phase cells expressing *cspL* were shorter than WT cells and had a ‘peanut-like’ shape (Figure 3A); note that a subsequent RNA-seq analysis showed that expression of *cspL* in *E. coli* results in the differential expression of multiple genes associated with cell size/shape, including *ftsZ*, *merC*, and *merD* (Zheng et al., 2016) (Supplementary file Table 6).

**Figure 2.**
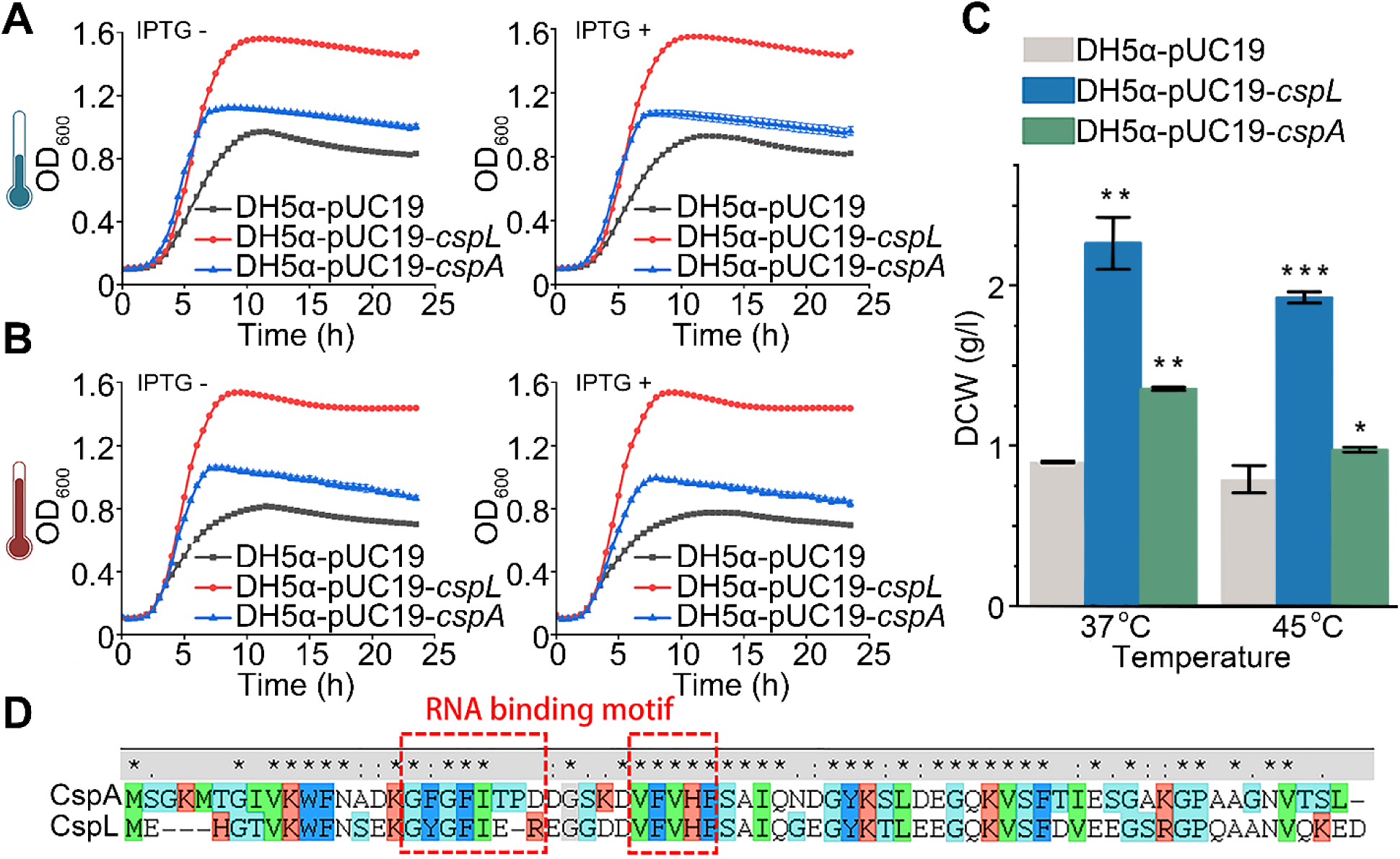
Functional comparison with homolog gene *cspA* in *E. coli*. (A) and (B), The growth curves show that CspL and CspA expressing in *E. coli* under different treatment conditions, including being induced by IPTG and cultured at 37 °C or 45 °C. (C) The bar chart shows DCW of CspL and CspA expressed in *E. coli* at 37 °C and 45 °C. Both DH5α-*cspL* and DH5α-*cspA*’s DCW had significantly improved. (D) Amino acid blast reveals *E. coli* CspA and *B. coagulans* 2-6 CspL have 66% identity and share the same RNA-binding motif. These two binding motifs are highly conserved.

**Figure 3.**
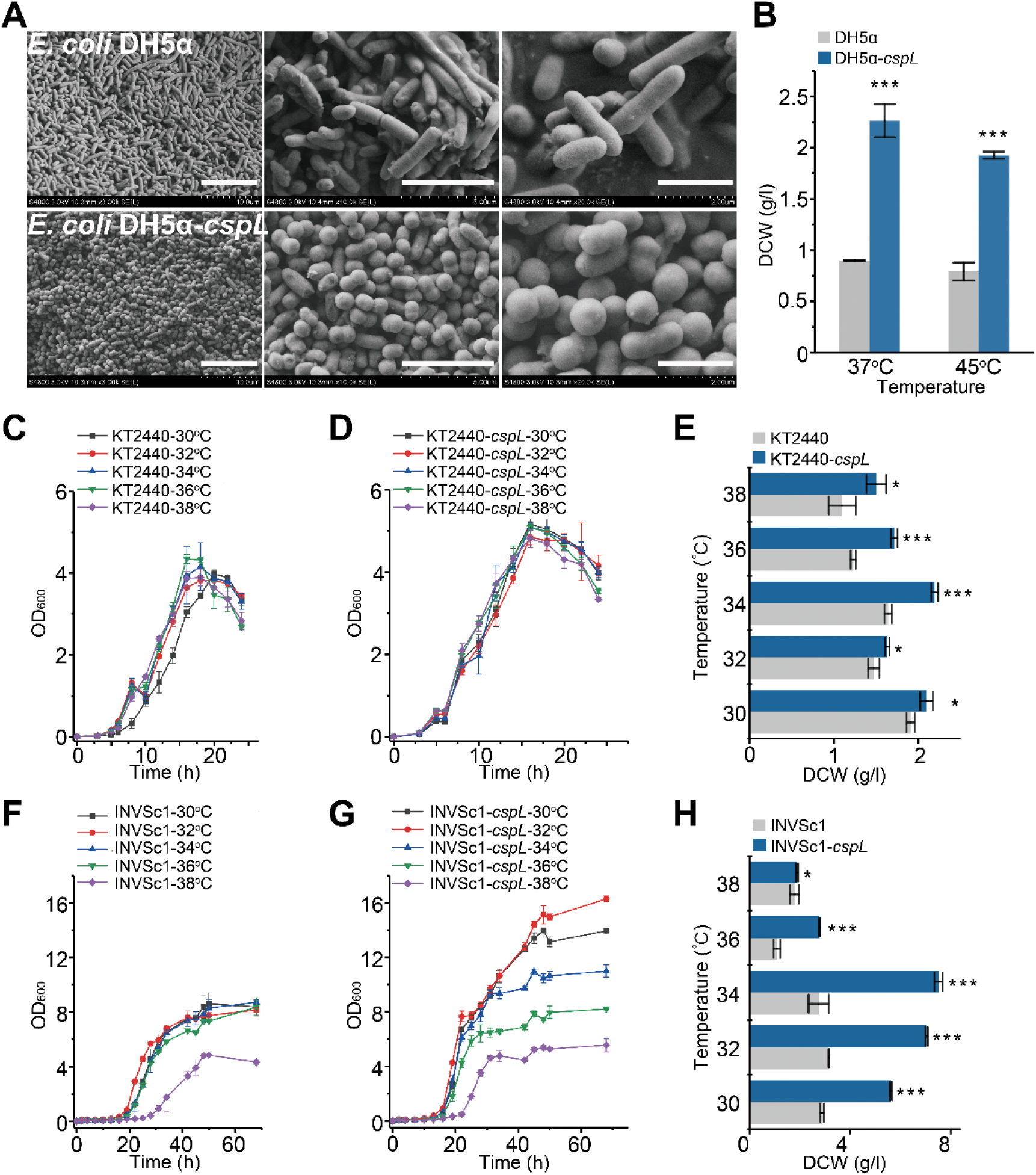
Expression of the *cspL* gene affects the cell morphology of *E. coli* and promotes the growth of *P. putida* and *S. cerevisiae* at high temperature. (A) Scanning electron microscopy images of the control (upper panel) and CspL (lower panel) strains cultured at 45 °C. (B) At both the 37 °C and 45 °C culturing conditions, statistically significant differences in dry cell weight (DCW) between strain DH5α-*cspL* and the control were observed. *** *P* < 0.001 (two-tailed Student’s *t*-test). (C) and (D) Cell growth curves of *P. putida* at different temperatures. (E) DCW of *P. putida* at different temperatures. Statistically significant differences between *P. putida* KT2440-*cspL* were observed at all temperatures tested. (F) and (G) Cell growth curves of *S. cerevisiae* at different temperatures. (H) DCW of *S. cerevisiae* at different temperatures. Statistically significant differences between *S. cerevisiae* INVSc1-*cspL* and the control were observed at all temperatures tested. * *P* < 0.05 and \*\*\**P* < 0.001 (two-tailed Student’s *t*-test). * *P*< 0.05 and \*\*\**P* < 0.001 (two-tailed Student’s *t*-test). The scale bar in image from left to right: 10 μm, 5 μm, and 2 μm.

To explore the exciting possibility that *cspL* may also help increase the high-temperature growth of other microorganisms, we heterologously expressed *cspL* in *S. cerevisiae* INVSc1 and *P. putida* KT2440 and conducted growth assays at elevated temperatures (30 °C–38 °C in 2 °C increments). In *P. putida* KT2440, both the growth rates and final biomass production of the *cspL-*expressing cells significantly outperformed those of the wild-type cells at all the tested temperatures (Figure 3C and D). At 36 °C, the *cspL-*expressing cells accumulated 1.4-fold more biomass than the empty-vector control cells (Figure 3E).

The high-temperature growth-promoting effects of *cspL* were even more obvious in *S. cerevisiae* INVSc1. Not only did the *cspL*-expressing cells significantly outperform the empty-vector control cells at all the temperatures tested, there was also an obvious ‘step’ trend in the growth of *cspL*-expressing cells at each increment in temperature (Figure 3F-G). These suggest that CspL’s growth-promoting effects in yeast are tightly linked to temperature. The most pronounced increase was observed at 34 °C, with the *cspL*-expressing cells accumulating 2.7-fold more biomass than the empty-vector control cells (Figure 3H). Note that we did not observe any ‘peanut-like’ phenotypes for *S. cerevisiae* or *P. putida* strains expressing CspL (Figure 3-figure supplement 1).

### CspL can bind RNA and its RNA-binding activity confers observed improvements in growth

As the structure of CspA has been characterized by X-ray diffraction and nuclear magnetic resonance spectroscopy (Schindelin et al., 1994; Newkirk et al., 1994), we used the CspA structure (PDB ID: 1MJC) as a template to predict that the structure of CspL was also a β-barrel comprising four flexible loops between five β-sheets that act as linkers to form a hollow structure (Figure 4A). As CspA is known to bind RNA molecules (Giuliodori et al., 2010; Rennella et al., 2017), this structural similarity suggested that CspL may function as an mRNA chaperone to confer the growth- and biomass accumulation-promoting effects that we observed in *E. coli*.

**Figure 4.**
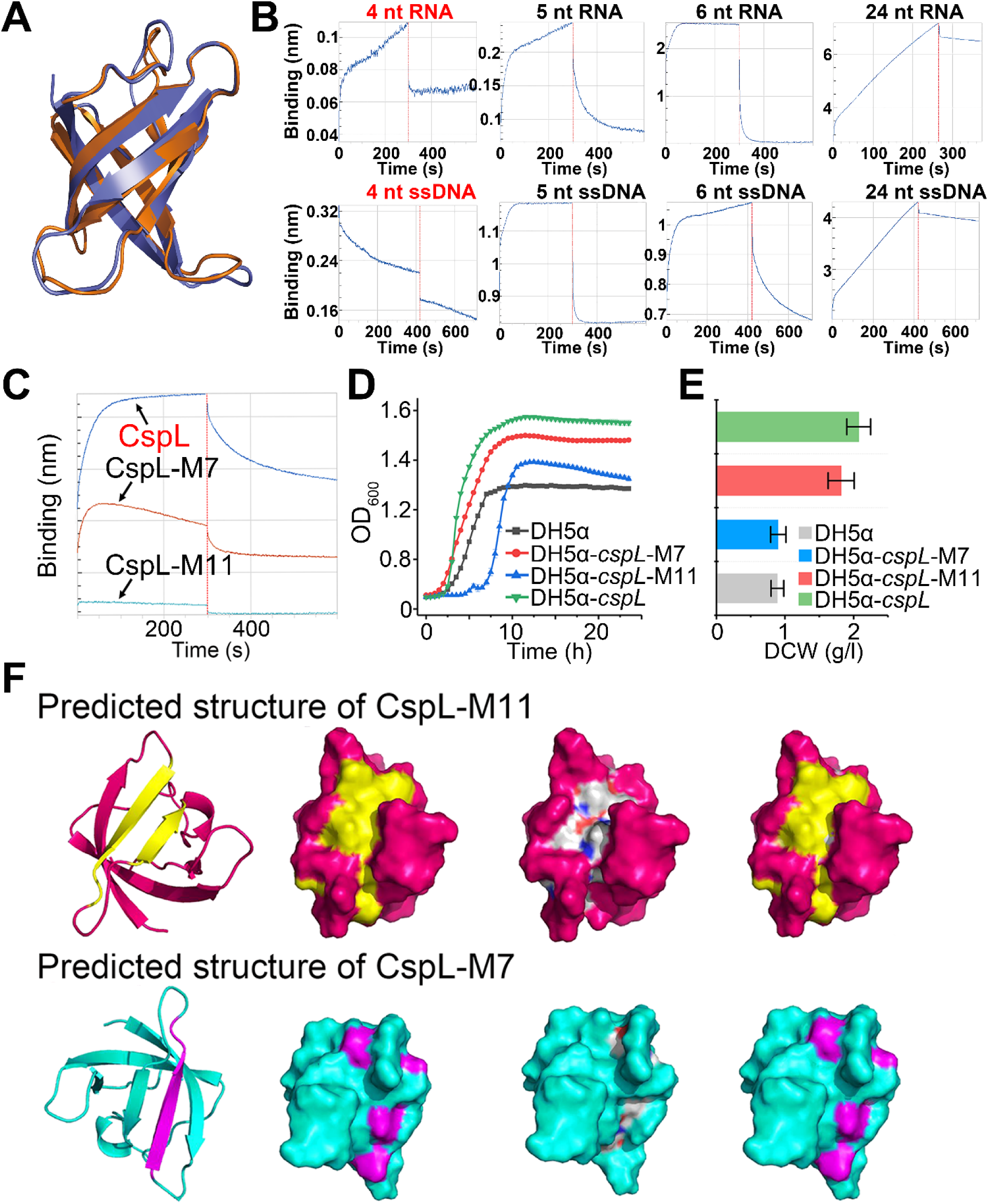
The function of CspL validation *in vitro*. (A) Comparison of the predicted CspL structure with the structure reported for *E. coli* CspA (PDB: 1mjc.1), RMSD: 0.97. (B) Binding of variously-sized RNA and ssDNA fragments to CspL. The vertical red line separates the association and disassociation steps. An absolute binding value < 0.05 nm indicates no binding signal. (C) **<u>B</u>**io **<u>L</u>**ayer **<u>I</u>**nterferometer (BLI) for testing 7 conserved amino acid-mutation. (D) and (E) The growth curves and DCW of mutations comparing against the WT and the empty vector control. (F) Two amino acid mutations were synthesized by a commercial company (Sunny, China). Sequences are listed in Supplementary Table 11. The 11 amino acid mutations predicted structure (row in red). The structure of CspL (hot pink) was predicted and a pair of anti-parallel β-sheets considered as putative ligand-binding domains (yellow) were found. When G14, Y15, G16, F17, I18, E19, R20, V26, F27, V28, and H29 in the putative ligand-binding domain were mutated to Ala (CspL-M11), the area of the ligand-binding domain significantly decreased. The ligand-binding domain mutated is colored in gray. The ligand-binding domain of CspL almost completely covered that of CspL-M11, which further demonstrated the shrink in size of the ligand-binding domain after mutation. The G14, Y15, G16, F17, I18, E19, and R20 in the putative ligand-binding domain mutated to Ala (CspL-M7, colored in blue), which slightly decreased the area of the ligand-binding domain. The ligand-binding domain mutated is colored in gray. The ligand-binding domain of CspL completely covered that of CspL-M7, which further demonstrated the shrink in size of ligand-binding domain after mutation. ** *P*< 0.01 and \*\*\**P* < 0.001 (two-tailed Student’s *t*-test).

We therefore used *in vitro* bio-layer interferometry (BLI) assays to biochemically assess the potential nucleotide-binding functions of CspL. In this analysis, we tested the interactions of HIS-tag purified CspL (20 μg/ml) with variously sized 5′-biotin-labeled RNA and DNA oligonucleotides that were immobilized to a scaffold using streptavidin. Representative data for the association and dissociation phases of these assays are presented in Figure 4B and Figure 4-figure supplement 1. We found that CspL could bind both RNA and single-stranded (ss)DNA, but it could not bind to double-stranded DNA (Figure 4-figure supplement 1C-F). It was able to bind RNA and ssDNA probes of 24, 6, and 5 nucleotides, but it was not able to bind RNA and ssDNA probes of only 4 nucleotides (Figure 4B). As the sequences of the scaffold-bound oligonucleotide targets were randomized, we speculate that CspL may function as a global chaperone for mRNA and/or ssDNA.

Using the conserved domain analysis tools available at NCBI, we identified 11 amino acids located in the putative nucleotide-binding domain of CspL and mutated all of them to alanine. BLI analysis revealed that this variant form of CspL lost its nucleotide-binding capacity (Figure 4C). Consistent with the hypothesis that CspL’s nucleotide-binding function is responsible for its growth-promoting effects, we observed no significant increases in growth rates or biomass production effects when we expressed this binding-function-dead form of CspL in *E. coli* and grew the cells at 45 °C (Figure 4D-E). Notably, we also generated a variant form of CspL with 7 residues mutated to alanine, and BLI assays showed that the nucleotide-binding function of this variant was partially impaired (Figure 4C). Growth assays at 45 °C showed that the growth and biomass production of this 7-mutated-residue variant were intermediate between wild-type CspL and the nucleotide-binding-dead variant (Figure 4D-E). We predicted the structures of mutant proteins that both two mutations could not form a stable binding pocket (Figure 4F). These findings further support our conclusion that the nucleotide-binding function of CspL confers its observed effects and raise the prospect that it may be possible to tailor the nucleotide-binding functions of CspL to promote growth at elevated temperatures yet further.

### RNA immunoprecipitation sequencing analysis reveals the interaction of CspL and hundreds of mRNA transcripts in cells

Having demonstrated the nucleotide-binding functions of CspL and having shown that the observed growth increases are mediated by such binding, we next used functional genomics profiling methods further probe CspL’s function(s) in living cells. Global transcriptome profiling with RNA-Seq revealed that *cspL* expression resulted in a striking increase in the accumulation of transcripts for numerous genes (Figure 5A): at 45 °C, transcripts for 1,160 genes (27% of all genes in the *E. coli* genome) were more abundant in *cspL-*expressing cells than in empty-vector control cells. Transcripts for 383 genes were less abundant in the *cspL*-expressing strain. In the *cspL*-expressing cells, the mRNA accumulation of *cspA* was almost 18-fold abundant than the empty-vector cells at 45 °C; however, only one tenth of abundant could be detected at 37 °C. It suggested that CspL could trigger *cspA* transcriptional expression, while *cspA* homologs had the same trends. The interaction network also shows that some genes have the closely connection with *cspA* (Figure 5B). One of them *rhlE* encodes a type of ATP-dependent RNA helicase (Jain 2008; Khemici et al., 2010), suggesting that RNA processing is regulated energetically under dual stress of temperature and presence of CspL at that exactly phase.

**Figure 5.**
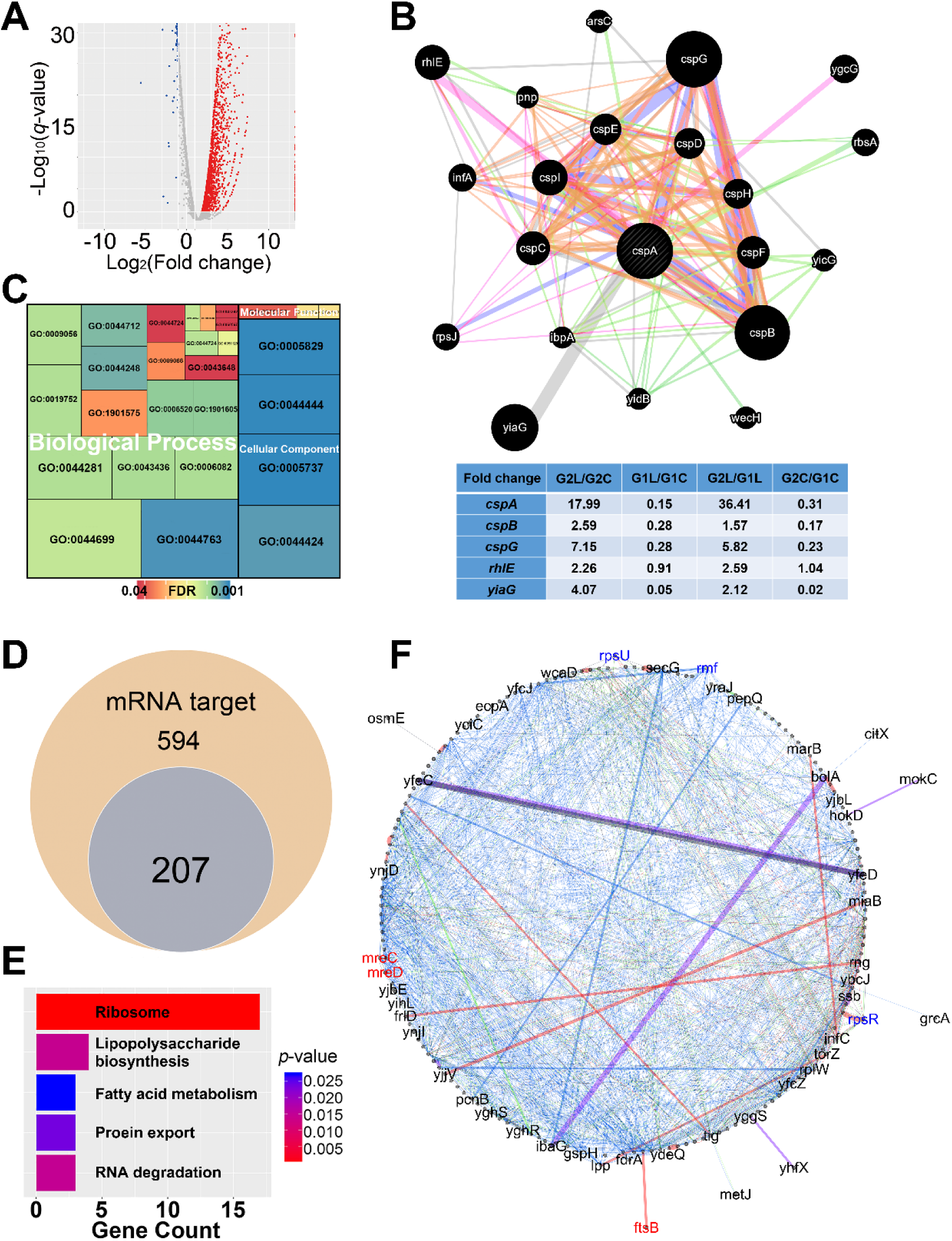
The integration function of CspL *in vivo*. (A) Volcano plot showing differentially expressed genes at the 45 °C condition. The red and blue dots indicate significantly up-regulated and down-regulated genes, respectively, and the gray dots indicate no significant difference. (B) Network of *cspL* gene interaction in *cspL*-expressing *E. coli* cells. The size of nodes represents the importance of the network interaction. Branch color indicates the type of the network. Green, genetic interactions; purple, co-expression; light blue, other; light orange, shared protein domains; and pink, physical interactions. The *cspA* was upregulated by CspL and heat stress. The table shows the closely connected genes’ expression level in different cells. G1 and G2 represent 37°C and 45°C culture condition; L and C represent *cspL*-expressing cells and empty-vector cells. (C) GO showing that most changes occurred in biological process and cellular component categories in *cspL*-expressing cells. (D) Venn diagram showing the number of *in vivo* CspL binding targets in *E. coli*. Significant enrichment was observed for 207 out of 594 mRNA fragments (*P* < 0.01). (E) CspL binding targets *in vivo*. The analysis revealed that 17 out of 207 mRNA target were related to ribosome category. The color of bars represents various *p*-values. (F) The regulation network of binding targets. The gene names in red (*mreC*, *mreD*, and *ftsB*) were directly related to cell wall synthetic processing, gene names in blue (*rpsR*, *rpsU*, and *rmf*) were responsible for ribosome synthetic processing. Branch color indicates the type of interaction. Blue, genetic interactions; purple, shared protein domain; pink, physical interactions; green, co-expression; and grey, other.

To explore translation level changes, the iTRAQ experiment distinguished 359 proteins with differential accumulation (*p* < 0.05, ≥ 2-fold change) between *E. coli* cells with or without expressing CspL grown at 45 °C (69 proteins increased and 294 decreased for the 60 °C samples). The GO analysis suggested that the most changes occurred in biological process and cellular component categories (Figure 5C). It is reasonable to connect with cell morphology changes in *E. coli* DH5α. Not surprisingly, including DnaK, DnaJ, HtpG, and related to DnaK/DnaJ proteins were both involved in high temperature stress response. It’s worth noting that heat shock regulator Sigma 32 (ơ^32^, coding by *rpoH*) and Sigma 70 (ơ^E^, coding by *rpoE*) took on some extent different trends. The ơ^32^ could not detect in control group, while heat stress stimulates ơ^32^ 2.65-fold increasing in *cspL*-expressing cells at 45 °C. The ơ^E^ expression level did not show significant alteration at two temperature conditions in empty-vector cells. However, at 45 °C and 37 °C conditions, the amounts of ơ^32^ were only 0.05 times and 0.88 times in *cspL*-expressing cells compared with empty-vector cells, respectively.

We next used RNA immunoprecipitation sequencing (RIP-seq) to identify the RNA binding targets of CspL in *E. coli* cells grown at 37 °C (we were unable to obtain RIP-Seq data from cells grown at 45 °C; recall the growth-promoting effect of *cspL* in *E. coli* cells grown at 37 °C) (Figure 5-figure supplement 1A). Consistent with the idea that CspL is a global chaperone of mRNA, the RIP-seq data demonstrated that, *in vivo*, CspL binds with the mRNA transcripts of approximately 600 genes (Figure 5D).

Our BLI binding assay data for randomized RNA sequences and our RNA-Seq data showed that expression of *cspL* significantly affected the accumulation of transcripts for fully 35% of the genes in the *E. coli* genome, supporting that CspL functions as a global chaperone for mRNA transcripts. We were cautious in our interpretation of these data sets with regard to possible trends of enrichment for particular annotation categories among the differentially accumulated transcripts or the CspL targets. We did note significant enrichment for genes related to ribosomes in the RIP-Seq data (Figure 5E), and moving forward it will be very interesting to confirm apparent trends for the preferential targeting of any distinct subsets of mRNA transcripts by CspL and to characterize any mechanism(s) that may confer any binding preferences (Figure 5E, Figure 5-figure supplement 1A). RNA decay assays proved that many binding targets of CspL were protected at different temperatures including *rmf*, *hns*, *mreC*, *mreD*, and *rpoE* (Figure 6).

**Figure 6.**
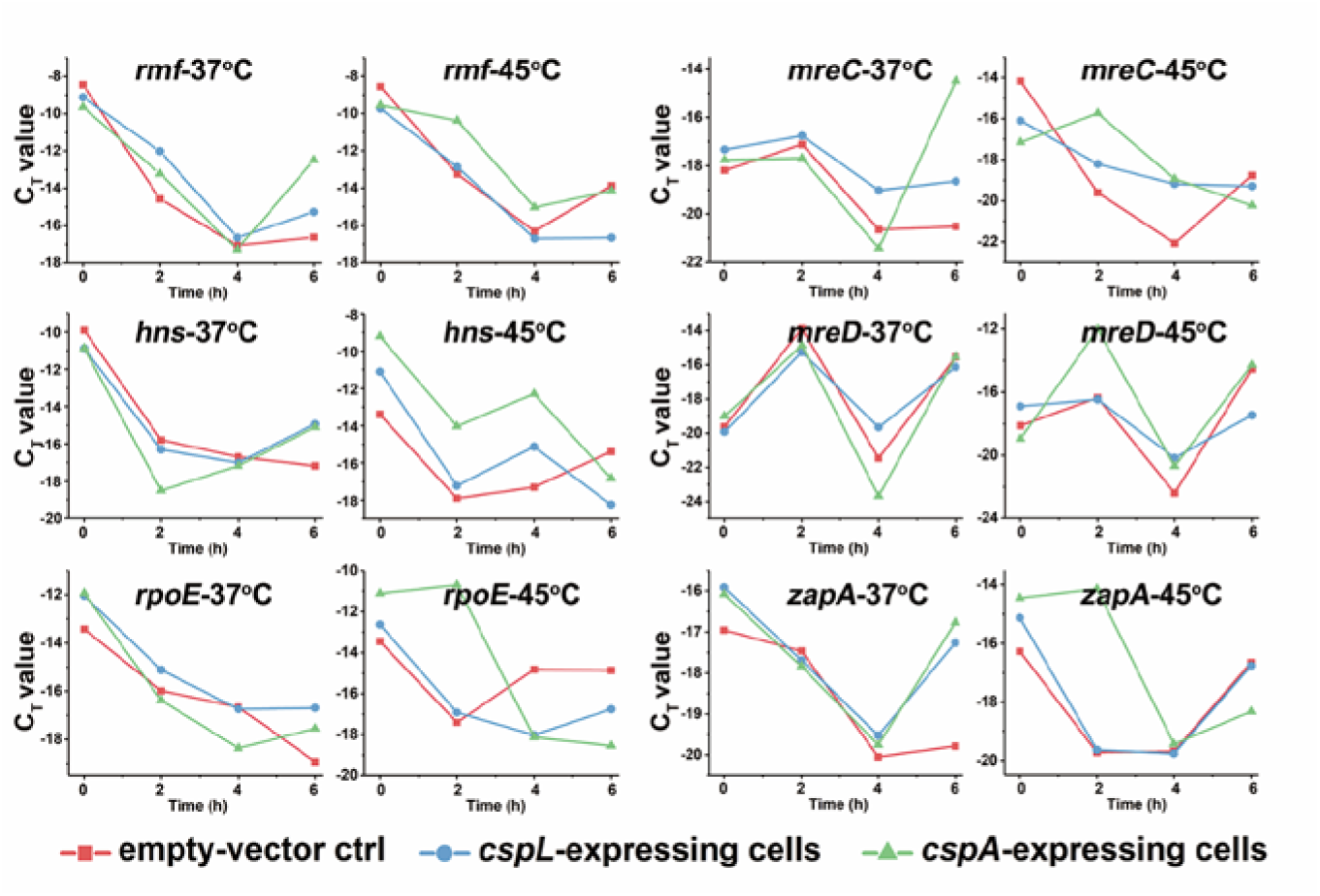
The expanding application in the *E. coli*-GFP expressing system and fermentation in the industrial microbe *A. pretiosum* and *B. licheniformis*. (A) The gene *cspL* improved the eGPF expressing strain growth at 45 °C. The strain the carried *cspL* had a shorter lag phase. (B) The HPLC analysis indicated that the production of AP-3 was increased by 160% from 29±3 mg/l to 71±6 mg/l in the culture of 31280::pLQ856-*cspL*. 1-4 represented four recombinants of 31280::pLQ856-*cspL*, respectively. (C) Comparison of D-lactate production and glucose consumption between *B. licheniformis* BN11-pEB03 (filled square) and BN11-pEB03-Pgrac100*cspL* (filled circle). Solid line and dashed line represent D-lactate production and glucose consumption. The growth of BN11-pEB03-Pgrac100*cspL* showed an obvious advantage over BN11-pEB03. The fed-batch time of BN11-pEB03-Pgrac100*cspL* was almost 5 hours ahead of the BN11-pEB03.

### CspL improves the growth and production capacity of a variety of industrial microorganisms

We further explored the practical potential of CspL in industrial biotechnology in several applications. First, when enhanced green fluorescent protein and CspL were simultaneously expressed in *E. coli* at 45 °C, the co-expression of both recombinant proteins significantly increased growth (reduced lag phase) without causing any deleterious effects on green fluorescent protein expression or function (Figure 7A). Second, the expression of CspL in *Actinosynnema pretiosum* ATCC31280 (Ning et al., 2017) significantly improved the production of the potent antitumor agent ansamitocin P-3 (a 1.6-fold increase) (Figure 7B). Third, the expression of CspL in high-temperature (50 °C) cultures of a recently engineered *Bacillus licheniformis* ATCC14580 strain (BN11) (Li et al., 2016) significantly increased both D-lactate production (1.4-fold increase) and glucose consumption (1.33-fold increase) (Figure 7C). These additional uses of CspL in fermentation applications highlight the strong industrial biotechnological promise of CspL.

**Figure 7.**
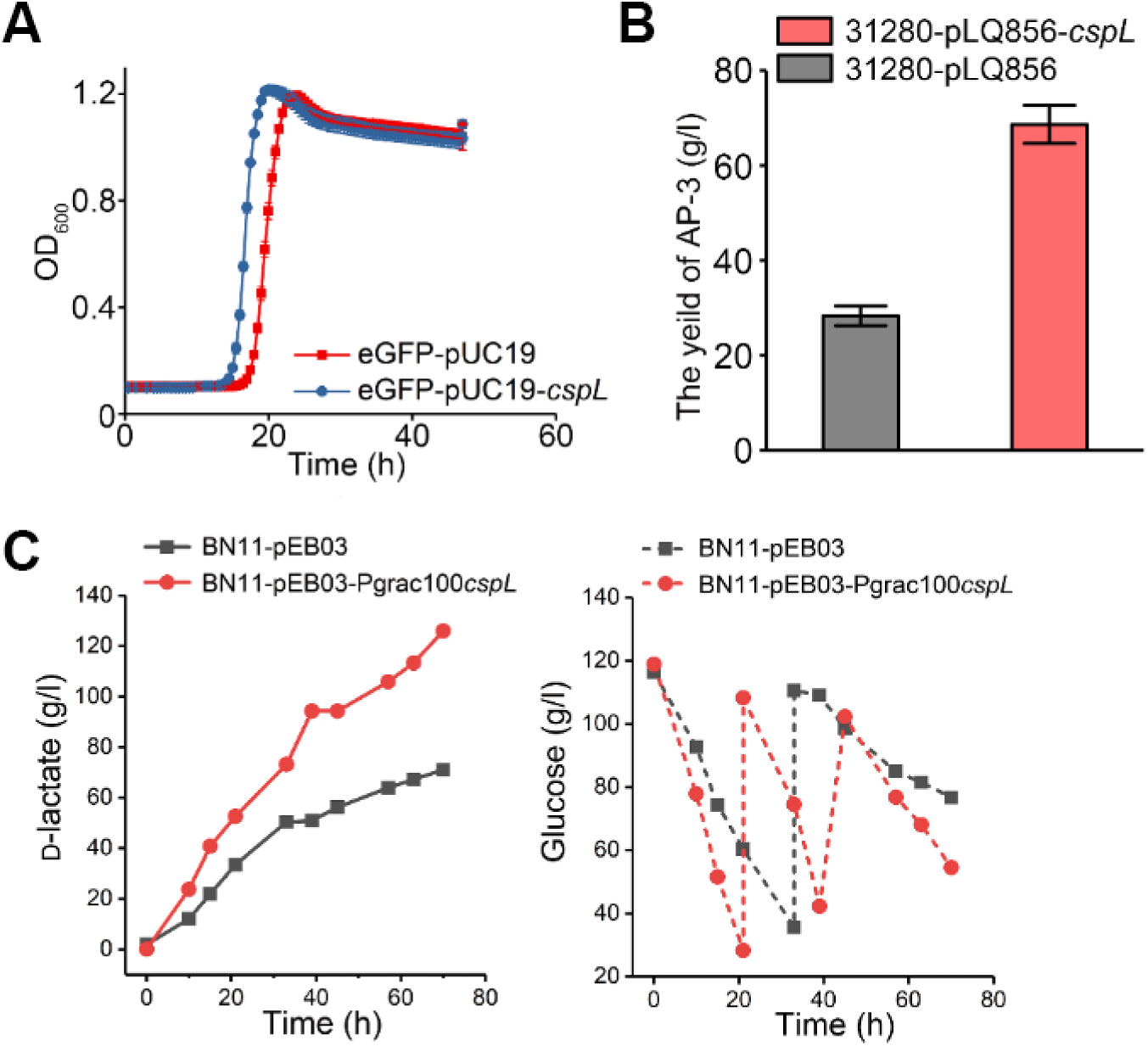
The RNA decay analysis. The RNA decay analysis were performed by RT-qPCR. Y-axe shows subtracted CT values. X-axe shows treated time points, from 0 hour point to 6 hour point.

## Discussion

Microorganisms have to survive a variety of stressful conditions. Temperature increases can damage homeostasis, interfere with essential functions, and change cellular structures. In this study, we report a novel cold shock protein, CspL. CspA from *E. coli* is the best studied bacterial cold shock protein, and the cold-temperature sensing mechanism of *cspA* mRNA is known (Giuliodori et al., 2010). CspA is known to be an RNA chaperone that accumulates during growth at low temperatures, and it affects both the transcription and translation of target genes that promote low temperature survival of cells (Jiang et al., 1997). Meanwhile, in our study, overexpressing CspA could promote growth at optimal and elevated temperatures as well (Fig 2 A to C). Clearly, *B. coagulans* employs a cold shock protein to aid in high-temperature growth, and evidently this CspL protein can promote the high-temperature growth of a large diversity of microorganisms. Unlike CspA, which has a sophisticated 5’-UTR thermal sensitive region (Giuliodori et al., 2010), CspL does not have a similar UTR region in *B. coagulans* 2-6. Additionally, we heterologously expressed CspL without the 5’-UTR, and the RNA binding function still worked in other organisms. The RNA decay showed decay rate of some CspL binding targets were decreased when *cspA* expressed *in vivo* as well. It might be indicated that even both CspA and CspL could binding mRNA fragments, but different preferential targeting (Figure 6).

Previous studies reported that ơ^E^ is essential for viability (De Las Penas et al., 1997), and ơ^E^ is required to maintain the integrity of the cell envelope (Hayden and Ades 2008; Nicoloff et al., 2017). Based on our proteome data, the amount of ơ^E^ in the *cspL*-expressing strain only had 0.05-fold change compared to the empty-vector cells, indicating that some unknown regulatory pathways were activated to replace the role of ơ^E^ in heat shock response when CspL is present (Figure 8). Analysis of functional pathways and biological processes enriched among RNAs detected by RIP-seq suggested that CspL binds to many genes that are to ribosome synthesis and RNA degradation. This strongly indicated that the amount of ribosomes increased and RNA decay was prolonged when CspL exists. That CspL promotes growth and changes cellular morphology at both 37 °C and 45 °C demonstrated that those changes did not depend on temperature induce. In most of our experiments, CspL acts as a global regulator in the stress response. The *cspL*-expressing cells could be to respond to stress immediately and make cells adapt to new circumstances. The CspL functional application in industrial microbes proved to be a successful practice at promoting ansamitocin P-3 and L-lactate acid productivities.

**Figure 8.**
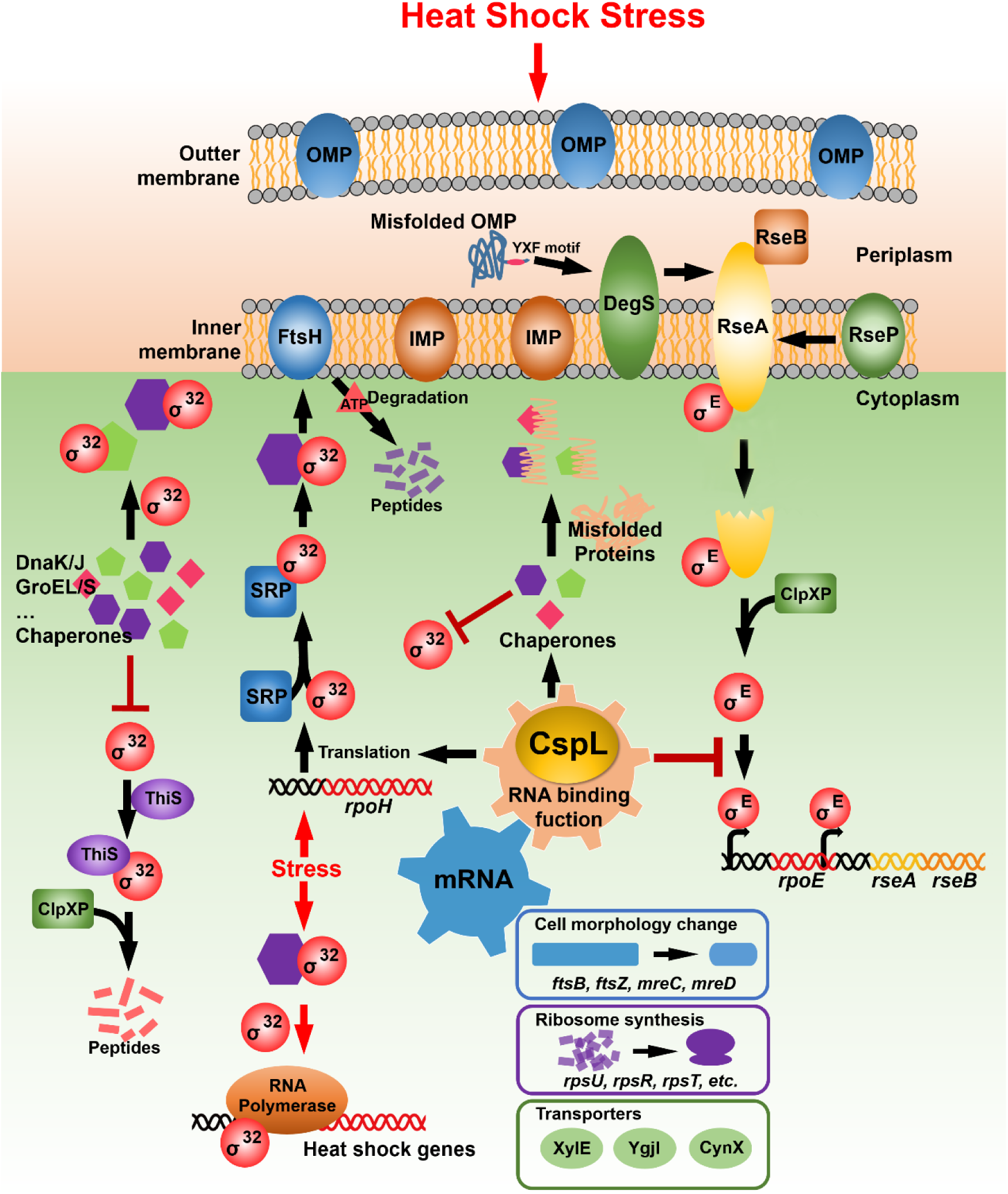
Regulation of the heat shock response and proposed regulatory controls of CspL in *E. coli*. The alternative sigma factors ơ^32^ and ơ^E^ are the most important heat shock regulators in *E. coli*. Both factors are cytoplasmic proteins that exist even at 37 °C, but they are regulated via several mechanisms. The sigma factor ơ^32^ is activated by denatured proteins that appear in the cytoplasm during heat insult. At the physiologically optimum conditions, a part of free ơ^32^ is sequestered either by the DnaK or GroEL chaperone system and forms the chaperone-ơ^32^ complex. Additionally, another part of free ơ^32^ molecules is bound either by the signal recognition particle (SRP) and guided to the FtsH protease, or modified by the ubiquitin-like protein ThiS and guided to the ClpXP protease for degradation. These regulatory circuits could maintain the ơ^32^ factor at a basal level. After heat insult, the chaperones dissociate from the complex and bind to the denatured proteins and fold proteins in the right way. This mechanism is called chaperone titration. Expression of CspL could promote an increased translation level of ơ^32^ that affects many biological processing *in vivo*. On the other hand, the sigma factor ơ^E^ different from ơ^32^ is activated by unfolded proteins in the periplasm under heat stress. The antisigma factor RseA is inserted into the inner membrane by one transmembrane domain. The co-antisigma factor RseB binds to part of the C-terminal exposed in the periplasm, while the N-terminal end extending into the cytoplasm sequesters ơ^E^ at 37°C. When growth temperature upshifts, the C-terminal ends of partially unfolded outer membrane proteins activate the DegS protease, which will cleave within the periplasmic domain of RseA. This leads to a conformational change within its inner membrane domain, activating the RseP protease. The protease cleaves within this domain, thereby releasing the remaining part of RseA with ơ^E^ still bound to it into the cytoplasm. It will be completely degraded by ClpXP or other protease to finally release ơ^E^. Here, the ơ^E^ translation level was significantly repressed under dual elements, heat stress, and CspL presence. When one of elements is removed, the ơ^E^ translation level did not show notable changes.

Moving forward, research will likely focus on the implementation and optimization of CspL expression as a tool to promote high-temperature growth, as well as delineating the range of microorganisms that can receive growth-promoting benefits from the transgenic expression of CspL. The large number of targets identified by our RIP-seq analysis make it clear CspL functions as a global RNA chaperone that somehow promotes high temperature growth. It will be interesting to determine the extent to which CspL targets particular subsets of transcripts based on some sequence motifs, or other higher-order structural features. Our findings highlight the fact that transgenic cspL expression improved *E. coli* growth at its normal 37 °C culture temperature, which underscores the importance of identifying the particular positive contribution that RNA binding by CspL makes to bacterial growth. The functions of CspL coincide with the core idea of current synthetic biology. How to neatly manipulate and modify the core metabolic network or basic functional cellular processing to achieve the goal by an easier way has long been desired to implement in the potential industrial bacteria. Design from the top level in synthetic biology could be have many advantages comparing with normal metabolic engineering. Likewise, when CspL exists, the transcriptional and post-transcriptional level regulation may make alters in the natural robustness of metabolic flux distribution in the industrial microbes, while not increasing the metabolic burden for cells.

## Materials and methods

### Bacterial strains and primers

Detailed information about the bacterial and yeast strains, as well as the plasmids and oligonucleotide primers used in this study are listed in S7, S8 and, S9 Table, respectively.

### Bacterial growth conditions and quantification of cell density

To evaluate the growth of Bacillus coagulans 2-6 under different temperatures, cells were grown overnight in 100 ml of GSY medium (20 g/l glucose, 10 g/l yeast extract, 5 g/l tryptone, 5 g/l CaCO3) without antibiotics in 250 ml Erlenmeyer flasks at 37 °C and 60 °C on a rotary shaker (200 r.p.m.). E. coli and Pseudomonas putida were grown in Luria-Bertani (LB) medium with appropriate antibiotics at 37 °C, and Saccharomyces cerevisiae was grown in yeast extract peptone dextrose medium (YEPD medium: 10 g/l yeast extract, 10 g/l peptone, 20 g/l glucose) with appropriate antibiotics at 37 °C. Cell densities were monitored by measuring optical density at 600 nm using a MAPADA V-1200 spectrophotometer. The overnight cultures were used to seed fresh medium (OD600 of 0.1 at the time of transfer). Growth at 37 °C and 60 °C was monitored every 2 hours throughout the incubation period. For dry cell weight measurements, a 30 ml cell suspension was centrifuged at 12,000 g for 8 min in pre-weighed microcentrifuge tubes. The cell pellets were washed twice in water and dried at 50 °C until the mass of each tube remained constant over time (typically after ∼48 h).

### Scanning electron microscopy

For scanning electron microscopy, cell cultures were grown in the appropriate medium with antibiotics. Cells were collected by centrifugation and washed three times with 0.1 M PBS buffer (pH 7.4). The cells were then fixed for 1 h at room temperature in 0.1 M PBS buffer containing 2.5% glutaraldehyde. Samples were dehydrated through a graded ethanol series up to absolute ethanol, critical point dried with liquid CO2, mounted on aluminium stubs and sputter coated with gold-palladium. Imaging was carried out on a Hitachi S-4800 scanning electron microscope.

### Preparation of proteomes, 2D-LC/MS analysis, and protein identification

Cells of *B. coagulans* 2-6 grown in GSY medium were harvested by centrifugation, washed twice in PBS buffer, and the cell pellets were lysed by resuspending in buffer (8 M urea, 0.05% SDS, 10 mM DTT, 10 mM Tris, pH 8.0) and grinding under liquid nitrogen. After centrifugation at 12,000 g (10 min, 4 °C), the supernatant was mixed with precooled acetone at a volume-to-volume ratio of 1:4. Following overnight incubation at -20 °C, the mixture was centrifuged at 12,000 g (10 min, 4 °C). The pellets were washed with precooled acetone three times, and resuspended in buffer containing 6 M Gu-HCl and 100 mM Tris, pH 8.3. The protein content was measured using a modified Bradford protocol. Total protein (100 μg) was resuspended in buffer containing 10 mM DTT and incubated at 56 °C for 0.5 h. Then 50 mM IAA was added, and the sample was incubated at 25 °C for 40 min. After 3 K ultrafiltration, membrane ultrafiltration, and flushing of the membrane with 100 mM NH4HCO3, the pH of the solution was adjusted to 8.0-8.5. Next, 40 μg of sequencing-grade modified trypsin was added to the extract and digestion was carried out overnight at 37 °C with gentle rotation (protein : trypsin ratio = 50 : 1).

In order to separate and analyze the peptides in the samples, we used multi-dimensional liquid chromatography with an Agilent 1100 LC system. Separation in the first dimension began with the elution of peptides from a strong cation exchange silica column (0.075 mm × 5 cm). Next, a C18 column (0.075 mm × 10 cm) (Column Technology Inc.) was used with a continuous linear salt gradient (0 - 130 min, 2% - 35%; 130 - 135 min, 35% - 90%; 135 - 140 min, 90%; 140 - 141 min, 90% - 2%; 141 - 180 min, 2%); Chromatography conditions: buffer A: H2O; buffer B: acetonitrile. Finally, a nanospray column was directly interfaced to the orifice of an LTQ Classic ion trap mass spectrometer (ThermoFisher). Nanospray ionization was accomplished with a spray voltage of 3.5 kV and capillary temperature of 200 °C. The m/z scan range was from 400 to 1800.

Database searches for MS and MS/MS spectra were conducted using proteomics discovery software V1.2 (ThermoFisher, CA, USA). Mass spectra were analyzed using Bioworks software. We generated a predicted protein database from the annotated *B. coagulans* 2-6 genome. The peptide matches with an assumed charge state of z = 1 and an XCorr score of > 2.2, or charge state of z = 3 and an XCorr score of > 3.75 were automatically accepted as valid. High scoring peptide matches were automatically identified and retained.

### RNA deep-sequencing and identification of differentially expressed mRNA

Total RNA was extracted from *B. coagulans* 2-6 and *E. coli* DH5α with the RNAiso Plus kit (Takara, Japan). RNA deep-sequencing was performed by a commercial sequencing company (Novogene, China). Briefly, sequencing libraries were generated from rRNA-depleted RNA using the NEBNext^®^ Ultra™ Directional RNA Library Prep Kit for Illumina^®^ (NEB, USA) following the manufacturer’s recommendations. The libraries were sequenced on the Illumina Hiseq 2000 platform, and 100 bp paired-end reads were generated. Demultiplexed and quality filtered reads were then aligned to *B. coagulans* 2-6 and *E. coli* DH5α reference genome sequence using TopHat (V.2.0.8). The mapped reads from each sample were assembled using Trinity with a reference-based approach.

Cuffdiff (v2.1.1) was used to calculate the RPKM (Reads Per Kilobase per Million mapped reads) of coding genes in each sample. Gene RPKM values were computed by summing the FPKM values of transcripts in each gene group. Cuffdiff provides a statistical method for determining differential expression of digital transcripts or gene expression data using a model based on the negative binomial distribution. Genes with a *P* value < 0.05, FKPRM > 0.5 and FC > 2 were classified as differentially expressed.

### Expressing B. coagulans 2-6 genes in E. coli, S. cerevisiae, and P. putida

PCR amplification was carried using the Phanta Super-Fidelity DNA Polymerase (Vazyme, China) according to the manufacturer’s instructions. The sequences of all the plasmids produced were verified by restriction mapping and/or DNA sequencing. We cloned 38 genes from *B. coagulans* 2-6 (*SI Appendix*, Table S5) into the pUC19 vector and then transformed these constructs into *E. coli* DH5α competent cells. Positive transformants were selected on LB medium plates containing ampicillin (50 mg/l) and were confirmed via PCR. We also cloned the *BCO26_cspL* gene into the pYES2 and pME6032 vectors and then transformed these constructs into *S. cerevisiae* and *P. putida* competent cells, respectively. Positive *S. cerevisiae* and *P. putida* transformants were selected on YPD medium containing 50 mg/l ampicillin and LB medium containing 25 mg/l tetracycline, respectively, and were confirmed via PCR.

The growth of the transformants at 37 °C and 45 °C was monitored via absorbance measurements using a Bioscreen C^®^ analyzer (Labsystems, Finland). All transformants were grown overnight at 42 °C and then inoculated into fresh media in Honeycomb plates (10 x 10 wells) containing the appropriate antibiotics. Quintuplicates of each engineered strain were aliquoted into 200 μl wells. The plates were shaken continuously; readings were taken at a wavelength of 600 nm.

### Bio-Layer Interferometry (BLI)

DNA-protein binding kinetics were measured using the Octet RED96 System (ForteBio). In the preparation stage, fixed concentrations (1 µM) of 5’biotin-TEG-labeled duplex oligonucleotide probes representing the A or G alleles of rs7279549 (sequences are presented in *SI Appendix*, Table S10) were immobilized on streptavidin-modified sensor surfaces. In the first phase, nuclear protein extracts were allowed to interact with the DNA immobilized on the sensor surface in dilution buffer for 300 s. In the second phase, PBS/Tween buffers were used to elute the DNA-protein compound from the sensor surface; this phase was sustained for 300 s. BLI experiments were repeated three times.

### RIP-seq experiments

Cells were cultured overnight in LB medium with ampicillin. After centrifugation the supernatant was discarded, and the pellet was washed twice with 5 ml of ice-cold PBS and lysed with 600 μl of lysis buffer (20 mM Tris-HCl pH 7.4, 150 mM NaCl, 5 mM MgCl2, and 1 mM DTT) containing 1% Triton X-100 and Turbo DNase I (Invitrogen) 25 U/ml. The sample was then clarified by centrifugation for 10 min at 20,000 g, 4 °C. The supernatant was incubated at 4 °C for 30 min with 60 μl of Dynabeads M-270 Streptavidin (Invitrogen) equilibrated with lysis buffer containing 1% Triton X-100. The beads were washed three times with lysis buffer containing 1% Triton X-100 and 1 M NaCl. His-BCO26-CspL and bound RNAs were eluted with 25 μl of lysis buffer containing 5 mM biotin at 4 °C for 30 min. RNAs were extracted with QIAzol (Qiagen) using the Direct-zol RNA miniprep (Zymo Research). Sequencing libraries were prepared using the TruSeq Stranded Total RNA Library Prep Kit with Ribo-Zero Gold (Illumina). Libraries were sequenced on the HiSeq2000/2500 (Illumina) platform.

### RNA decay analysis

*E. coli* overnight cultures were diluted 1:150 into 50 ml fresh LB media with proper antibiotic and incubated at 37 °C and 45 °C until reaching an optical density (OD600) of 0.5. The 250 μl Rifampicin (100 mg/ml, for a final concentration of 500 μg/ml) were immediately added to the culture to inhibit RNA synthesis. Selected time points were sampled by collecting 2 ml from the culture into a pre-chilled tube containing 220 μl of ice-cold stop solution (90% ethanol and 10% saturated phenol) to deactivate cellular processes and RNA-decay. The sample was quickly vortexed and then placed on ice until all time points were collected. RNA extraction was performed by RNAprep pure Cell / Bacteria Kit (Tiangen, DP430). The cDNA synthesis was performed by FastKing RT Kit (Tiangen, KR116), then the cDNA products used as templates for RT-qPCR. The RT-qPCR was performed by Talent RT-qRCR Kit (SYBR Green) (Tiangen, FP209). The CT value of 16s rRNA was used as reference, testing genes expression CT values were subtracted reference to eliminate inevitable errors.

### Validating the function of CspL in *E. coli* eGFP expressing system, *Actinosynnema pretiosum*, and *Bacillus licheniformis* fermentation

For validating the function of CspL, *E. coli* eGFP expressing system (strain *E. coli* eGFP) was constructed. The gene sequence of eGFP (Pfam: 01353) was synthesized by Sangon (Shanghai) and transferred into *E. coli* DH5α competent cell by harboring plasmid pET28a. Using LB plate and PCR to confirm positive transformants, the plasmid pUC19 harboring *cspL* was then transferred into those competent positive tranformants, and confirmed via PCR. The control group used empty pUC19 vector in place of pUC19-*cspL*. Wild type, control, and *E. coli* eGFP-pUC19-*cspL* were grown overnight in LB medium at 45 °C. The overnight culture was inoculated 1 : 100 into fresh LB medium with appropriate antibiotic (20 μg/ml kanamycin and 100 μg/ml ampicilin) to create a culture stock. The 250 ml Erlenmeyer flasks contained 50 ml of culture stock and cultured in the shaker at 45 °C. Every two hours, the cultures were checked for cell density and fluorescence signal. The method of cell density was mentioned previously. An aliquot of the fresh culture stock (200 μl) was transferred into polystyrene 96-well Costar Assay Plate (black with clear flat bottom, Corning Inc., New York). The plate was shaken in linear mode for 30 s and green fluorescence (λex = 485 nm, λem = 535 nm) was monitored using an Infinite F200 multimode reader (TECAN, San Jose, CA).

For evaluating the function of CspL in high value-added industrial microbes, *A.pretiosum* ATCC31280 (used as AP-3 producing strain) [31] and its derivatives were cultured at 30 °C on YMG agar (0.4% yeast extract, 1.0% malt extract, 0.4% glucose, 2.0% agar (w/v), pH 7.2-7.3). For metabolites analysis, the first seed medium (3.0% tryptone soya broth powder, 0.5% yeast extract and 5.0% sucrose (w/v), pH 7.5) was inoculated with agar-grown mycelia and cultivated at 30 °C, 220rpm for 24h. Subsequently, the second medium (3.0% tryptone soya broth powder, 0.5% yeast extract and 2.5% sucrose, 1.0% soluble starch (w/v), 0.05% isobutanol and 0.05% isopropanol (v/v), pH 7.5) was inoculated with 1 ml of the first seed culture, inoculated for another 24 h at 30 °C. Fermentation medium (yeast extract 0.8%, malt extract 1.0%, sucrose 1.5%, soluble starch 2.5% (w/v), isobutanol 0.5%, isopropanol 1.2% (v/v), pH 7.5) was inoculated with the second seed culture at 10% (v/v) and inoculated at 25 °C and 220 r.p.m. for 7 days. *E. coli* DH10B and *E. coli* ET12567/pUZ8002 were used for plasmid construction and intergeneric conjugation, respectively. For overexpression of the gene *cspL*, the plasmid pLQ856 with the kasOp* promoter cloned into BamHI/SpeI-digested plasmid pDR3 was used. The sequenced *cspL* encoding gene was inserted into plasmid pLQ856 under the control of kasOp* promoter, generating plasmid pLQ856-*cspL*. The recombinant plasmid was introduced into ATCC31280 from *E. coli* ET12567/pUZ8002 through intergeneric conjugation. Additionally, the plasmid pLQ856 was introduced into ATCC31280 generating a control strain ATCC31280::pLQ856. To quantify AP-3 production, the supernatant of the fermentation broth was extracted with 2 volume of ethyl acetate and evaporated. The residues were dissolved in methanol, passed through 0.22-μm filters, and applied to HPLC. The HPLC analysis was operated on Agilent series 1260 (Agilent Technology, USA) with an Agilent Eclipse Plus-C18 column (4.6×150 mm, 5 μm). AP-3 analyzed at a flow rate of 0.5 ml/min, with the following gradient: 0-5 min 10%-50% B, 5-10 min 50%-60% B, 10-20 min 60%-75% B, 20-30 min 75%-95% B, 30-38 min 95% B, 38-39 min 95%-10% B, 39-48 min 10% B (solvent A: water, solvent B: methanol), and detected at 236nm and 254nm.

For validating the function of CspL in industrial microbes, the Gram-positive bacteria *Bacillus licheniformis* BN11 was used, which is an efficient genetically modified D-lactate producer. BN11 were statically cultivated in GSY medium at 50°C for 24 h. The D-lactate fermentation medium is composed of 100 g/l glucose and 40 g/l peanut meal. Neutral protease (0.3 g/l was added at the beginning of fermentation. The synthetic DNA fragment with the inducible promoter Pgrac100 and the codon-optimized gene *cspL* was digested with EcoRI and BamHI, and ligated into the *E. coli*-*B. subtilis* shuttle vector pEB03 to generate the plasmid pEB03-Pgrac100*cspL*. The plasmid pEB03-Pgrac100*cspL* and pEB03 were transformed into *E. coli* S17-1 to generate the donors in conjugation with *B. licheniformis* BN11. The conjugation and transformation experiments were performed as described previously. Spectinomycin (100 mg/l) and polymyxin B (40 mg/l) was added for selection of recombinant *B. licheniformis* BN11 harboring the shuttle plasmid pEB03 or pEB03-Pgrac100*cspL*. The D-lactate fed-batch fermentation experiments were carried out in a 5 L-bioreactor containing 2.7 liter of fresh medium. The fermentation temperature was controlled at 50 °C and the inoculum volume was 10% (v/v). The pH was maintained at 7.0 by the automated addition of 25% (w/v) Ca(OH)2. The glucose concentration was maintained between 20 g/l and 120 g/l by adding glucose powder. After 10 h of fermentation, 0.1 mM IPTG was added. Glucose concentration was estimated by a SBA-40D biosensor analyzer. The fermentation samples were heated to 100 °C for 10 min and then acidified with the addition of an equal volume of 2 M H2SO4 for 10 min. The acidized samples were diluted 50-fold and centrifuged at 8,000 r.p.m. for 10 min. Filtered supernatant can be used for subsequent detection. Lactate concentration was determined by high-performance liquid chromatography (HPLC) using a Bio-Rad Aminex HPX-87H column.

## Acknowledgements

This study was supported by grants from the Chinese National Science Foundation for Excellent Young Scholars (31422004), by grants from the Science and Technology Commission of Shanghai Municipality (17JC1403300), by the “Shuguang Program” (17SG09) which is supported by the Shanghai Education Development Foundation and Shanghai Municipal Education Commission, and by grants from the Chinese National Science Foundation (31870088). We also thank John Hugh Snyder for editing the manuscript.

## Author contributions

Z.Z., H.T. and P.X. conceived the project and wrote the manuscript. Z.Z., H.T., W.W. and L. Z. designed and performed all the experiments. Z.Z., H.T., F.S., and F.T. analyzed the results.

## Competing financial interests

The authors declare no competing financial interests.

## Supplementary legends

**Figure 1–figure supplement 1.**
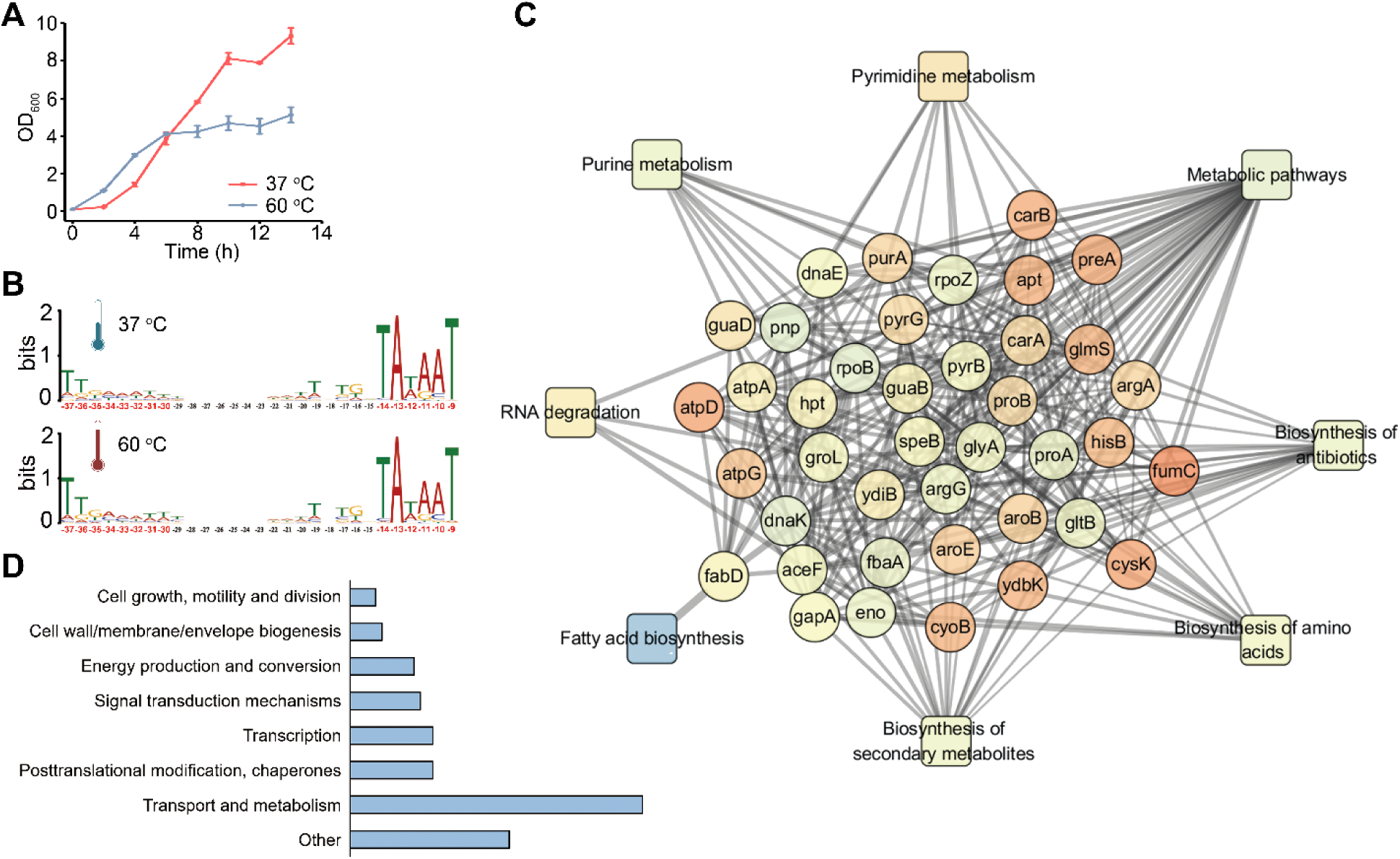
Transcriptome and proteome analysis of *B. coagulans* 2-6. (A) The growth curves of *B. coagulans* 2-6 at 37 °C and 60 °C conditions. (B) Sequence elements of promoters under different conditions. Culturing at 37 °C and 60 °C, the transcriptional start sites, operon structure and the length of 5’ UTR does not shown significant difference. The regions -14 to -9 and -37 to -30 are highly conservation under different conditions. (C) Protein-protein interaction network of *B. coagulans* 2-6. Circle node color indicates genetic neighborhood connectivity. Light blue, low; light yellow, medium; light orange, high. Round rectangle nodes color indicates different classification of GO analysis. Under different conditions of *B. coagulans* 2-6, a subset of highly connected protein nodes involved in key cellular processes undergoes temperature fluctuation. (D) GO Slim Mapper analysis was performed on the label-free proteome data set to identify the processes of up-regulated protein expressions at 60 °C.

**Figure 1–figure supplement 2.**
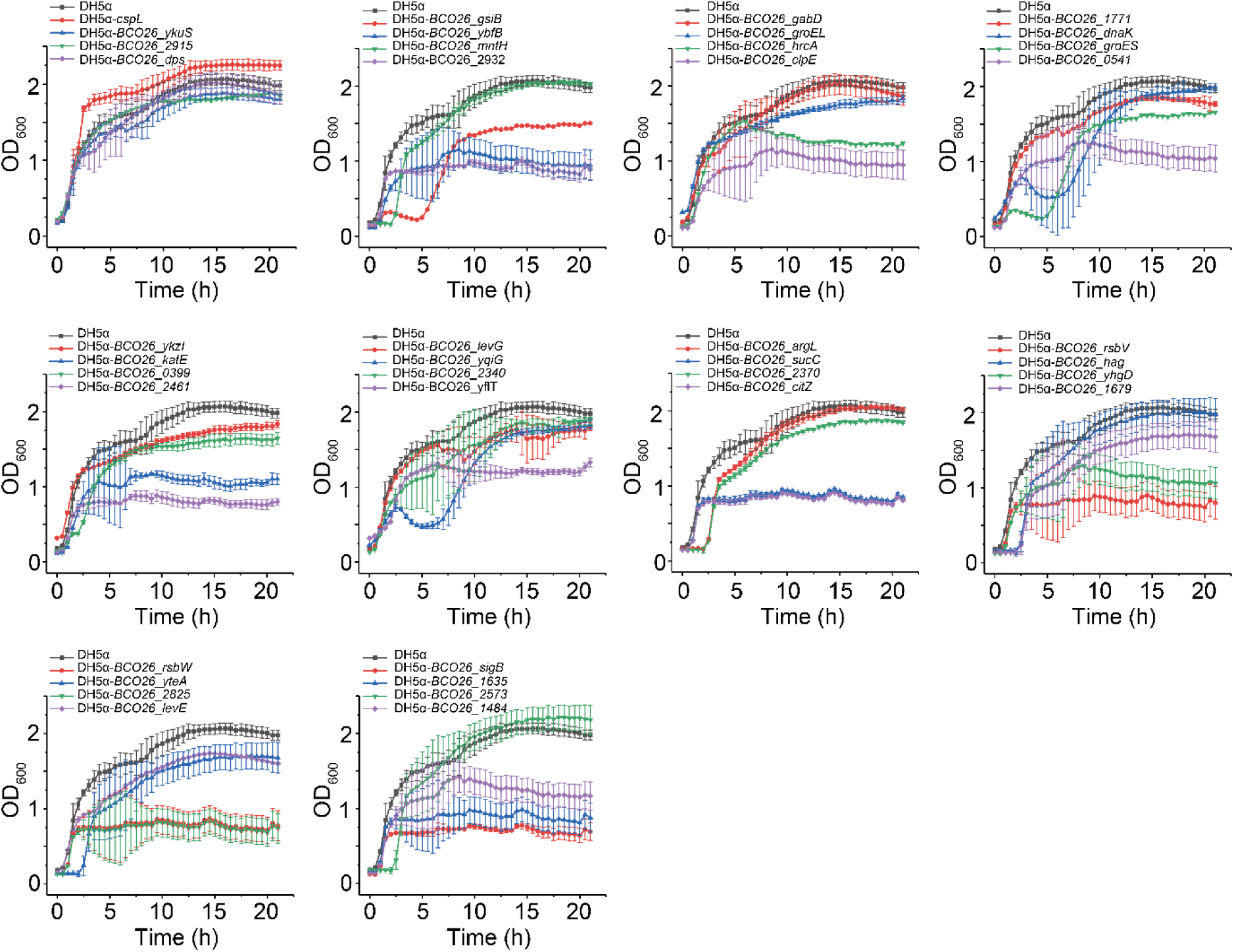
Growth curves of 38 candidate genes associated with high temperature growth.

**Figure 3–figure supplement 1.**
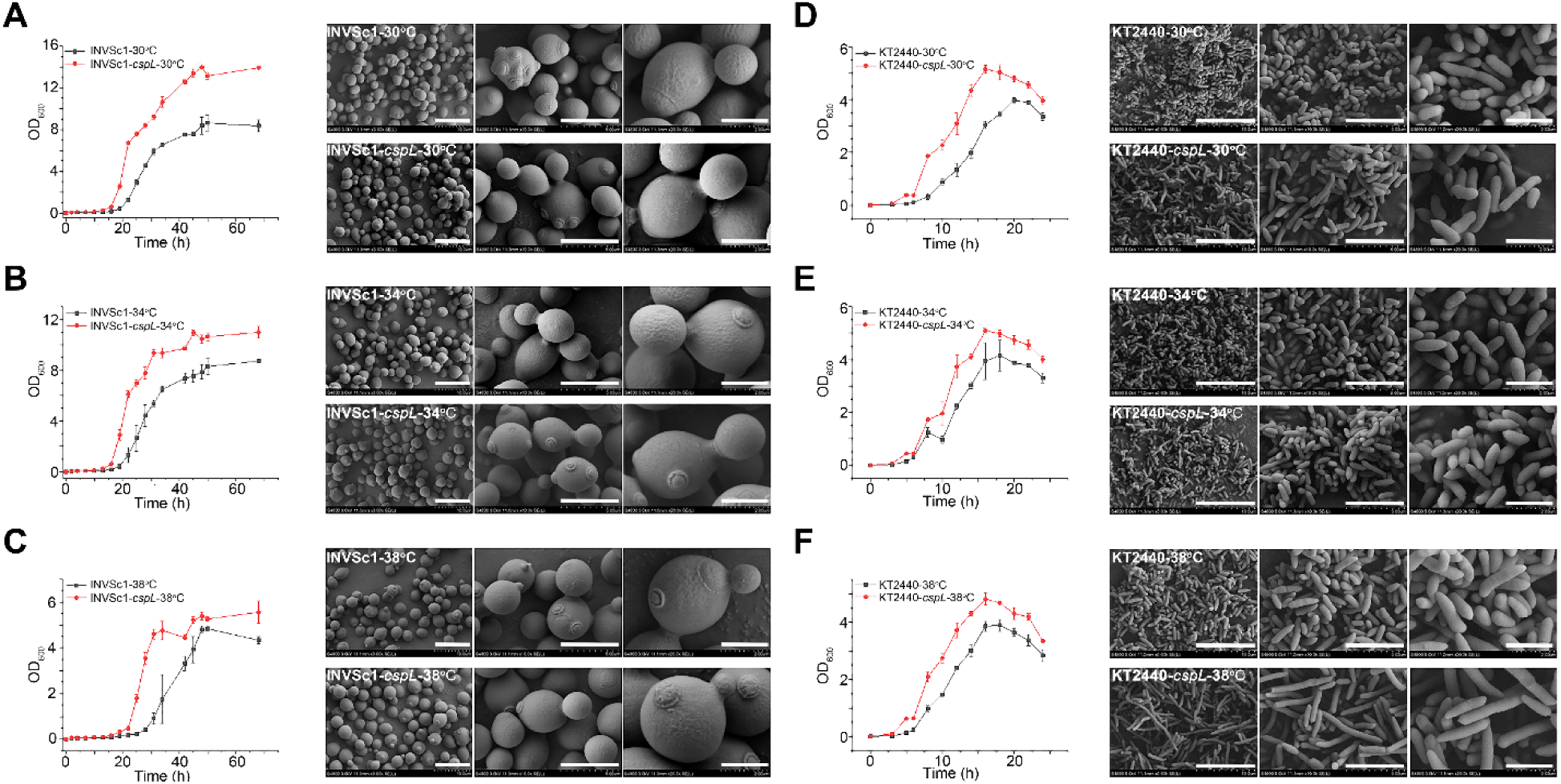
Growth curves and SEM images of *S. cerevisiae* and *P. putida* at three temperature gradient, namely, 30 °C, 34 °C, and 38 °C. In *S. cerevisiae* INVSc1, (A), (B), and (C), showed CspL expressing grows better than control at different temperature conditions. All three groups have no significant difference in cell morphology, except, the surface texture of CspL expressing strain showed obvious smoother than control. In *P. putida* KT2440, (D), (E), and (F), showed CspL expressing strain grows better than control at different temperature conditions. Two groups of 30 °C and 34 °C have no significant difference. In 38 °C group, the CspL expressing strain seemed slightly longer than control. (scare bar from left to right: 10 μm, 5 μm, and 2 μm)

**Figure 4-figure supplement 1.**
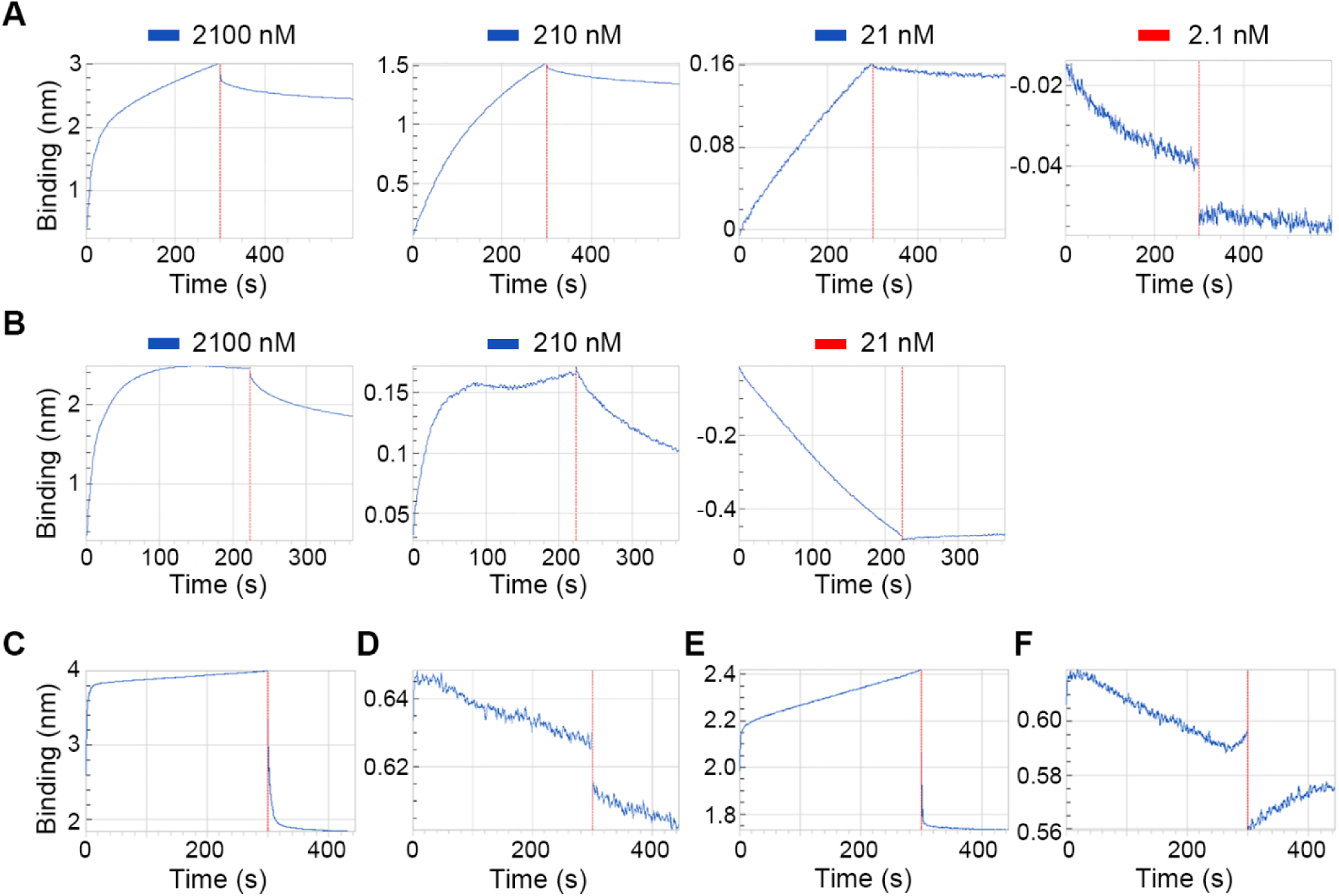
Concentration limitation of CspL binds to RNA and ssDNA in vitro. (A) Concentration limitation of CspL binds to 18 nt RNA fragment. The concentration gradient sets as 2100 nM, 210 nM, 21 nM, and 2.1 nM. When the concentration of CspL set as 2.1 nM, it lost the binding capacity. (B) Concentration limitation of CspL binds to 18 nt ssDNA fragment. The concentration gradient sets as 2100 nM, 210 nM, and 21 nM. When the concentration of CspL set as 21 nM, it lost the binding capacity. (C) Using ssDNA gaaC (biotin-CCGCAGAGAACGACGAGAGC) to bind to CspL, it showed positive binding signal. (D) Using complementary double strand of gaaC instead of ssDNA, it showed negative binding signal. (E) Using ssDNA random (biotin-CCGCAGATCCAGACGAGAGC) to bind to CspL, it showed positive binding signal. (F) Using complementary double strand of random gene instead of random ssDNA, it showed negative binding signal.

**Figure 5-figure supplement 1.**
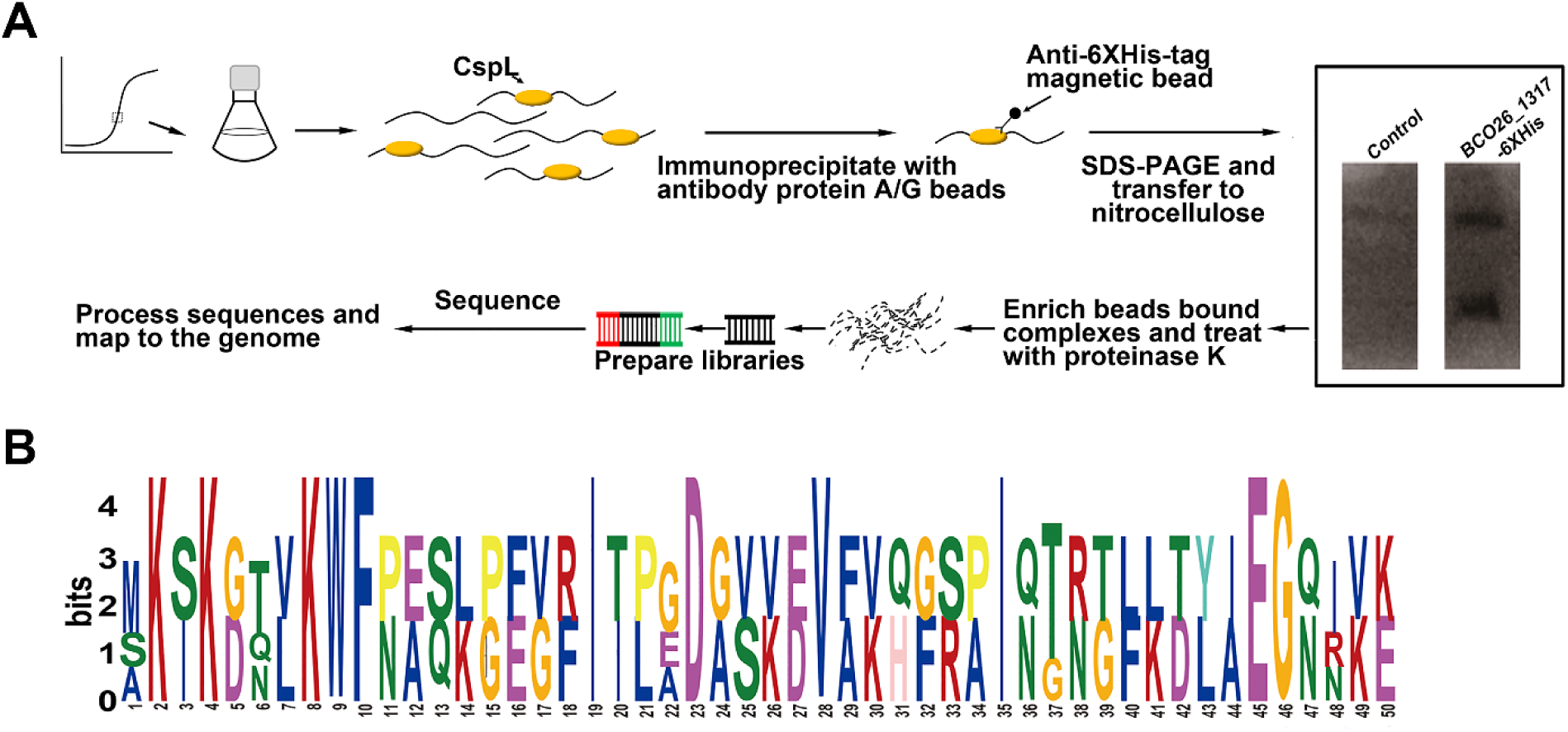
The work flow of RIP-seq and analysis. (A) RIP-seq processing. (B) Putative motif analysis of CspL’s binding targets.

