## Supplemental Tables for "A cold shock protein from a thermophile bacterium promotes the high-temperature growth of bacteria and fungi through binding to diverse RNA species"

**Table S1. Up-regulated genes in *B. coagulans* 2-6 by RNA-seq**

| Locus | Gene | $Log_2FC$[*] | Q-value[**] | Product |
|---|---|---|---|---|
| BCO26_0715 | *uvr*B | 1.3477 | 0.0488 | excinuclease ABC subunit B |
| BCO26_1892 | *clp*X | 1.4211 | 0.0479 | ATP-dependent Clp protease, ATP-binding subunit ClpX |
| BCO26_2971 | *trm*E | 1.4927 | 0.0289 | tRNA modification GTPase TrmE |
| BCO26_2177 | *yut*G | 1.5406 | 0.0377 | phosphatidylglycerophosphatase A |
| BCO26_1589 | *res*D | 1.5476 | 0.0199 | winged helix family two component transcriptional regulator |
| BCO26_0096 | *glt*X | 1.5499 | 0.0111 | glutamyl-tRNA synthetase |
| BCO26_0306 | *yqi*G | 1.6094 | 0.0296 | NADH:flavin oxidoreductase/NADH oxidase |
| BCO26_2190 | *suf*S | 1.6196 | 0.0230 | cysteine desulfurase |
| BCO26_0964 | *pdh*C | 1.6202 | 0.0363 | hypothetical protein |
| BCO26_2951 | *rpl*I | 1.6710 | 0.0290 | 50S ribosomal protein L9 |
| BCO26_2197 | *yus*I | 1.7026 | 0.0200 | arsenate reductase and-like protein |
| BCO26_1748 | *hrc*A | 1.7120 | 0.0134 | heat-inducible transcription repressor HrcA |
| BCO26_2975 | *rpm*H | 1.7207 | 0.0492 | 50S ribosomal protein L34 |
| BCO26_0063 | *fol*B | 1.7284 | 0.0340 | dihydroneopterin aldolase |
| BCO26_1185 | *ylx*Y | 1.7293 | 0.0117 | sporulation protein, polysaccharide deacetylase family |

| | | | | |
|---|---|---|---|---|
| BCO26_1352 | *cit*B | 1.7431 | 0.0276 | aconitate hydratase 1 |
| BCO26_2932 | - | 1.7449 | 0.0194 | malate/quinone oxidoreductase |
| BCO26_0951 | *yku*F | 1.7638 | 0.0302 | short-chain dehydrogenase/reductase SDR |
| BCO26_0090 | *mcs*B | 1.7676 | 0.0036 | ATP:guanido phosphotransferase |
| BCO26_0833 | *mec*A | 1.7769 | 0.0045 | Negative regulator of genetic competence |
| BCO26_1746 | *dna*K | 1.7928 | 0.0036 | chaperone protein DnaK |
| BCO26_0837 | *yjb*I | 1.7956 | 0.0116 | globin |
| BCO26_1704 | *pbp*A | 1.8446 | 0.0015 | penicillin-binding protein transpeptidase |
| BCO26_2943 | *yyc*J | 1.8537 | 0.0272 | beta-lactamase domain-containing protein |
| BCO26_2952 | *yyb*T | 1.8883 | 0.0019 | diguanylate cyclase and phosphoesterase |
| BCO26_0091 | *clp*C | 1.8921 | 0.0019 | ATPase AAA-2 domain-containing protein |
| BCO26_2331 | *glt*D | 1.9013 | 0.0189 | glutamate synthase, NADH/NADPH, small subunit |
| BCO26_2036 | *ytp*P | 1.9092 | 0.0200 | Thioredoxin domain-containing protein |
| BCO26_0601 | - | 1.9444 | 0.0340 | glycoside hydrolase clan GH-D |
| BCO26_0973 | *suh*B | 1.9543 | 0.0036 | inositol monophosphatase |
| BCO26_0963 | *pdh*B | 1.9953 | 0.0049 | transketolase central region |
| BCO26_0714 | - | 2.0601 | 0.0007 | hypothetical protein |

| | | | | |
|---|---|---|---|---|
| BCO26_1117 | *suc*C | 2.0623 | 0.0075 | succinyl-CoA synthetase subunit beta |
| BCO26_2480 | *ywa*C | 2.0796 | 0.0004 | RelA/SpoT domain-containing protein |
| BCO26_0753 | *yha*R | 2.1031 | 0.0021 | enoyl-CoA hydratase/isomerase |
| BCO26_2608 | *kip*A | 2.1125 | 0.0087 | urea amidolyase-like protein |
| BCO26_1276 | *odh*B | 2.1401 | 0.0019 | 2-oxoglutarate dehydrogenase, E2 subunit, dihydrolipoamide succinyltransferase |
| BCO26_2370 | - | 2.1486 | 0.0070 | anti-sigma-factor antagonist |
| BCO26_2499 | *mtb*P | 2.1487 | 0.0436 | Modification methylase |
| BCO26_0089 | *mcs*A | 2.1513 | 0.0467 | UvrB/UvrC protein |
| BCO26_1679 | - | 2.1551 | 0.0104 | ribonucleoside-diphosphate reductase, adenosylcobalamin-dependent |
| BCO26_2249 | - | 2.1588 | 0.0054 | hypothetical protein |
| BCO26_2942 | *htr*C | 2.2230 | 0.0073 | peptidase S1 and S6 chymotrypsin/Hap |
| BCO26_1769 | *ysf*B | 2.2363 | 0.0246 | transcriptional regulator CdaR |
| BCO26_2562 | *bdh*A | 2.2874 | 0.0001 | alcohol dehydrogenase zinc-binding domain-containing protein |
| BCO26_0542 | *lev*F | 2.2885 | 0.0036 | phosphotransferase system PTS sorbose-specific IIC subunit |
| BCO26_2541 | - | 2.2965 | 0.0467 | SMC domain-containing protein |
| BCO26_2741 | *sig*B | 2.3006 | 0.0001 | Sig B/F/G subfamily RNA polymerase sigma-28 subunit |
| BCO26_0241 | *gab*D | 2.3475 | 0.0002 | succinic semialdehyde dehydrogenase |

| | | | | |
|---|---|---|---|---|
| BCO26_0171 | *arg*I | 2.3475 | 0.0271 | arginase |
| BCO26_1428 | *exo*A<br>A | 2.3652 | 0.0105 | exodeoxyribonuclease III |
| BCO26_1353 | - | 2.3995 | 0.0194 | hypothetical protein |
| BCO26_2824 | *yhc*G | 2.4143 | 0.0351 | ABC transporter-like protein |
| BCO26_1771 | - | 2.4602 | 0.0019 | heat shock protein Hsp20 |
| BCO26_1419 | *rtp* | 2.4726 | 0.0436 | replication terminator protein |
| BCO26_2570 | *rbs*K | 2.4936 | 0.0237 | ribokinase |
| BCO26_0895 | *yue*B | 2.4990 | 0.0116 | hypothetical protein |
| BCO26_0543 | *lev*G | 2.5090 | 0.0049 | PTS system, mannose/fructose/sorbose family, IID subunit |
| BCO26_0541 | - | 2.5409 | 0.0001 | PTS system, mannose/fructose/sorbose family, IIA subunit |
| BCO26_0590 | - | 2.6812 | 0.0481 | hypothetical protein |
| BCO26_0591 | - | 2.6841 | 0.0036 | NADPH-dependent FMN reductase |
| BCO26_1635 | *yqi*W | 2.6885 | 0.0077 | hypothetical protein |
| BCO26_2944 | *yyc*I | 2.7032 | 0.0000 | YycI protein |
| BCO26_3000 | | 2.7343 | 0.0194 | putative transposase |
| BCO26_0474 | *yvf*I | 2.7422 | 0.0018 | GntR domain-containing protein |

| | | | | |
|---|---|---|---|---|
| BCO26_2742 | *rsb*W | 2.7626 | 0.0000 | putative anti-sigma regulatory factor, serine/threonine protein kinase |
| BCO26_2743 | *rsb*V | 2.8194 | 0.0000 | anti-sigma-factor antagonist |
| BCO26_2424 | *yte*A | 2.8282 | 0.0001 | TraR/DksA family transcriptional regulator |
| BCO26_0740 | - | 2.8361 | 0.0000 | hypothetical protein |
| BCO26_2980 | - | 2.9117 | 0.0000 | hypothetical protein BcoaDRAFT_4102 |
| BCO26_0934 | *clp*E | 2.9677 | 0.0234 | ATPase AAA-2 domain-containing protein |
| BCO26_2825 | - | 2.9723 | 0.0001 | GntR family transcriptional regulator |
| BCO26_0737 | - | 2.9815 | 0.0005 | hypothetical protein |
| BCO26_0739 | - | 3.0232 | 0.0000 | hypothetical protein |
| BCO26_0540 | *lev*E | 3.1100 | 0.0001 | PTS system, mannose/fructose/sorbose family, IIB subunit |
| BCO26_2725 | *gro*ES | 3.2119 | 0.0000 | chaperonin Cpn10 |
| BCO26_2375 | *ybf*B | 3.2466 | 0.0000 | major facilitator superfamily protein |
| BCO26_2724 | *gro*EL | 3.3122 | 0.0000 | chaperonin GroEL |
| BCO26_0781 | *yhg*E | 3.3378 | 0.0000 | YhgE/Pip N-terminal domain-containing protein |
| BCO26_0400 | - | 3.3643 | 0.0000 | hypothetical protein |
| BCO26_2022 | *acs*A | 3.3911 | 0.0033 | AMP-dependent synthetase and ligase |
| BCO26_0738 | - | 3.3949 | 0.0001 | hypothetical protein |

| | | | | |
|---|---|---|---|---|
| BCO26_0780 | *yhg*D | 3.4007 | 0.0000 | TetR family transcriptional regulator |
| BCO26_0399 | - | 3.4057 | 0.0000 | hypothetical protein |
| BCO26_2638 | *gsi*B | 3.4856 | 0.0000 | general stress protein |
| BCO26_1964 | *cit*Z | 3.4935 | 0.0000 | 2-methylcitrate synthase/citrate synthase II |
| BCO26_0880 | *yku*S | 3.5367 | 0.0000 | hypothetical protein |
| BCO26_2535 | *kat*E | 3.5737 | 0.0132 | Catalase |
| BCO26_1318 | - | 3.6097 | 0.0000 | hypothetical protein |
| BCO26_1317 | *csp*C | 3.6187 | 0.0000 | cold-shock DNA-binding domain-containing protein |
| BCO26_2573 | - | 3.6433 | 0.0000 | hypothetical protein |
| BCO26_2080 | *dps* | 3.6944 | 0.0012 | Ferritin Dps family protein |
| BCO26_2371 | - | 3.7143 | 0.0000 | hypothetical protein |
| BCO26_0972 | *ykz*I | 3.8006 | 0.0000 | hypothetical protein |
| BCO26_1896 | - | 3.9051 | 0.0075 | hypothetical protein |
| BCO26_0370 | - | 3.9154 | 0.0000 | alcohol dehydrogenase GroES domain-containing protein |
| BCO26_2136 | - | 3.9586 | 0.0000 | short-chain dehydrogenase/reductase SDR |
| BCO26_2572 | - | 4.0593 | 0.0007 | hypothetical protein |
| BCO26_2079 | - | 4.0609 | 0.0004 | hypothetical protein |

| | | | | |
|---|---|---|---|---|
| BCO26_2925 | - | 4.1339 | 0.0000 | hypothetical protein |
| BCO26_2461 | - | 4.2752 | 0.0125 | hypothetical protein |
| BCO26_2926 | - | 4.3365 | 0.0000 | alanine racemase domain-containing protein |
| BCO26_2860 | *yfl*T | 4.4587 | 0.0000 | hypothetical protein |
| BCO26_1328 | *yxa*B | 4.5939 | 0.0033 | polysaccharide pyruvyl transferase |
| BCO26_2340 | - | 4.8911 | 0.0001 | hypothetical protein |
| BCO26_1065 | - | - | 0.0488 | transposase IS4 family protein |

*$Log_2FC$：the logarithm of fold change, **Q-value：adjustment of *P*-value

**Table S2. Down-regulated genes in *B.coagulans* 2-6 by RNA-seq**

| Locus | Gene | $log_2FC$* | Q-value** | Product |
|---|---|---|---|---|
| BCO26_1863 | *com*C | -9.4071 | 0.0036 | Prepilin peptidase |
| BCO26_0208 | *gnt*P | -8.2333 | 0.0196 | gluconate transporter |
| BCO26_1379 | *yxl*A | -6.6937 | 0.0202 | cytosine/purines uracil thiamine allantoin permease |
| BCO26_1506 | *pbu*X | -6.0585 | 0.0012 | xanthine permease |
| BCO26_2004 | *ycg*N | -5.3972 | 0.0000 | delta-1-pyrroline-5-carboxylate dehydrogenase |
| BCO26_0804 | *yxa*H | -4.8155 | 0.0144 | hypothetical protein |
| BCO26_2663 | *pur*S | -4.7138 | 0.0000 | phosphoribosylformylglycinamidine synthase, purS |
| BCO26_2429 | *yua*F | -4.4123 | 0.0015 | hypothetical protein |
| BCO26_0140 | - | -4.3427 | 0.0049 | hypothetical protein |
| BCO26_2003 | *ycg*M | -4.2934 | 0.0000 | Proline dehydrogenase |
| BCO26_0828 | - | -4.2739 | 0.0000 | binding-protein-dependent transport systems inner membrane component |
| BCO26_2660 | *pur*F | -4.1184 | 0.0013 | amidophosphoribosyltransferase |
| BCO26_2659 | *pur*M | -4.1062 | 0.0006 | phosphoribosylformylglycinamidine cyclo-ligase |
| BCO26_1866 | - | -3.9611 | 0.0208 | Tfp pilus assembly protein ATPase PilM-like protein |
| BCO26_0639 | *lyt*E | -3.9496 | 0.0064 | NLP/P60 protein |
| BCO26_2658 | *pur*N | -3.8162 | 0.0010 | phosphoribosylglycinamide formyltransferase |
| BCO26_0070 | - | -3.7475 | 0.0228 | hypothetical protein |
| BCO26_2664 | *pur*C | -3.7138 | 0.0000 | phosphoribosylaminoimidazole-succinocarboxamide synthase |
| BCO26_2661 | *pur*L | -3.6640 | 0.0000 | phosphoribosylformylglycinamidine synthase II |
| BCO26_2428 | *yua*G | -3.5600 | 0.0026 | hypothetical protein |
| BCO26_1865 | - | -3.5231 | 0.0208 | Fimbrial assembly family protein |
| BCO26_0395 | - | -3.5059 | 0.0077 | cell envelope-related transcriptional attenuator |
| BCO26_0982 | *cta*B | -3.3928 | 0.0004 | protoheme IX farnesyltransferase |
| BCO26_1441 | *yhc*I | -3.3455 | 0.0363 | hypothetical protein |

| | | | | |
|---|---|---|---|---|
| BCO26_1869 | - | -3.2651 | 0.0285 | twitching motility protein |
| BCO26_1530 | *pan*B | -3.1314 | 0.0000 | 3-methyl-2-oxobutanoate hydroxymethyltransferase |
| BCO26_1198 | *yuf*O | -2.9613 | 0.0000 | ABC transporter-like protein |
| BCO26_0872 | *nat*B | -2.9289 | 0.0031 | ABC-2 type transporter |
| BCO26_2467 | *fru*R | -2.8028 | 0.0234 | DeoR family transcriptional regulator |
| BCO26_2268 | *pit* | -2.7363 | 0.0005 | phosphate transporter |
| BCO26_1235 | *csp*D | -2.7183 | 0.0116 | cold-shock DNA-binding domain-containing protein |
| BCO26_0254 | *czc*D | -2.6722 | 0.0069 | cation diffusion facilitator family transporter |
| BCO26_1529 | *pan*C | -2.6180 | 0.0005 | pantoate/beta-alanine ligase |
| BCO26_0376 | *lyt*D | -2.5334 | 0.0152 | Mannosyl-glycoprotein endo-beta-N-acetylglucosaminidase |
| BCO26_0312 | *yub*B | -2.5161 | 0.0009 | undecaprenol kinase |
| BCO26_2931 | - | -2.5026 | 0.0214 | polar amino acid ABC transporter inner membrane subunit |
| BCO26_2798 | *atp*F | -2.4974 | 0.0009 | ATP synthase F0 subunit B |
| BCO26_0795 | - | -2.4424 | 0.0010 | hypothetical protein |
| BCO26_1871 | - | -2.3884 | 0.0024 | hypothetical protein |
| BCO26_0871 | *nat*A | -2.3256 | 0.0199 | ABC transporter-like protein |
| BCO26_0126 | *rpl*P | -2.3083 | 0.0250 | 50S ribosomal protein L16 |
| BCO26_1980 | *ald* | -2.2652 | 0.0125 | alanine dehydrogenase |
| BCO26_1197 | *yuf*N | -2.2543 | 0.0003 | basic membrane lipoprotein |
| BCO26_2802 | *gly*A | -2.2229 | 0.0105 | glycine hydroxymethyltransferase |
| BCO26_0134 | *rpl*R | -2.1865 | 0.0018 | 50S ribosomal protein L18 |
| BCO26_2796 | *atp*A | -2.1209 | 0.0114 | ATP synthase F1 subunit alpha |
| BCO26_1438 | *mnt*H | -2.0894 | 0.0056 | Mn2+/Fe2+ transporter, NRAMP family |
| BCO26_0981 | *cta*A | -2.0466 | 0.0481 | cytochrome oxidase assembly |
| BCO26_0640 | - | -2.0442 | 0.0101 | methyl-accepting chemotaxis sensory transducer |
| BCO26_0142 | *rps*M | -2.0159 | 0.0302 | 30S ribosomal protein S13 |

| | | | | |
|---|---|---|---|---|
| BCO26_0125 | *rps*C | -2.0037 | 0.0245 | 30S ribosomal protein S3 |
| BCO26_0443 | - | -1.9891 | 0.0116 | glycerol dehydrogenase |
| BCO26_0827 | - | -1.9859 | 0.0053 | family 5 extracellular solute-binding protein |
| BCO26_1125 | *hsl*U | -1.9747 | 0.0015 | heat shock protein HslVU, ATPase subunit HslU |
| BCO26_0132 | *rps*H | -1.9086 | 0.0228 | 30S ribosomal protein S8 |
| BCO26_0135 | *rps*E | -1.8813 | 0.0083 | 30S ribosomal protein S5 |
| BCO26_0530 | - | -1.8492 | 0.0488 | hypothetical protein |
| BCO26_2267 | *yka*A | -1.8447 | 0.0281 | hypothetical protein |
| BCO26_1126 | *cod*Y | -1.8339 | 0.0467 | GTP-sensing pleiotropic transcriptional repressor CodY |
| BCO26_0531 | *ldh* | -1.7642 | 0.0254 | L-lactate dehydrogenase |
| BCO26_1253 | - | -1.7337 | 0.0467 | methyl-accepting chemotaxis sensory transducer |
| BCO26_2665 | *pur*B | -1.7075 | 0.0481 | adenylosuccinate lyase |
| BCO26_0660 | *hag* | -1.6965 | 0.0374 | flagellin |
| BCO26_0670 | - | -1.6779 | 0.0458 | Gamma-glutamyltransferase |

*$Log_2$FC：the logarithm of fold change, **Q-value：adjustment of *P*-value

**Table S3. Up-regulated proteins in *B.coagulans* 2-6 by iTRAQ**

| Locus | Gene | $log_2FC$* | Q-value** | Description |
|---|---|---|---|---|
| BCO26_0136 | *rpmD* | 7.0227 | 0.0220 | 50S ribosomal protein L30 |
| BCO26_2925 | - | 5.9648 | 0.0001 | Uncharacterized protein |
| BCO26_0880 | - | 5.8541 | 0.0001 | UPF0180 protein BCO26_0880 |
| BCO26_1317 | *cspL* | 5.6614 | 0.0000 | Cold-shock DNA-binding domain protein |
| BCO26_0255 | *glmS* | 5.5967 | 0.3156 | Glutamine--fructose-6-phosphate aminotransferase |
| BCO26_2102 | *copZ* | 5.5448 | 0.0001 | Copper ion binding protein |
| BCO26_2080 | - | 5.5186 | 0.0000 | Ferritin Dps family protein |
| BCO26_0548 | *ysnF* | 5.4055 | 0.0000 | Uncharacterized protein |
| BCO26_0504 | - | 5.2885 | 0.0000 | Uncharacterized protein |
| BCO26_1740 | *yflL* | 5.0693 | 0.0000 | Acylphosphatase (EC 3.6.1.7) |
| BCO26_0043 | - | 4.8450 | 0.0001 | Transcriptional regulator, AbrB family |
| BCO26_0141 | *infA* | 4.7946 | 0.0000 | Translation initiation factor IF-1 |
| BCO26_0425 | - | 4.7523 | 0.0000 | Glycine betaine/L-proline ABC transporter, ATPase subunit |
| BCO26_1491 | *yneJ* | 4.5668 | 0.0140 | Uncharacterized protein |
| BCO26_2638 | *gsiB* | 4.4977 | 0.0184 | General stress protein |
| BCO26_0441 | - | 4.4727 | 0.0025 | Uncharacterized protein |
| BCO26_0164 | - | 4.4092 | 0.0000 | Cytochrome bd ubiquinol oxidase subunit I |
| BCO26_2375 | - | 4.2469 | 0.0000 | Major facilitator superfamily MFS_1 |
| BCO26_0525 | - | 4.2197 | 0.0589 | Transcriptional regulator |
| BCO26_1438 | *mntH* | 4.1401 | 0.0000 | Divalent metal cation transporter MntH |
| BCO26_1778 | - | 4.0029 | 0.0000 | Heat shock protein Hsp20 |
| BCO26_1458 | *proB* | 3.9946 | 0.0007 | Glutamate 5-kinase (EC 2.7.2.11) (Gamma-glutamyl kinase) (GK) |
| BCO26_0167 | - | 3.7879 | 0.0002 | ABC transporter, CydDC cysteine exporter (CydDC-E) family, permease/ATP-binding |

| | | | | |
|---|---|---|---|---|
| | | | | protein CydC |
| BCO26_0423 | - | 3.7786 | 0.0005 | Substrate-binding region of ABC-type glycine betaine transport system |
| BCO26_0505 | - | 3.7451 | 0.0002 | Quinone oxidoreductase, YhdH/YhfP family |
| BCO26_0921 | *mrgA* | 3.5279 | 0.0005 | Ferritin Dps family protein |
| BCO26_1788 | - | 3.5157 | 0.0000 | Alkyl hydroperoxide reductase, F subunit |
| BCO26_2932 | *mqo* | 3.5035 | 0.0000 | Probable malate:quinone oxidoreductase |
| BCO26_0241 | - | 3.5013 | 0.0000 | Aldehyde dehydrogenase |
| BCO26_0939 | *ptsH* | 3.4935 | 0.0000 | Phosphotransferase system, phosphocarrier protein HPr |
| BCO26_1725 | *rpoD* | 3.4662 | 0.0004 | RNA polymerase sigma factor SigA |
| BCO26_2724 | *groEL* | 3.3843 | 0.0000 | 60 kDa chaperonin (GroEL protein) (Protein Cpn60) |
| BCO26_0166 | *cydC* | 3.2730 | 0.0000 | ABC transporter, CydDC cysteine exporter (CydDC-E) family, permease/ATP-binding protein CydD |
| BCO26_0424 | - | 3.2591 | 0.0002 | Binding-protein-dependent transport systems inner membrane component |
| BCO26_0610 | - | 3.2583 | 0.0005 | Uncharacterized protein |
| BCO26_0300 | - | 3.1971 | 0.0001 | Glycerol-3-phosphate dehydrogenase (EC 1.1.5.3) |
| BCO26_0207 | *treR* | 3.1921 | 0.0007 | Transcriptional regulator, GntR family |
| BCO26_0092 | *radA* | 3.1826 | 0.0045 | DNA repair protein RadA |
| BCO26_2962 | - | 3.0935 | 0.0029 | Uncharacterized protein |
| BCO26_1721 | - | 3.0791 | 0.0011 | Uncharacterized protein |
| BCO26_1748 | *hrcA* | 3.0552 | 0.0007 | Heat-inducible transcription repressor HrcA |
| BCO26_1648 | - | 3.0521 | 0.0000 | DNA repair protein RecN (Recombination protein N) |
| BCO26_0240 | - | 3.0461 | 0.0001 | Anti-sigma-factor antagonist |
| BCO26_2617 | *htpG* | 3.0134 | 0.0001 | Chaperone protein HtpG (Heat shock protein HtpG) (High temperature protein G) |
| BCO26_2151 | - | 2.9273 | 0.0000 | NADH:flavin oxidoreductase/NADH oxidase |
| BCO26_0934 | - | 2.9246 | 0.0000 | ATPase AAA-2 domain protein |
| BCO26_1052 | - | 2.9075 | 0.0038 | Uncharacterized protein |

| | | | | |
|---|---|---|---|---|
| BCO26_2898 | - | 2.9038 | 0.0000 | Drug resistance transporter, EmrB/QacA subfamily |
| BCO26_2515 | *argD* | 2.9004 | 0.0742 | Multifunctional fusion protein |
| BCO26_2685 | - | 2.8904 | 0.0001 | Anion transporter |
| BCO26_2513 | - | 2.8499 | 0.0000 | FAD-dependent pyridine nucleotide-disulfide oxidoreductase |
| BCO26_0308 | - | 2.8427 | 0.0003 | Metal dependent phosphohydrolase |
| BCO26_0586 | - | 2.8244 | 0.0002 | BAAT/Acyl-CoA thioester hydrolase |
| BCO26_2416 | *yfkM* | 2.8229 | 0.0000 | Intracellular protease, PfpI family |
| BCO26_0589 | - | 2.8102 | 0.0000 | Aldo/keto reductase |
| BCO26_0735 | - | 2.7917 | 0.0001 | Phosphotransferase system, phosphocarrier protein HPr |
| BCO26_1709 | - | 2.7618 | 0.0001 | Nucleotidase (EC 3.1.3.-) |
| BCO26_0496 | *metC* | 2.7363 | 0.0000 | Cystathionine gamma-synthase |
| BCO26_0298 | *glpK* | 2.6713 | 0.0032 | Glycerol kinase |
| BCO26_1429 | - | 2.6689 | 0.0000 | 3-hydroxyisobutyrate dehydrogenase |
| BCO26_1771 | - | 2.6197 | 0.0095 | Heat shock protein Hsp20 |
| BCO26_1118 | *sucD* | 2.5942 | 0.0000 | Succinate--CoA ligase [ADP-forming] subunit alpha (EC 6.2.1.5) (Succinyl-CoA synthetase subunit alpha) (SCS-alpha) |
| BCO26_2126 | - | 2.5806 | 0.0032 | PTS system, glucose subfamily, IIA subunit |
| BCO26_0653 | - | 2.5640 | 0.0020 | YvyF |
| BCO26_2482 | *yhdN* | 2.5571 | 0.0000 | Aldo/keto reductase |
| BCO26_0731 | - | 2.5548 | 0.0000 | NUDIX hydrolase |
| BCO26_1746 | *dnaK* | 2.5337 | 0.0000 | Chaperone protein DnaK (HSP70) (Heat shock 70 kDa protein) (Heat shock protein 70) |
| BCO26_2725 | *groES* | 2.5200 | 0.0153 | 10 kDa chaperonin (GroES protein) (Protein Cpn10) |
| BCO26_2167 | - | 2.4931 | 0.0873 | Peptidase M17 leucyl aminopeptidase domain protein |
| BCO26_0541 | *ptnA* | 2.4777 | 0.0000 | PTS system, mannose/fructose/sorbose family, IIA subunit |
| BCO26_2743 | *rsbV* | 2.4404 | 0.0001 | Anti-sigma factor antagonist |
| BCO26_1223 | *mutL* | 2.4133 | 0.0002 | DNA mismatch repair protein MutL |

| | | | | |
|---|---|---|---|---|
| BCO26_0186 | *cobB* | 2.4064 | 0.0012 | NAD-dependent protein deacetylase (EC 3.5.1.-) (Regulatory protein SIR2 homolog) |
| BCO26_0660 | - | 2.4053 | 0.0000 | Flagellin |
| BCO26_0420 | *galR* | 2.3949 | 0.0029 | Transcriptional regulator, LacI family |
| BCO26_0213 | - | 2.3943 | 0.0030 | Flavodoxin/nitric oxide synthase |
| BCO26_2103 | - | 2.3803 | 0.0026 | Uncharacterized protein |
| BCO26_1978 | - | 2.3657 | 0.0001 | UPF0173 metal-dependent hydrolase BCO26_1978 |
| BCO26_0259 | - | 2.3632 | 0.0000 | NAD-dependent epimerase/dehydratase |
| BCO26_0874 | - | 2.3489 | 0.0000 | Uncharacterized protein |
| BCO26_0832 | *spxA* | 2.3374 | 0.0170 | Regulatory protein Spx |
| BCO26_0587 | - | 2.3109 | 0.0041 | Esterase/lipase-like protein |
| BCO26_1784 | - | 2.2983 | 0.0001 | Uncharacterized protein |
| BCO26_2957 | *ribH* | 2.2752 | 0.0011 | 6,7-dimethyl-8-ribityllumazine synthase (DMRL synthase) (LS) (Lumazine synthase) (EC 2.5.1.78) |
| BCO26_0402 | *yfmJ* | 2.2720 | 0.0000 | Alcohol dehydrogenase zinc-binding domain protein |
| BCO26_1841 | *yrbC* | 2.2719 | 0.0000 | Probable transcriptional regulatory protein BCO26_1841 |
| BCO26_1768 | - | 2.2708 | 0.0020 | Glycolate oxidase, subunit GlcD |
| BCO26_2495 | - | 2.2634 | 0.0000 | Nitroreductase |
| BCO26_1789 | - | 2.2622 | 0.0000 | Peroxiredoxin |
| BCO26_0938 | - | 2.2612 | 0.0032 | Uncharacterized protein |
| BCO26_2307 | - | 2.2471 | 0.0001 | Ferric uptake regulator, Fur family |
| BCO26_1359 | *plsY* | 2.2396 | 0.0001 | Glycerol-3-phosphate acyltransferase (Acyl-PO4 G3P acyltransferase) (Acyl-phosphate--glycerol-3-phosphate acyltransferase) (G3P acyltransferase) (GPAT) (EC 2.3.1.n3) (Lysophosphatidic acid synthase) (LPA synthase) |
| BCO26_0780 | *yhgD* | 2.2154 | 0.0005 | Transcriptional regulator, TetR family |
| BCO26_1977 | - | 2.2027 | 0.0057 | Putative signal transduction protein with CBS and DRTGG domains |
| BCO26_0622 | *galK* | 2.2010 | 0.0016 | Galactokinase (EC 2.7.1.6) (Galactose kinase) |

| | | | | |
|---|---|---|---|---|
| BCO26_2021 | - | 2.1981 | 0.0057 | Molybdenum ABC transporter, periplasmic molybdate-binding protein |
| BCO26_1454 | *selD* | 2.1598 | 0.0000 | Selenide, water dikinase (EC 2.7.9.3) (Selenium donor protein) (Selenophosphate synthase) |
| BCO26_1764 | *aroE* | 2.1254 | 0.0024 | Shikimate dehydrogenase (NADP(+)) (SDH) (EC 1.1.1.25) |
| BCO26_0165 | *cydB* | 2.1109 | 0.0004 | Cytochrome d ubiquinol oxidase, subunit II |
| BCO26_1679 | - | 2.0960 | 0.0000 | Vitamin B12-dependent ribonucleotide reductase (EC 1.17.4.1) |
| BCO26_0263 | - | 2.0749 | 0.0000 | Aminotransferase class I and II |
| BCO26_0171 | - | 2.0426 | 0.0001 | Arginase (EC 3.5.3.1) |
| BCO26_2374 | - | 2.0188 | 0.0003 | Methionine synthase vitamin-B12 independent |
| BCO26_2212 | - | 2.0129 | 0.0000 | Catalase (EC 1.11.1.6) |
| BCO26_1539 | - | 2.0036 | 0.0000 | Peptidase membrane zinc metallopeptidase putative |
| BCO26_1117 | *sucC* | 1.9921 | 0.0005 | Succinate--CoA ligase [ADP-forming] subunit beta (EC 6.2.1.5) (Succinyl-CoA synthetase subunit beta) (SCS-beta) |
| BCO26_0544 | - | 1.9870 | 0.0002 | Uncharacterized protein |
| BCO26_2279 | - | 1.9664 | 0.0042 | Extracellular solute-binding protein family 3 |
| BCO26_2070 | *ytmA* | 1.9553 | 0.0001 | BAAT/Acyl-CoA thioester hydrolase |
| BCO26_1964 | - | 1.9500 | 0.0016 | Citrate synthase |
| BCO26_0190 | - | 1.9480 | 0.0003 | Methyl-accepting chemotaxis sensory transducer with Cache sensor |
| BCO26_2636 | - | 1.9436 | 0.0000 | Diacylglycerol kinase catalytic region |
| BCO26_2370 | - | 1.9399 | 0.0006 | Anti-sigma-factor antagonist |
| BCO26_0767 | *yhaH* | 1.9251 | 0.0000 | Uncharacterized protein |
| BCO26_2181 | *lipA* | 1.9101 | 0.0002 | Lipoyl synthase (EC 2.8.1.8) |
| BCO26_0434 | - | 1.8896 | 0.0002 | RmlC-like cupin |
| BCO26_2878 | - | 1.8706 | 0.0000 | Cys-tRNA(Pro)/Cys-tRNA(Cys) deacylase (EC 4.2.-.-) |
| BCO26_0543 | *manN* | 1.8642 | 0.0000 | PTS system, mannose/fructose/sorbose family, IID subunit |
| BCO26_2303 | - | 1.8491 | 0.0000 | Acetolactate synthase, catabolic |

| | | | | |
|---|---|---|---|---|
| BCO26_0306 | *yqiG* | 1.8249 | 0.0000 | NADH:flavin oxidoreductase/NADH oxidase |
| BCO26_2556 | - | 1.8216 | 0.0001 | Heavy metal translocating P-type ATPase |
| BCO26_2595 | *deoC* | 1.8132 | 0.0000 | Deoxyribose-phosphate aldolase |

*Log2FC：the logarithm of fold change, **Q-value：adjustment of *P*-value

**Table S4. Down-regulated proteins in *B.coagulans* 2-6 by iTRAQ**

| Locus | Gene | $log_2FC$* | Q-value** | Description |
|---|---|---|---|---|
| BCO26_2160 | - | -7.5824 | 0.0058 | Thioesterase superfamily protein |
| BCO26_0352 | - | -7.5092 | 0.0001 | dTDP-4-dehydrorhamnose reductase |
| BCO26_0361 | - | -6.9636 | 0.3778 | Uncharacterized protein |
| BCO26_2488 | *ureG* | -6.7583 | 0.0001 | Urease accessory protein UreG |
| BCO26_0519 | - | -6.6545 | 0.0003 | Cell envelope-related transcriptional attenuator |
| BCO26_2428 | - | -6.4803 | 0.0000 | Band 7 protein |
| BCO26_2275 | *queG* | -6.4136 | 0.0190 | Epoxyqueuosine reductase |
| BCO26_0819 | - | -5.9874 | 0.0077 | Oligopeptide/dipeptide ABC transporter, ATPase subunit |
| BCO26_1048 | *pyrD* | -5.9114 | 0.0000 | Dihydroorotate dehydrogenase |
| BCO26_0034 | *spoVG* | -5.7888 | 0.0000 | Putative septation protein SpoVG (Stage V sporulation protein G) |
| BCO26_2663 | *purS* | -5.6376 | 0.0001 | Phosphoribosylformylglycinamidine synthase subunit PurS (FGAM synthase) |
| BCO26_2368 | *ilvD* | -5.5863 | 0.0004 | Dihydroxy-acid dehydratase (DAD) (EC 4.2.1.9) |
| BCO26_1650 | - | -5.5171 | 0.0000 | Hemolysin A |
| BCO26_2174 | - | -5.4611 | 0.0003 | Uncharacterized protein |
| BCO26_2285 | *pflA* | -5.3524 | 0.0001 | Pyruvate formate-lyase-activating enzyme (EC 1.97.1.4) |
| BCO26_1607 | *spoIIAB* | -5.0539 | 0.0028 | Anti-sigma F factor (EC 2.7.11.1) (Stage II sporulation protein AB) |
| BCO26_1393 | *yfiR* | -4.9993 | 0.0045 | Transcriptional regulator, TetR family |
| BCO26_0894 | *yfjA* | -4.8611 | 0.0008 | ESAT-6-like protein |
| BCO26_1282 | - | -4.8083 | 0.0000 | UPF0176 protein BCO26_1282 |
| BCO26_0828 | - | -4.7767 | 0.0000 | Binding-protein-dependent transport systems inner membrane component |
| BCO26_1240 | - | -4.7517 | 0.0001 | ABC transporter related protein |
| BCO26_1484 | *cspD* | -4.7356 | 0.0002 | Cold-shock DNA-binding domain protein |
| BCO26_1941 | - | -4.6901 | 0.0043 | 2-dehydropantoate 2-reductase (EC 1.1.1.169) (Ketopantoate reductase) |

| | | | | |
|---|---|---|---|---|
| BCO26_1608 | - | -4.6612 | 0.0000 | Anti-sigma F factor antagonist (Stage II sporulation protein) |
| BCO26_1403 | - | -4.6556 | 0.0012 | Transcriptional regulator, MerR family |
| BCO26_2618 | - | -4.6063 | 0.0002 | Terpenoid cyclases/Protein prenyltransferase |
| BCO26_1933 | - | -4.5611 | 0.0000 | tRNA/rRNA methyltransferase (SpoU) |
| BCO26_2665 | - | -4.5541 | 0.0001 | Adenylosuccinate lyase (ASL) (EC 4.3.2.2) (Adenylosuccinase) |
| BCO26_1864 | - | -4.5241 | 0.0064 | Uncharacterized protein |
| BCO26_0555 | - | -4.4862 | 0.0001 | ABC transporter related protein |
| BCO26_0347 | - | -4.3992 | 0.0004 | Putative glycosyl transferase |
| BCO26_0280 | - | -4.3586 | 0.0001 | TPR-like protein |
| BCO26_2119 | - | -4.3128 | 0.0000 | Polysaccharide deacetylase |
| BCO26_2310 | - | -4.2956 | 0.0278 | ATP/cobalamin adenosyltransferase |
| BCO26_1507 | *xpt* | -4.2842 | 0.0002 | Xanthine phosphoribosyltransferase (XPRTase) (EC 2.4.2.22) |
| BCO26_0443 | *ypjH* | -4.2763 | 0.0000 | Glycerol dehydrogenase |
| BCO26_2156 | - | -4.2631 | 0.0000 | Peptidyl-prolyl cis-trans isomerase (PPIase) (EC 5.2.1.8) |
| BCO26_1894 | *ysoA* | -4.1856 | 0.0001 | TPR-like protein |
| BCO26_0075 | - | -4.1131 | 0.0001 | Deoxynucleoside kinase |
| BCO26_0584 | - | -4.1129 | 0.0000 | Cof-like hydrolase |
| BCO26_1556 | *ndk* | -4.0916 | 0.0915 | Nucleoside diphosphate kinase (NDK) (NDP kinase) (EC 2.7.4.6) (Nucleoside-2-P kinase) |
| BCO26_1044 | *pyrC* | -4.0482 | 0.0004 | Dihydroorotase (DHOase) (EC 3.5.2.3) |
| BCO26_1036 | - | -4.0019 | 0.0027 | RNA-binding S4 domain protein |
| BCO26_1043 | *pyrB* | -3.9945 | 0.0000 | Aspartate carbamoyltransferase (EC 2.1.3.2) (Aspartate transcarbamylase) (ATCase) |
| BCO26_1856 | *minC* | -3.9420 | 0.0001 | Probable septum site-determining protein MinC |
| BCO26_2364 | *leuA* | -3.8793 | 0.0752 | 2-isopropylmalate synthase (EC 2.3.3.13) (Alpha-IPM synthase) (Alpha-isopropylmalate synthase) |
| BCO26_0567 | - | -3.8746 | 0.0001 | Aldehyde-alcohol dehydrogenase |

| | | | | |
|---|---|---|---|---|
| BCO26_1371 | - | -3.8619 | 0.0029 | 2-nitropropane dioxygenase NPD |
| BCO26_0726 | - | -3.8503 | 0.0000 | HAD-superfamily hydrolase, subfamily IA, variant 1 |
| BCO26_2917 | - | -3.8472 | 0.0001 | TIGR00697: conserved hypothetical integral |
| BCO26_0670 | - | -3.7142 | 0.0003 | Gamma-glutamyltransferase |
| BCO26_0343 | - | -3.7141 | 0.0002 | NAD-dependent epimerase/dehydratase |
| BCO26_1529 | *panC* | -3.6906 | 0.0001 | Pantothenate synthetase (PS) (EC 6.3.2.1) (Pantoate--beta-alanine ligase) (Pantoate-activating enzyme) |
| BCO26_2768 | *malL* | -3.6481 | 0.0000 | Oligo-1,6-glucosidase |
| BCO26_1408 | *acoC* | -3.6025 | 0.0162 | Dihydrolipoamide acetyltransferase component of pyruvate dehydrogenase complex (EC 2.3.1.-) |
| BCO26_1049 | *pyrF* | -3.5786 | 0.0008 | Orotidine 5'-phosphate decarboxylase (EC 4.1.1.23) (OMP decarboxylase) (OMPDCase) (OMPdecase) |
| BCO26_1045 | *carA* | -3.5253 | 0.0000 | Carbamoyl-phosphate synthase small chain (EC 6.3.5.5) (Carbamoyl-phosphate synthetase glutamine chain) |
| BCO26_1173 | *rimP* | -3.4935 | 0.0012 | Ribosome maturation factor RimP |
| BCO26_0829 | *oppC* | -3.4615 | 0.0000 | Binding-protein-dependent transport systems inner membrane component |
| BCO26_1722 | - | -3.4578 | 0.0028 | Uncharacterized protein |
| BCO26_1652 | - | -3.4278 | 0.0087 | Polyprenyl synthetase |
| BCO26_2619 | - | -3.4053 | 0.0003 | Uncharacterized protein |
| BCO26_2545 | - | -3.3487 | 0.0322 | Type I site-specific deoxyribonuclease, HsdR family |
| BCO26_0296 | - | -3.3480 | 0.0006 | Glycerol uptake operon antiterminator regulatory protein |
| BCO26_0202 | - | -3.3374 | 0.0006 | Glycine betaine/L-proline ABC transporter, ATPase subunit |
| BCO26_1166 | *uppS* | -3.3127 | 0.0014 | Isoprenyl transferase (EC 2.5.1.-) |
| BCO26_1632 | - | -3.2619 | 0.0023 | CheW protein |
| BCO26_0871 | - | -3.2586 | 0.0004 | ABC transporter related protein |
| BCO26_1040 | *lspA* | -3.2580 | 0.0002 | Lipoprotein signal peptidase (EC 3.4.23.36) (Prolipoprotein signal peptidase) (Signal |

| | | | | |
|---|---|---|---|---|
| | | | | peptidase II) (SPase II) |
| BCO26_1384 | *thiE* | -3.2397 | 0.0000 | Thiamine-phosphate synthase (TP synthase) (TPS) (EC 2.5.1.3) (Thiamine-phosphate pyrophosphorylase) (TMP pyrophosphorylase) (TMP-PPase) |
| BCO26_1942 | - | -3.1936 | 0.0043 | NADPH-dependent FMN reductase |
| BCO26_1831 | *yrzD* | -3.1820 | 0.0392 | Uncharacterized protein |
| BCO26_1720 | *ispH* | -3.1819 | 0.0004 | 4-hydroxy-3-methylbut-2-enyl diphosphate reductase (EC 1.17.7.4) |
| BCO26_0831 | - | -3.1643 | 0.0001 | ABC transporter related protein |
| BCO26_0678 | - | -3.1311 | 0.0000 | Peptidase M23 |
| BCO26_0568 | *deaD* | -3.1212 | 0.0000 | DEAD/DEAH box helicase domain protein |
| BCO26_1386 | *yflK* | -3.1057 | 0.0050 | MOSC domain containing protein |
| BCO26_2362 | *leuC* | -3.1016 | 0.0001 | 3-isopropylmalate dehydratase large subunit (EC 4.2.1.33) (Alpha-IPM isomerase) (IPMI) (Isopropylmalate isomerase) |
| BCO26_2873 | - | -3.0291 | 0.0000 | Uncharacterized protein |
| BCO26_0174 | *ybbM* | -3.0113 | 0.0027 | Putative transmembrane anti-sigma factor |
| BCO26_1761 | - | -2.9811 | 0.0247 | Metal dependent phosphohydrolase |
| BCO26_0532 | - | -2.9766 | 0.0004 | 3D domain protein |
| BCO26_1544 | *qcrA* | -2.8935 | 0.0102 | Rieske (2Fe-2S) domain protein |
| BCO26_2123 | - | -2.8916 | 0.0000 | Rhodanese domain protein |
| BCO26_2657 | *purH* | -2.8563 | 0.0008 | Bifunctional purine biosynthesis protein PurH [Includes: IMP cyclohydrolase (EC 3.5.4.10) (IMP synthase) (Inosinicase) (ATIC); Phosphoribosylaminoimidazolecarboxamide formyltransferase (EC 2.1.2.3) (AICAR transformylase)] |
| BCO26_1657 | *nusB* | -2.8266 | 0.0001 | N utilization substance protein B homolog (Protein NusB) |
| BCO26_2903 | *deoC* | -2.8003 | 0.0000 | Deoxyribose-phosphate aldolase (DERA) (EC 4.1.2.4) (2-deoxy-D-ribose 5-phosphate aldolase) (Phosphodeoxyriboaldolase) (Deoxyriboaldolase) |
| BCO26_1314 | *msrA* | -2.7925 | 0.0000 | Peptide methionine sulfoxide reductase MsrA (Protein-methionine-S-oxide reductase) |

| | | | | |
|---|---|---|---|---|
| | | | | (EC 1.8.4.11) (Peptide-methionine (S)-S-oxide reductase) (Peptide Met(O) reductase) |
| BCO26_0638 | - | -2.7451 | 0.0000 | Uncharacterized protein |
| BCO26_1200 | - | -2.7242 | 0.0001 | Inner-membrane translocator |
| BCO26_0203 | - | -2.7112 | 0.0000 | Substrate-binding region of ABC-type glycine betaine transport system |
| BCO26_1992 | - | -2.6967 | 0.0000 | AMP-dependent synthetase and ligase |
| BCO26_2924 | - | -2.6938 | 0.0002 | Carboxynorspermidine decarboxylase |
| BCO26_0811 | - | -2.6897 | 0.0007 | Uncharacterized protein |
| BCO26_0872 | *yhaP* | -2.6678 | 0.0003 | ABC-2 type transporter |
| BCO26_2688 | - | -2.6461 | 0.0000 | Uncharacterized protein |
| BCO26_0830 | - | -2.6101 | 0.0000 | Oligopeptide/dipeptide ABC transporter, ATPase subunit |
| BCO26_1497 | *ynzC* | -2.6075 | 0.0001 | UPF0291 protein BCO26_1497 |
| BCO26_0728 | - | -2.5846 | 0.0022 | Tetratricopeptide TPR_2 repeat protein |
| BCO26_2922 | - | -2.5739 | 0.0001 | Anti-sigma-factor antagonist |
| BCO26_2664 | *purC* | -2.5465 | 0.0000 | Phosphoribosylaminoimidazole-succinocarboxamide synthase (EC 6.3.2.6) (SAICAR synthetase) |
| BCO26_2491 | *ureC* | -2.5391 | 0.0001 | Urease subunit alpha (EC 3.5.1.5) (Urea amidohydrolase subunit alpha) |
| BCO26_0304 | - | -2.5149 | 0.0000 | Uncharacterized protein |
| BCO26_2547 | - | -2.4971 | 0.0004 | Type I restriction-modification system, M subunit |
| BCO26_1936 | - | -2.4754 | 0.0001 | Uncharacterized protein |
| BCO26_2592 | *guaC* | -2.4506 | 0.0000 | GMP reductase (EC 1.7.1.7) (Guanosine 5'-monophosphate oxidoreductase) (Guanosine monophosphate reductase) |
| BCO26_1797 | *udk* | -2.4486 | 0.0129 | Uridine kinase (EC 2.7.1.48) (Cytidine monophosphokinase) (Uridine monophosphokinase) |
| BCO26_1601 | - | -2.4359 | 0.0000 | Diaminopimelate decarboxylase (EC 4.1.1.20) |
| BCO26_2821 | - | -2.4351 | 0.0008 | Response regulator receiver protein |
| BCO26_0554 | - | -2.4201 | 0.0002 | NMT1/THI5 like domain protein |

| | | | | |
|---|---|---|---|---|
| BCO26_2649 | *pcrA* | -2.4122 | 0.0003 | DNA helicase (EC 3.6.4.12) |
| BCO26_1775 | - | -2.4005 | 0.0007 | SirA family protein |
| BCO26_0345 | - | -2.3925 | 0.0003 | Putative lipopolysaccharide biosynthesis protein |
| BCO26_1046 | *carB* | -2.3867 | 0.0001 | Carbamoyl-phosphate synthase large chain (EC 6.3.5.5) (Carbamoyl-phosphate synthetase ammonia chain) |
| BCO26_2739 | - | -2.3688 | 0.0000 | RNA binding S1 domain protein |
| BCO26_2384 | - | -2.3454 | 0.0000 | Chromosome segregation ATPase-like protein |
| BCO26_2950 | *dnaC* | -2.3363 | 0.0042 | Replicative DNA helicase (EC 3.6.4.12) |
| BCO26_2606 | | -2.3146 | 0.0003 | Uncharacterized protein |
| BCO26_1890 | *lonA lon* | -2.3133 | 0.0000 | Lon protease (EC 3.4.21.53) (ATP-dependent protease La) |
| BCO26_0057 | *hprT* | -2.2931 | 0.0000 | Hypoxanthine phosphoribosyltransferase |
| BCO26_1527 | - | -2.2737 | 0.0126 | DnaQ family exonuclease/DinG family helicase |
| BCO26_1243 | - | -2.2705 | 0.0000 | Uncharacterized protein |
| BCO26_0391 | - | -2.2550 | 0.0013 | Arabinogalactan endo-beta-1,4-galactanase (EC 3.2.1.89) |
| BCO26_2814 | *tdk* | -2.2521 | 0.0000 | Thymidine kinase (EC 2.7.1.21) |
| BCO26_0232 | - | -2.2426 | 0.0001 | Periplasmic binding protein |
| BCO26_0348 | - | -2.2171 | 0.0002 | NAD-dependent epimerase/dehydratase |
| BCO26_1953 | *dnaB* | -2.2143 | 0.0031 | Replication initiation and membrane attachment family protein |
| BCO26_2386 | - | -2.2108 | 0.0136 | Uncharacterized protein |
| BCO26_1435 | - | -2.2091 | 0.0004 | DSBA oxidoreductase |
| BCO26_1927 | - | -2.1915 | 0.0004 | Cell division protein ZapA |
| BCO26_1230 | *hflX* | -2.1667 | 0.0000 | GTPase HflX (GTP-binding protein HflX) |
| BCO26_2476 | *rlmN* | -2.1596 | 0.0000 | Probable dual-specificity RNA methyltransferase RlmN (EC 2.1.1.192) (23S rRNA (adenine(2503)-C(2))-methyltransferase) (23S rRNA m2A2503 methyltransferase) (Ribosomal RNA large subunit methyltransferase N) (tRNA (adenine(37)-C(2))-methyltransferase) (tRNA m2A37 methyltransferase) |

| | | | | |
|---|---|---|---|---|
| BCO26_2304 | - | -2.1399 | 0.0008 | Transcriptional regulator, LysR family |
| BCO26_2608 | - | -2.1276 | 0.0236 | Urea amidolyase related protein |
| BCO26_2644 | - | -2.1254 | 0.0003 | Malate synthase (EC 2.3.3.9) |
| BCO26_1050 | *pyrE* | -2.1166 | 0.0000 | Orotate phosphoribosyltransferase (OPRT) (OPRTase) (EC 2.4.2.10) |
| BCO26_2693 | - | -2.1107 | 0.0000 | Transposase IS4 family protein |
| BCO26_0427 | - | -2.0539 | 0.0000 | Transferase hexapeptide repeat containing protein |
| BCO26_0083 | - | -2.0469 | 0.0021 | Uncharacterized protein |
| BCO26_1810 | *mnmA* | -2.0469 | 0.0017 | tRNA-specific 2-thiouridylase MnmA (EC 2.8.1.13) |
| BCO26_2648 | *pcrA* | -2.0207 | 0.0001 | DNA helicase (EC 3.6.4.12) |
| BCO26_1922 | - | -2.0203 | 0.0003 | Transcriptional regulator, TetR family |
| BCO26_1101 | *smc* | -2.0146 | 0.0000 | Chromosome partition protein Smc |
| BCO26_1687 | *aroK* | -2.0087 | 0.0011 | Shikimate kinase (SK) (EC 2.7.1.71) |
| BCO26_1998 | - | -1.9980 | 0.0005 | Putative GAF sensor protein |
| BCO26_1636 | *fruK* | -1.9887 | 0.0003 | 1-phosphofructokinase |
| BCO26_0019 | - | -1.9715 | 0.0035 | Methyltransferase small |
| BCO26_1509 | - | -1.9706 | 0.0001 | Putative RNA methylase |
| BCO26_2801 | *upp* | -1.9023 | 0.0000 | Uracil phosphoribosyltransferase (EC 2.4.2.9) (UMP pyrophosphorylase) (UPRTase) |
| BCO26_0267 | - | -1.8989 | 0.0008 | UPF0210 protein BCO26_0267 |
| BCO26_1112 | - | -1.8932 | 0.0001 | Signal peptidase I (EC 3.4.21.89) |
| BCO26_2472 | *yxeH* | -1.8477 | 0.0000 | Cof-like hydrolase |
| BCO26_1008 | *bshC* | -1.7849 | 0.0000 | Putative cysteine ligase BshC (EC 6.-.-.-) |

*Log2FC：the logarithm of fold change, **Q-value：adjustment of *P*-value

**Table S5. Commonly differentially accumulated in the RNA-seq and iTRAQ for *B. coagulans* 2-6**

| Locus | Gene | $log_2$(37/60) | Product |
|---|---|---|---|
| BCO26_2925 | - | 5.96 | hypothetical protein |
| BCO26_0880 | *yku*S | 5.85 | hypothetical protein |
| BCO26_1317 | *csp*C | 5.66 | cold-shock DNA-binding domain-containing protein |
| BCO26_2080 | *dps* | 5.51 | Ferritin dps family protein |
| BCO26_2638 | *gsi*B | 4.49 | general stress protein |
| BCO26_2375 | *ybf*B | 4.24 | major facilitator superfamily protein |
| BCO26_1438 | *mnt*H | 4.14 | $Mn^{2+}$/$Fe^{2+}$ transporter, NRAMP family |
| BCO26_2932 | - | 3.50 | malate/quinone oxidoreductase |
| BCO26_0241 | *gab*D | 3.50 | succinic semialdehyde dehydrogenase |
| BCO26_2724 | *gro*EL | 3.38 | chaperonin GroEL |
| BCO26_1748 | *hrc*A | 3.05 | heat-inducible transcription repressor HrcA |
| BCO26_0934 | *clp*E | 2.92 | ATPase AAA-2 domain-containing protein |
| BCO26_1771 | - | 2.61 | heat shock protein Hsp20 |
| BCO26_1746 | *dna*K | 2.53 | chaperone protein DnaK |
| BCO26_2725 | *gro*ES | 2.51 | chaperonin Cpn10 |
| BCO26_0541 | - | 2.47 | PTS system, mannose/fructose/sorbose family, IIA subunit |
| BCO26_2743 | *rsb*V | 2.44 | anti-sigma-factor antagonist |
| BCO26_0660 | *hag* | 2.40 | flagellin |
| BCO26_0780 | *yhg*D | 2.21 | TetR family transcriptional regulator |
| BCO26_1679 | - | 2.09 | adenosylcobalamin-dependent ribonucleoside-diphosphate reductase |
| BCO26_0171 | *arg*I | 2.04 | arginase |
| BCO26_1117 | *suc*C | 1.99 | succinyl-CoA synthetase subunit beta |
| BCO26_1964 | *cit*Z | 1.94 | 2-methylcitrate synthase/citrate synthase II |
| BCO26_2370 | - | 1.93 | anti-sigma-factor antagonist |

| | | | |
|---|---|---|---|
| BCO26_0543 | *lev*G | 1.86 | PTS system, mannose/fructose/sorbose family, IID subunit |
| BCO26_0306 | *yqi*G | 1.82 | NADH:flavin oxidoreductase/NADH oxidase |
| BCO26_2608 | *kip*A | -2.12 | urea amidolyase-like protein |
| BCO26_2664 | *pur*C | -2.54 | phosphoribosylaminoimidazole-succinocarboxamide synthase |
| BCO26_0872 | *nat*B | -2.66 | ABC-2 type transporter |
| BCO26_0639 | *lyt*E | -2.74 | NLP/P60 protein |
| BCO26_0871 | *nat*A | -3.25 | ABC transporter-like protein |
| BCO26_1529 | *pan*C | -3.69 | pantoate/beta-alanine ligase |
| BCO26_0670 | - | -3.71 | gamma-glutamyltransferase |
| BCO26_0443 | - | -4.27 | glycerol dehydrogenase |
| BCO26_2665 | *pur*B | -4.55 | adenylosuccinate lyase |
| BCO26_0828 | - | -4.77 | binding-protein-dependent transport systems inner membrane component |
| BCO26_2663 | *pur*S | -5.63 | phosphoribosylformylglycinamidine synthase, purS |
| BCO26_2428 | *yua*G | -6.48 | hypothetical protein |

**Table S6. List of the binding mRNA targets of CspL in *E. coli* DH5α by RIP-seq**

| Gene_name | Locus_ID | Description | Foldchange | *P*-Value | FDR | CspL-RIP | Input-RIP |
|---|---|---|---|---|---|---|---|
| acpP | ECD_01090 | acyl carrier protein (ACP) | 3.209574 | 0.011899 | 0.353285 | 1597.78 | 497.6643 |
| acrE | ECD_03123 | cytoplasmic membrane lipoprotein | 5.40998 | 0.033148 | 0.519559 | 20.44069 | 3.597574 |
| acrF | ECD_03124 | multidrug efflux system protein | 3.447072 | 0.02446 | 0.46674 | 74.94919 | 21.58544 |
| actP | ECD_03939 | acetate transporter | 5.017942 | 0.000673 | 0.060613 | 1655.696 | 329.7776 |
| ahpC | ECD_00574 | alkyl hydroperoxide reductase, C22 subunit | 6.277318 | 0.00017 | 0.02059 | 671.1359 | 106.728 |
| aslB | ECD_03675 | putative AslA-specific sulfatase-maturating enzyme | 4.007795 | 0.010328 | 0.320887 | 91.9831 | 22.78463 |
| aspA | ECD_04009 | aspartate ammonia-lyase | 3.189925 | 0.01204 | 0.353285 | 2486.95 | 779.4743 |
| atoB | ECD_02151 | acetyl-CoA acetyltransferase | 3.056902 | 0.021741 | 0.456012 | 235.0679 | 76.74824 |
| atpC | ECD_03615 | F1 sector of membrane-bound ATP synthase, epsilon subunit | 8.418546 | 1.64E-05 | 0.004439 | 960.7124 | 113.9232 |
| bamB | ECD_02404 | BamABCDE complex OM biogenesis lipoprotein | 5.713884 | 0.000256 | 0.028998 | 2241.662 | 392.1355 |
| bax | ECD_03422 | putative glucosaminidase | 4.753606 | 0.000945 | 0.072227 | 2377.933 | 500.0627 |
| bolA | ECD_00386 | stationary-phase morphogene, transcriptional repressor for mreB; also regulator for dacA, dacC, and ampC | 3.849887 | 0.003751 | 0.177985 | 3767.9 | 978.54 |
| bssS | ECD_01056 | biofilm regulator | 2.773061 | 0.025726 | 0.477901 | 8929.174 | 3219.828 |
| chbC | ECD_01706 | N,N'-diacetylchitobiose-specific enzyme IIC component of PTS | 4.538809 | 0.007841 | 0.298191 | 71.54241 | 15.58949 |
| citX | ECD_00582 | apo-citrate lyase phosphoribosyl-dephospho-CoA transferase | 13.08699 | 0.000935 | 0.072227 | 34.06781 | 2.398382 |
| creC | ECD_04275 | sensory histidine kinase in two-component regulatory system with CreB or PhoB | 3.46148 | 0.032593 | 0.519559 | 54.5085 | 15.58949 |
| cspC | ECD_01793 | stress protein, member of the CspA-family | 5.556331 | 0.000546 | 0.0518 | 347.4917 | 62.35794 |
| cspE | ECD_00593 | constitutive cold shock family transcription antitermination protein; negative regulator of cspA transcription; RNA melting protein; ssDNA-binding protein | 7.54401 | 4.64E-05 | 0.008151 | 616.6274 | 81.545 |
| cutC | ECD_01845 | copper homeostasis protein | 6.964267 | 6.17E-05 | 0.009852 | 1529.645 | 219.452 |

| | | | | | | | |
|---|---|---|---|---|---|---|---|
| cynX | ECD_00295 | putative cyanate transporter | 9.186281 | 0.00679 | 0.270888 | 23.84747 | 2.398382 |
| ddpC | ECD_01443 | D,D-dipeptide ABC transporter permease | 8.085941 | 0.004943 | 0.211665 | 30.66103 | 3.597574 |
| diaA | ECD_03016 | DnaA initiator-associating factor for replication initiation | 3.495332 | 0.019935 | 0.442983 | 88.57632 | 25.18302 |
| dtpC | ECD_04001 | dipeptide and tripeptide permease | 3.676945 | 0.024268 | 0.46674 | 57.91528 | 15.58949 |
| dusB | ECD_03118 | tRNA-dihydrouridine synthase B | 5.506834 | 0.00128 | 0.086421 | 139.678 | 25.18302 |
| ecpA | ECD_00252 | ECP pilin | 31.32438 | 3.33E-05 | 0.007055 | 44.28816 | 1.199191 |
| elyC | ECD_00924 | envelope biogenesis factor; DUF218 superfamily protein | 3.134626 | 0.039049 | 0.561312 | 68.13563 | 21.58544 |
| eno | ECD_02624 | enolase | 2.523128 | 0.042581 | 0.582241 | 4466.29 | 1770.006 |
| fabF | ECD_01091 | 3-oxoacyl-[acyl-carrier-protein | 3.019607 | 0.016262 | 0.404932 | 3849.663 | 1274.74 |
| fabZ | ECD_00178 | (3R)-hydroxymyristol acyl carrier protein dehydratase | 3.161011 | 0.019625 | 0.43887 | 197.5933 | 62.35794 |
| fdhF | ECD_03951 | formate dehydrogenase-H, selenopolypeptide subunit | 2.570259 | 0.041695 | 0.574325 | 780.1529 | 303.3954 |
| fdrA | ECD_00468 | putative NAD(P)-binding acyl-CoA synthetase | 10.48652 | 0.003397 | 0.167941 | 27.25425 | 2.398382 |
| fetB | ECD_00442 | iron export ABC transporter permease; peroxide resistance protein | 4.323343 | 0.010321 | 0.320887 | 68.13563 | 15.58949 |
| fhuA | ECD_00149 | ferrichrome outer membrane transporter | 3.385106 | 0.009279 | 0.308725 | 844.8818 | 249.4318 |
| frdD | ECD_04023 | fumarate reductase (anaerobic), membrane anchor subunit | 4.378659 | 0.003917 | 0.183385 | 163.5255 | 37.17493 |
| frlD | ECD_03224 | fructoselysine 6-kinase | 4.350342 | 0.016253 | 0.404932 | 47.69494 | 10.79272 |
| frwD | ECD_03838 | putative enzyme IIB component of PTS | 6.585809 | 0.029994 | 0.518486 | 17.03391 | 2.398382 |
| ftsB | ECD_02598 | cell division protein | 2.929617 | 0.039821 | 0.561491 | 98.79666 | 33.57735 |
| fucI | ECD_02653 | L-fucose isomerase | 3.423418 | 0.013042 | 0.361367 | 197.5933 | 57.56118 |
| gadX | ECD_03364 | acid resistance regulon transcriptional activator; autoactivator | 3.434245 | 0.009939 | 0.317234 | 391.7799 | 113.9232 |
| gfcC | ECD_00988 | putative O-antigen capsule production periplasmic protein | 6.585809 | 0.029994 | 0.518486 | 17.03391 | 2.398382 |
| glgS | ECD_02919 | motility and biofilm regulator | 4.199755 | 0.002111 | 0.119535 | 4578.714 | 1090.065 |
| glnB | ECD_02445 | regulatory protein P-II for glutamine synthetase | 4.187669 | 0.02311 | 0.464072 | 40.88138 | 9.59353 |
| glvBC | ECD_03566 | arbutin specific enzyme IIBC component of PTS | 4.966913 | 0.020774 | 0.453793 | 30.66103 | 5.995956 |
| grcA | ECD_02473 | autonomous glycyl radical cofactor | 3.77471 | 0.008271 | 0.305695 | 177.1526 | 46.76846 |

| | | | | | | | |
|---|---|---|---|---|---|---|---|
| greB | ECD_03258 | transcript cleavage factor | 7.706497 | 0.001895 | 0.110868 | 47.69494 | 5.995956 |
| gspH | ECD_03180 | putative general secretory pathway component, cryptic | 62.45439 | 0.002355 | 0.127194 | 13.62713 | 0 |
| hns | ECD_01213 | global DNA-binding transcriptional dual regulator H-NS | 7.787963 | 2.09E-05 | 0.005234 | 14822.91 | 1903.116 |
| hofB | ECD_00106 | T2SE secretion family protein; P-loop ATPase superfamily protein | 3.192656 | 0.041713 | 0.574325 | 57.91528 | 17.98787 |
| hokD | ECD_01532 | Qin prophage; small toxic polypeptide | 833.1739 | 2.46E-21 | 8.63E-18 | 3181.934 | 3.597574 |
| hpf | ECD_03068 | ribosome hibernation promoting factor HPF; stabilizes 100S dimers | 3.038436 | 0.021803 | 0.456012 | 255.5086 | 83.94338 |
| hspQ | ECD_00970 | heat shock protein involved in degradation of mutant DnaA; hemimethylated oriC DNA-binding protein | 2.819931 | 0.024334 | 0.46674 | 1887.357 | 669.1487 |
| hupB | ECD_00392 | HU, DNA-binding transcriptional regulator, beta subunit | 8.143978 | 2.46E-05 | 0.005762 | 694.9834 | 85.14257 |
| hybD | ECD_02869 | maturation protease for hydrogenase 2 | 3.668497 | 0.01606 | 0.404932 | 88.57632 | 23.98382 |
| hyfG | ECD_02379 | hydrogenase 4, subunit | 8.62491 | 9.06E-05 | 0.013824 | 146.4916 | 16.78868 |
| ibaG | ECD_03055 | acid stress protein; putative BolA family transcriptional regulator | 6.585809 | 0.029994 | 0.518486 | 17.03391 | 2.398382 |
| ibsD | ECD_04363 | toxic membrane protein | 5.40998 | 0.033148 | 0.519559 | 20.44069 | 3.597574 |
| insA-20 | ECD_02513 | IS1 protein InsA | 5.40998 | 0.033148 | 0.519559 | 20.44069 | 3.597574 |
| insF-2 | ECD_01360 | IS3 element protein InsF | 17.00581 | 3.62E-05 | 0.007055 | 64.72885 | 3.597574 |
| iscX | ECD_02416 | Fe(2+) donor and activity modulator for cysteine desulfurase | 3.46148 | 0.032593 | 0.519559 | 54.5085 | 15.58949 |
| lpp | ECD_01646 | murein lipoprotein | 20.37404 | 4.48E-09 | 3.94E-06 | 29567.46 | 1451.021 |
| lpxA | ECD_00179 | UDP-N-acetylglucosamine acetyltransferase | 5.159657 | 0.001008 | 0.072227 | 279.3561 | 53.9636 |
| lpxB | ECD_00180 | tetraacyldisaccharide-1-P synthase | 3.344291 | 0.018232 | 0.43887 | 132.8645 | 39.57331 |
| lpxC | ECD_00097 | UDP-3-O-acyl N-acetylglucosamine deacetylase | 6.350676 | 0.000112 | 0.015761 | 3192.154 | 502.4611 |
| lpxD | ECD_00177 | UDP-3-O-(3-hydroxymyristoyl)-glucosamine N-acyltransferase | 3.271445 | 0.013084 | 0.361367 | 384.9663 | 117.5207 |
| lspA | ECD_00031 | prolipoprotein signal peptidase (signal peptidase II) | 4.139157 | 0.006016 | 0.251459 | 139.678 | 33.57735 |

| | | | | | | | |
|---|---|---|---|---|---|---|---|
| marB | ECD_01491 | periplasmic mar operon regulator | 12.14387 | 0.009409 | 0.308725 | 17.03391 | 1.199191 |
| mcrC | ECD_04211 | 5-methylcytosine-specific restriction enzyme McrBC, subunit McrC | 16.939 | 0.001795 | 0.106822 | 23.84747 | 1.199191 |
| mepS | ECD_02105 | murein DD-endopeptidase, space-maker hydrolase, mutational suppressor of prc thermosensitivity, outer membrane lipoprotein, weak murein LD-carboxypeptidase | 3.217454 | 0.042785 | 0.582241 | 54.5085 | 16.78868 |
| metJ | ECD_03824 | transcriptional repressor, S-adenosylmethionine-binding | 3.84058 | 0.034643 | 0.520846 | 37.4746 | 9.59353 |
| mlaA | ECD_02272 | ABC transporter maintaining OM lipid asymmetry, OM lipoprotein component | 2.742079 | 0.031826 | 0.519559 | 483.763 | 176.2811 |
| mltD | ECD_00204 | putative membrane-bound lytic murein transglycosylase D | 6.158456 | 0.000168 | 0.02059 | 1086.763 | 176.2811 |
| mokB | ECD_01377 | regulatory peptide | 3.337777 | 0.024422 | 0.46674 | 88.57632 | 26.38221 |
| mokC | ECD_00017 | regulatory protein for HokC | 62.45439 | 0.002355 | 0.127194 | 13.62713 | 0 |
| mreC | ECD_03109 | cell wall structural complex MreBCD transmembrane component MreC | 3.16955 | 0.021326 | 0.453793 | 160.1187 | 50.36603 |
| mreD | ECD_03108 | cell wall structural complex MreBCD transmembrane component MreD | 4.623171 | 0.02182 | 0.456012 | 34.06781 | 7.195147 |
| nadB | ECD_02468 | quinolinate synthase, L-aspartate oxidase (B protein) subunit | 2.9327 | 0.028079 | 0.510807 | 204.4069 | 69.55309 |
| nagC | ECD_00633 | N-acetylglucosamine-inducible nag divergent operon transcriptional repressor | 2.972479 | 0.025544 | 0.477041 | 221.4408 | 74.34985 |
| nrdH | ECD_02529 | hydrogen donor for NrdEF electron transport system; glutaredoxin-like protein | 4.796089 | 0.034567 | 0.520846 | 23.84747 | 4.796765 |
| osmE | ECD_01708 | osmotically-inducible lipoprotein | 8.729785 | 1.06E-05 | 0.003721 | 1509.204 | 172.6835 |
| pcnB | ECD_00142 | poly(A) polymerase | 2.971236 | 0.048139 | 0.640216 | 68.13563 | 22.78463 |
| pdhR | ECD_00112 | pyruvate dehydrogenase complex repressor; autorepressor | 2.719794 | 0.038463 | 0.555741 | 238.4747 | 87.54096 |
| pgaB | ECD_01025 | poly-beta-1,6-N-acetyl-D-glucosamine (PGA) N-deacetylase outer membrane export lipoprotein | 3.84058 | 0.034643 | 0.520846 | 37.4746 | 9.59353 |

| | | | | | | | |
|---|---|---|---|---|---|---|---|
| pgrR | ECD_01306 | murein peptide degradation regulator | 3.318453 | 0.046767 | 0.624324 | 44.28816 | 13.1911 |
| pitB | ECD_02863 | phosphate transporter | 5.51483 | 0.012477 | 0.353285 | 34.06781 | 5.995956 |
| pnp | ECD_03031 | polynucleotide phosphorylase/polyadenylase | 4.463405 | 0.001525 | 0.099178 | 1451.289 | 324.9808 |
| priB | ECD_04068 | primosomal protein N | 7.538298 | 0.000998 | 0.072227 | 64.72885 | 8.394338 |
| purA | ECD_04044 | adenylosuccinate synthetase | 2.748753 | 0.034027 | 0.520846 | 316.8307 | 115.1224 |
| rhsB | ECD_03331 | Rhs protein with DUF4329 family putative toxin domain; putative neighboring cell growth inhibitor | 3.246389 | 0.024304 | 0.46674 | 105.6102 | 32.37816 |
| rmf | ECD_00957 | ribosome modulation factor | 51.17602 | 8.01E-13 | 1.41E-09 | 11978.24 | 233.8423 |
| rof | ECD_00187 | modulator of Rho-dependent transcription termination | 2.987261 | 0.019071 | 0.43887 | 756.3055 | 253.0293 |
| rplA | ECD_03860 | 50S ribosomal subunit protein L1 | 2.731246 | 0.030338 | 0.518486 | 858.5089 | 314.1881 |
| rplI | ECD_04070 | 50S ribosomal subunit protein L9 | 5.732205 | 0.000532 | 0.0518 | 269.1357 | 46.76846 |
| rplK | ECD_03859 | 50S ribosomal subunit protein L11 | 2.582293 | 0.04626 | 0.619923 | 310.0171 | 119.9191 |
| rplQ | ECD_03145 | 50S ribosomal subunit protein L17 | 4.581168 | 0.001313 | 0.087007 | 1209.407 | 263.8221 |
| rplT | ECD_01685 | 50S ribosomal subunit protein L20 | 4.593493 | 0.001203 | 0.082835 | 2204.188 | 479.6765 |
| rpmA | ECD_03050 | 50S ribosomal subunit protein L27 | 2.687881 | 0.035911 | 0.529768 | 425.8477 | 158.2932 |
| rpmB | ECD_03494 | 50S ribosomal subunit protein L28 | 5.541826 | 0.00872 | 0.308177 | 40.88138 | 7.195147 |
| rpmG | ECD_03493 | 50S ribosomal subunit protein L33 | 8.80233 | 0.000807 | 0.070868 | 54.5085 | 5.995956 |
| rpmI | ECD_01686 | 50S ribosomal subunit protein L35 | 4.446064 | 0.003241 | 0.16257 | 187.373 | 41.97169 |
| rpoA | ECD_03146 | RNA polymerase, alpha subunit | 2.493898 | 0.045829 | 0.616501 | 2255.289 | 904.1902 |
| rpoE | ECD_02467 | RNA polymerase sigma E factor | 3.447987 | 0.00876 | 0.308177 | 616.6274 | 178.6795 |
| rpsB | ECD_00167 | 30S ribosomal subunit protein S2 | 3.134611 | 0.013402 | 0.364757 | 2176.933 | 694.3317 |
| rpsF | ECD_04067 | 30S ribosomal subunit protein S6 | 4.51296 | 0.004869 | 0.211035 | 109.017 | 23.98382 |
| rpsK | ECD_03148 | 30S ribosomal subunit protein S11 | 2.499139 | 0.0493 | 0.65072 | 575.7461 | 230.2447 |
| rpsO | ECD_03032 | 30S ribosomal subunit protein S15 | 3.409615 | 0.008777 | 0.308177 | 936.8649 | 274.6148 |
| rpsQ | ECD_03162 | 30S ribosomal subunit protein S17 | 3.992406 | 0.004247 | 0.196216 | 364.5256 | 91.13853 |
| rpsR | ECD_04069 | 30S ribosomal subunit protein S18 | 12.14387 | 0.009409 | 0.308725 | 17.03391 | 1.199191 |

| | | | | | | | |
|---|---|---|---|---|---|---|---|
| rpsT | ECD_00027 | 30S ribosomal subunit protein S20 | 6.177334 | 0.000578 | 0.053393 | 156.7119 | 25.18302 |
| rpsU | ECD_02935 | 30S ribosomal subunit protein S21 | 13.34723 | 4.18E-07 | 0.000245 | 947.0852 | 70.75228 |
| rraA | ECD_03814 | ribonuclease E (RNase E) inhibitor protein | 3.913945 | 0.003444 | 0.167941 | 2483.544 | 634.3721 |
| rsmE | ECD_02776 | 16S rRNA m(3)U1498 methyltransferase, SAM-dependent | 4.760002 | 0.008624 | 0.308177 | 57.91528 | 11.99191 |
| rspB | ECD_01549 | putative Zn-dependent NAD(P)-binding oxidoreductase | 3.044887 | 0.034497 | 0.520846 | 95.38988 | 31.17897 |
| ryfB | ECD_04346 | hypothetical protein | 6.352104 | 0.002965 | 0.153499 | 54.5085 | 8.394338 |
| rzpD | ECD_00508 | DLP12 prophage; putative murein endopeptidase | 4.418997 | 0.035098 | 0.520846 | 27.25425 | 5.995956 |
| secE | ECD_03857 | preprotein translocase membrane subunit | 3.01071 | 0.026356 | 0.487029 | 173.7459 | 57.56118 |
| secG | ECD_03040 | preprotein translocase membrane subunit | 8.602427 | 6.06E-05 | 0.009852 | 187.373 | 21.58544 |
| sieB | ECD_01331 | phage superinfection exclusion protein, Rac prophage | 3.941004 | 0.014502 | 0.379968 | 71.54241 | 17.98787 |
| skp | ECD_00176 | periplasmic chaperone | 3.276121 | 0.012614 | 0.354298 | 432.6612 | 131.911 |
| ssb | ECD_03931 | single-stranded DNA-binding protein | 4.722417 | 0.001074 | 0.075438 | 1178.746 | 249.4318 |
| symE | ECD_04213 | toxic peptide regulated by antisense sRNA symR | 5.51483 | 0.012477 | 0.353285 | 34.06781 | 5.995956 |
| tdcA | ECD_02985 | tdc operon transcriptional activator | 7.886045 | 0.014032 | 0.370415 | 20.44069 | 2.398382 |
| torY | ECD_01844 | TMAO reductase III (TorYZ), cytochrome c-type subunit | 3.415476 | 0.02446 | 0.46674 | 78.35597 | 22.78463 |
| trxA | ECD_03659 | thioredoxin 1 | 6.090052 | 0.000161 | 0.02059 | 2067.916 | 339.3711 |
| uxuB | ECD_04192 | D-mannonate oxidoreductase, NAD-dependent | 2.959111 | 0.029658 | 0.518486 | 160.1187 | 53.9636 |
| vioB | ECD_01941 | VioB, involved in dTDP-N-acetylviosamine synthesis | 3.51241 | 0.032995 | 0.519559 | 51.10172 | 14.39029 |
| waaV | ECD_03480 | putative beta1,3-glucosyltransferase | 3.267396 | 0.030569 | 0.518486 | 74.94919 | 22.78463 |
| wcaD | ECD_01962 | putative colanic acid polymerase | 17.00581 | 3.62E-05 | 0.007055 | 64.72885 | 3.597574 |
| xthA | ECD_01718 | exonuclease III | 5.962049 | 0.000976 | 0.072227 | 122.6441 | 20.38625 |
| xylE | ECD_03903 | D-xylose transporter | 2.712545 | 0.036868 | 0.541356 | 296.39 | 109.1264 |
| yaaW | ECD_00011 | UPF0174 family protein | 4.187669 | 0.02311 | 0.464072 | 40.88138 | 9.59353 |
| yadV | ECD_00139 | putative periplasmic pilin chaperone | 7.886045 | 0.014032 | 0.370415 | 20.44069 | 2.398382 |
| ybcJ | ECD_00478 | ribosome-associated protein; putative RNA-binding protein | 15.29454 | 3.15E-06 | 0.00123 | 149.8984 | 9.59353 |
| ybcK | ECD_00494 | DLP12 prophage; putative recombinase | 6.206146 | 0.002108 | 0.119535 | 68.13563 | 10.79272 |

| | | | | | | | |
|---|---|---|---|---|---|---|---|
| ybgF | ECD_00702 | periplasmic TolA-binding protein | 2.643229 | 0.0376 | 0.545683 | 551.8986 | 208.6593 |
| ybjS | ECD_00873 | putative NAD(P)H-dependent oxidoreductase | 2.741351 | 0.030532 | 0.518486 | 667.7292 | 243.4358 |
| yceD | ECD_01084 | DUF177 family protein | 3.246461 | 0.021017 | 0.453793 | 132.8645 | 40.7725 |
| ycgI | ECD_01148 | hypothetical protein | 9.746308 | 0.023263 | 0.464072 | 13.62713 | 1.199191 |
| ycgX | ECD_01136 | DUF1398 family protein | 5.16591 | 0.009699 | 0.315305 | 44.28816 | 8.394338 |
| yciC | ECD_01229 | UPF0259 family inner membrane protein | 2.94429 | 0.04277 | 0.582241 | 85.16954 | 28.78059 |
| yciY | ECD_04308 | uncharacterized protein | 42.77316 | 1.42E-09 | 1.66E-06 | 265.7289 | 5.995956 |
| ydcC | ECD_01418 | H repeat-associated putative transposase | 5.40998 | 0.033148 | 0.519559 | 20.44069 | 3.597574 |
| ydcH | ECD_01383 | DUF465 family protein | 2.735667 | 0.029241 | 0.518486 | 1250.289 | 456.8918 |
| ydeQ | ECD_01460 | putative fimbrial-like adhesin protein | 62.45439 | 0.002355 | 0.127194 | 13.62713 | 0 |
| ydfK | ECD_01503 | cold shock protein, function unknown, Qin prophage | 3.27404 | 0.011718 | 0.352687 | 640.4749 | 195.4682 |
| ydhZ | ECD_01644 | uncharacterized protein | 3.093031 | 0.037005 | 0.541356 | 78.35597 | 25.18302 |
| ydjH | ECD_01741 | putative kinase | 9.746308 | 0.023263 | 0.464072 | 13.62713 | 1.199191 |
| yebO | ECD_01795 | putative inner membrane protein | 2.834947 | 0.033672 | 0.520846 | 194.1865 | 68.3539 |
| yecH | ECD_01874 | DUF2492 family protein | 3.603295 | 0.019449 | 0.43887 | 78.35597 | 21.58544 |
| yecT | ECD_01848 | uncharacterized protein | 3.979716 | 0.034967 | 0.520846 | 34.06781 | 8.394338 |
| yegK | ECD_01977 | ser/thr phosphatase-related protein | 4.534759 | 0.015553 | 0.395689 | 44.28816 | 9.59353 |
| yfbR | ECD_02216 | 5'-nucleotidase | 4.163844 | 0.035158 | 0.520846 | 30.66103 | 7.195147 |
| yfcJ | ECD_02247 | putative arabinose efflux transporter | 10.33865 | 4.40E-05 | 0.008132 | 126.0509 | 11.99191 |
| yfcZ | ECD_02267 | UPF0381 family protein | 14.93671 | 1.38E-06 | 0.000604 | 218.034 | 14.39029 |
| yfeD | ECD_02305 | DUF1323 family putative DNA-binding protein | 2.764708 | 0.028824 | 0.518242 | 756.3055 | 273.4156 |
| ygbA | ECD_02582 | uncharacterized protein | 3.316346 | 0.024402 | 0.46674 | 91.9831 | 27.5814 |
| ygdR | ECD_02681 | DUF903 family verified lipoprotein | 22.50025 | 5.14E-07 | 0.000258 | 139.678 | 5.995956 |
| ygfK | ECD_02711 | putative Fe-S subunit oxidoreductase subunit | 7.191291 | 0.000108 | 0.015761 | 303.2035 | 41.97169 |
| yggD | ECD_02760 | MtlR family putative transcriptional repressor | 3.561411 | 0.016737 | 0.41094 | 98.79666 | 27.5814 |
| yggI | ECD_02774 | Zn-dependent metalloprotease-related protein | 4.375114 | 0.022569 | 0.464072 | 37.4746 | 8.394338 |

| | | | | | | | |
|---|---|---|---|---|---|---|---|
| yggS | ECD_02781 | UPF0001 family protein, PLP-binding | 3.818282 | 0.018485 | 0.43887 | 64.72885 | 16.78868 |
| yghR | ECD_02860 | putative ATP-binding protein | 93.18158 | 0.000319 | 0.033909 | 20.44069 | 0 |
| yghS | ECD_02861 | putative ATP-binding protein | 16.939 | 0.001795 | 0.106822 | 23.84747 | 1.199191 |
| ygiN | ECD_02901 | quinol monooxygenase | 3.899535 | 0.004836 | 0.211035 | 374.746 | 95.9353 |
| ygiZ | ECD_02899 | inner membrane protein | 7.538298 | 0.000998 | 0.072227 | 64.72885 | 8.394338 |
| ygjI | ECD_02947 | putative transporter | 7.193954 | 0.009097 | 0.308725 | 27.25425 | 3.597574 |
| yhaB | ECD_02987 | uncharacterized protein | 4.418997 | 0.035098 | 0.520846 | 27.25425 | 5.995956 |
| yhfX | ECD_03233 | putative pyridoxal 5'-phosphate binding protein | 9.746308 | 0.023263 | 0.464072 | 13.62713 | 1.199191 |
| yhiKL | ECD_03339 | hypothetical protein | 7.379136 | 0.001624 | 0.103693 | 54.5085 | 7.195147 |
| yiaB | ECD_03415 | YiaAB family inner membrane protein | 9.746308 | 0.023263 | 0.464072 | 13.62713 | 1.199191 |
| yifE | ECD_03643 | UPF0438 family protein | 5.89942 | 0.000473 | 0.048798 | 248.695 | 41.97169 |
| yihD | ECD_03744 | DUF1040 protein YihD | 4.968943 | 0.007961 | 0.298191 | 54.5085 | 10.79272 |
| yihL | ECD_03757 | putative DNA-binding transcriptional regulator | 11.78675 | 0.001755 | 0.106822 | 30.66103 | 2.398382 |
| yjbE | ECD_03898 | extracellular polysaccharide production threonine-rich protein | 16.939 | 0.001795 | 0.106822 | 23.84747 | 1.199191 |
| yjbJ | ECD_03917 | stress-induced protein, UPF0337 family | 3.500016 | 0.006655 | 0.268586 | 12462.01 | 3560.399 |
| yjbL | ECD_03919 | uncharacterized protein | 12.14387 | 0.009409 | 0.308725 | 17.03391 | 1.199191 |
| yjhX | ECD_04174 | UPF0386 family protein | 13.08699 | 0.000935 | 0.072227 | 34.06781 | 2.398382 |
| yjjV | ECD_04254 | putative DNase | 3.246266 | 0.030295 | 0.518486 | 78.35597 | 23.98382 |
| ykiA | ECD_00339 | hypothetical protein | 4.619387 | 0.006433 | 0.262646 | 78.35597 | 16.78868 |
| ymdF | ECD_04303 | KGG family protein | 4.678155 | 0.007337 | 0.289456 | 68.13563 | 14.39029 |
| ymdF | ECD_01008 | KGG family protein | 3.863167 | 0.003654 | 0.17575 | 4221.002 | 1092.463 |
| ynaE | ECD_01346 | cold shock protein, Rac prophage | 3.27404 | 0.011718 | 0.352687 | 640.4749 | 195.4682 |
| ynaJ | ECD_01309 | DUF2534 family putative inner membrane protein | 8.532225 | 1.26E-05 | 0.004025 | 1556.899 | 182.2771 |
| yneL | ECD_01468 | putative transcriptional regulator | 77.81799 | 0.000833 | 0.071296 | 17.03391 | 0 |
| yniA | ECD_01694 | fructosamine kinase family protein | 3.659973 | 0.005595 | 0.236674 | 1154.899 | 315.3873 |
| ynjD | ECD_01725 | putative ABC transporter ATPase | 10.44608 | 0.000248 | 0.028998 | 64.72885 | 5.995956 |

| | | | | | | | |
|---|---|---|---|---|---|---|---|
| ynjI | ECD_01731 | inner membrane protein | 13.08699 | 0.000935 | 0.072227 | 34.06781 | 2.398382 |
| yqfA | ECD_02731 | hemolysin III family HylIII inner membrane protein | 2.615779 | 0.03955 | 0.561312 | 558.7122 | 213.456 |
| yqiA | ECD_02903 | acyl CoA esterase | 3.46148 | 0.032593 | 0.519559 | 54.5085 | 15.58949 |
| yqiB | ECD_02905 | DUF1249 protein YqiB | 3.447072 | 0.02446 | 0.46674 | 74.94919 | 21.58544 |
| yraH | ECD_03009 | putative fimbrial-like adhesin protein | 4.406606 | 0.004523 | 0.20261 | 132.8645 | 29.97978 |
| yraJ | ECD_03011 | putative outer membrane protein | 13.59433 | 3.22E-07 | 0.000226 | 1127.645 | 82.74419 |
| yrbL | ECD_03072 | Mg(2+)-starvation-stimulated protein | 3.052986 | 0.034929 | 0.520846 | 91.9831 | 29.97978 |
| ytfK | ECD_04088 | DUF1107 family protein | 3.931535 | 0.00317 | 0.161301 | 22201.99 | 5646.991 |
| ytfP | ECD_04093 | GGCT-like protein | 3.818045 | 0.004559 | 0.20261 | 783.5597 | 205.0617 |
| yzfA | ECD_04094 | hypothetical protein | 16.98769 | 0.000167 | 0.02059 | 44.28816 | 2.398382 |
| zapA | ECD_02742 | FtsZ stabilizer | 8.311171 | 1.39E-05 | 0.004065 | 2742.459 | 329.7776 |
| | ECD_03459 | hypothetical protein | 93.18158 | 0.000319 | 0.033909 | 20.44069 | 0 |
| | ECD_00022 | hypothetical protein | 12.14387 | 0.009409 | 0.308725 | 17.03391 | 1.199191 |
| | ECD_00815 | integrase for prophage | 5.730951 | 0.000523 | 0.0518 | 275.9493 | 47.96765 |
| | ECD_02652 | hypothetical protein | 5.51483 | 0.012477 | 0.353285 | 34.06781 | 5.995956 |
| | ECD_04314 | hypothetical protein | 4.474227 | 0.006136 | 0.253473 | 91.9831 | 20.38625 |
| | ECD_03695 | Magnesium and cobalt transport protein corA | 3.84058 | 0.034643 | 0.520846 | 37.4746 | 9.59353 |
| | ECD_02621 | hypothetical protein | 3.595737 | 0.011338 | 0.349194 | 160.1187 | 44.37007 |
| | ECD_00840 | hypothetical protein | 3.379743 | 0.031841 | 0.519559 | 61.32207 | 17.98787 |
| | ECD_02855 | hypothetical protein | 3.113105 | 0.013798 | 0.370415 | 2800.374 | 899.3934 |
| | ECD_03786 | putative glycoporin | 3.085895 | 0.02023 | 0.446718 | 252.1018 | 81.545 |

**Table S7. Bacterial strains used in this study.**

| Strain | Relevant characteristic(s) | Reference or source |
|---|---|---|
| *B. coagulans* 2-6 | Wild type | Ref. 22 |
| *E. coli* DH5α | *supE44ΔlacU169* (*Φ80 lacZΔM15*) *hsdR17 recA1 endA1 gyrA96 thi-1 relA1* | Novagen |
| *E. coli* BL21 (DE3) | *F- ompT hsdSB (rB-mB-) gal (λ c I 857 ind1 Sam7 nin5 lacUV5 T7gene1) dcm (DE3)* | Novagen |
| *P. putida* KT2440 | *rmo- mod+* | ATCC |
| *S. cerevisiae* INVSc1 | *MATa his3D1 leu2 trp1-289 ura3-52 MAT his3D1 leu2 trp1-289 ura3-52* | ATCC |
| ATCC31280::pLQ856 | wild-type harboring pLQ856 | Ref. 31 |
| ATCC31280::pLQ856-cspL | wild-type harboring pLQ856-*cspL* | This study |
| DH5α | *E. coli* DH5α harboring pUC19 empty vector | This study |
| DH5α-*cspL* | *E. coli* DH5α harboring pUC19-*cspL* | This study |
| DH5α-*cspD* | *E. coli* DH5α harboring pUC19-*cspD* | This study |
| DH5α-*BCO26_YkuS* | *E. coli* DH5α harboring pUC19-*BCO26_ykuS* | This study |
| DH5α-*BCO26_2915* | *E. coli* DH5α harboring pUC19*BCO26_2915* | This study |
| DH5α-*BCO26_Dps* | *E. coli* DH5α harboring pUC19*BCO26_dps* | This study |
| DH5α-*BCO26_GsiB* | *E. coli* DH5α harboring pUC19*BCO26_gsiB* | This study |
| DH5α-*BCO26_YbfB* | *E. coli* DH5α harboring pUC19*BCO26_ybfB* | This study |
| DH5α-*BCO26_MntH* | *E. coli* DH5α harboring pUC19*BCO26_mntH* | This study |
| DH5α-*BCO26_2932* | *E. coli* DH5α harboring pUC19*BCO26_2932* | This study |
| DH5α-*BCO26_GabD* | *E. coli* DH5α harboring pUC19*BCO26_gabD* | This study |
| DH5α-*BCO26_GroEL* | *E. coli* DH5α harboring pUC19*BCO26_groEL* | This study |
| DH5α-*BCO26_HrcA* | *E. coli* DH5α harboring pUC19*BCO26_hrcA* | This study |
| DH5α-*BCO26_ClpE* | *E. coli* DH5α harboring pUC19*BCO26_clpE* | This study |

| | | |
|---|---|---|
| DH5α-*BCO26_1771* | *E. coli* DH5α harboring pUC19*BCO26_1771* | This study |
| DH5α-*BCO26_DnaK* | *E. coli* DH5α harboring pUC19*BCO26_dnaK* | This study |
| DH5α-*BCO26_GroES* | *E. coli* DH5α harboring pUC19*BCO26_groES* | This study |
| DH5α-*BCO26_0541* | *E. coli* DH5α harboring pUC19*BCO26_0541* | This study |
| DH5α-*BCO26_YkzI* | *E. coli* DH5α harboring pUC19*BCO26_ykzI* | This study |
| DH5α-*BCO26_KatE* | *E. coli* DH5α harboring pUC19*BCO26_katE* | This study |
| DH5α-*BCO26_0399* | *E. coli* DH5α harboring pUC19*BCO26_0399* | This study |
| DH5α-*BCO26_2461* | *E. coli* DH5α harboring pUC19*BCO26_2461* | This study |
| DH5α-*BCO26_LevG* | *E. coli* DH5α harboring pUC19*BCO26_levG* | This study |
| DH5α-*BCO26_YqiG* | *E. coli* DH5α harboring pUC19*BCO26_yqiG* | This study |
| DH5α-*BCO26_2340* | *E. coli* DH5α harboring pUC19*BCO26_2340* | This study |
| DH5α-*BCO26_YflT* | *E. coli* DH5α harboring pUC19*BCO26_yflT* | This study |
| DH5α-*BCO26_ArgI* | *E. coli* DH5α harboring pUC19*BCO26_argL* | This study |
| DH5α-*BCO26_SucC* | *E. coli* DH5α harboring pUC19*BCO26_sucC* | This study |
| DH5α-*BCO26_2370* | *E. coli* DH5α harboring pUC19*BCO26_2370* | This study |
| DH5α-*BCO26_CitZ* | *E. coli* DH5α harboring pUC19*BCO26_citZ* | This study |
| DH5α-*BCO26_RsbV* | *E. coli* DH5α harboring pUC19*BCO26_rsbV* | This study |
| DH5α-*BCO26_Hag* | *E. coli* DH5α harboring pUC19*BCO26_rhag* | This study |
| DH5α-*BCO26_YhgD* | *E. coli* DH5α harboring pUC19*BCO26_yhgD* | This study |
| DH5α-*BCO26_1679* | *E. coli* DH5α harboring pUC19*BCO26_1679* | This study |
| DH5α-*BCO26_RsbW* | *E. coli* DH5α harboring pUC19*BCO26_rsbW* | This study |
| DH5α-*BCO26_YteA* | *E. coli* DH5α harboring pUC19*BCO26_yteA* | This study |
| DH5α-*BCO26_2825* | *E. coli* DH5α harboring pUC19*BCO26_2825* | This study |
| DH5α-*BCO26_LevE* | *E. coli* DH5α harboring pUC19*BCO26_levE* | This study |
| DH5α-*BCO26_SigB* | *E. coli* DH5α harboring pUC19*BCO26_sigB* | This study |

| | | |
|---|---|---|
| DH5α-*BCO26_2573* | *E. coli* DH5α harboring pUC19*BCO26_2573* | This study |
| DH5α-*BCO26_1484* | *E. coli* DH5α harboring pUC19*BCO26_1484* | This study |
| BL21-*cspL* | *E. coli* BL21(DE3) harboring pET28a-*cspL* | This study |
| BL21-*cspL*-M11 | *E. coli* BL21(DE3) harboring pET28a-*cspL* with mutated G14 Y15 G16 F17 I18 E19 R20 V26 F27 V28 H29 | This study |
| BL21-*cspL*-M7 | *E. coli* BL21(DE3) harboring pET28a-*cspL* with mutated G14 Y15 G16 F17 I18 E19 R20 | This study |
| INVSc1 | *S. cerevisiae* INVSc1 harboring pYES2 empty vector | This study |
| INVSc1-*cspL* | *S. cerevisiae* INVSc1 harboring pYES2-*cspL* | This study |
| KT2440 | *P. putida* KT2440 harboring pME6032 empty vector | This study |
| KT2440-*cspL* | *P. putida* KT2440 harboring pME6032-*cspL* | This study |
| DH5α-*cspA* | *E. coli* DH5α harboring pUC19-*cspA* | This study |

**Table S8. Plasmids used in this study.**

| Plasmid | Relevant characteristic(s)[a] | Reference or source |
|---|---|---|
| pUC19 | Amp$^R$, *pMB1 ori*, P*lacZ* | Novagen |
| pET28a(+) | Kan$^R$, *pBR322 ori*, PT7 | Novagen |
| pUC19-*cspL* | pUC19 harboring *cspL* from *B. coagulans 2-6* | This study |
| pUC19-*BCO26_cspD* | pUC19 harboring *BCO26_cspD* from *B. coagulans 2-6* | This study |
| pUC19-*BCO26_ykuS* | pUC19 harboring *BCO26_ykuS* from *B. coagulans 2-6* | This study |
| pUC19-*BCO26_2915* | pUC19 harboring *BCO26_2915* from *B. coagulans 2-6* | This study |
| pUC19-*BCO26_dps* | pUC19 harboring *BCO26_dps* from *B. coagulans 2-6* | This study |
| pUC19-*BCO26_gsiB* | pUC19 harboring *BCO26_gsiB* from *B. coagulans 2-6* | This study |
| pUC19-*BCO26_ybfB* | pUC19 harboring *BCO26_ybfB* from *B. coagulans 2-6* | This study |
| pUC19-*BCO26_mntH* | pUC19 harboring *BCO26_mntH* from *B. coagulans 2-6* | This study |
| pUC19-*BCO26_2932* | pUC19 harboring *BCO26_2932* from *B. coagulans 2-6* | This study |
| pUC19-*BCO2_gabD* | pUC19 harboring *BCO26_gabD* from *B. coagulans 2-6* | This study |
| pUC19-*BCO2_groEL* | pUC19 harboring *BCO26_groEL* from *B. coagulans 2-6* | This study |
| pUC19-*BCO26_hrcA* | pUC19 harboring *BCO26_hrcA* from *B. coagulans 2-6* | This study |
| pUC19-*BCO26_clpE* | pUC19 harboring *BCO26_clpE* from *B. coagulans 2-6* | This study |
| pUC19-*BCO26_1771* | pUC19 harboring *BCO26_1771* from *B. coagulans 2-6* | This study |
| pUC19-*BCO26_dnaK* | pUC19 harboring *BCO26_dnaK* from *B. coagulans 2-6* | This study |
| pUC19-*BCO26_groES* | pUC19 harboring *BCO26_groES* from *B. coagulans 2-6* | This study |
| pUC19-*BCO26_0541* | pUC19 harboring *BCO26_0541* from *B. coagulans 2-6* | This study |
| pUC19-*BCO26_ykzI* | pUC19 harboring *BCO26_ykzI* from *B. coagulans 2-6* | This study |

| | | |
|---|---|---|
| pUC19-*BCO26_katE* | pUC19 harboring *BCO26_katE* from *B. coagulans 2-6* | This study |
| pUC19-*BCO26_0399* | pUC19 harboring *BCO26_0399* from *B. coagulans 2-6* | This study |
| pUC19-*BCO26_2461* | pUC19 harboring *BCO26_2461* from *B. coagulans 2-6* | This study |
| pUC19-*BCO26_levG* | pUC19 harboring *BCO26_levG* from *B. coagulans 2-6* | This study |
| pUC19-*BCO26_yqiG* | pUC19 harboring *BCO26_yqiG* from *B. coagulans 2-6* | This study |
| pUC19-*BCO26_2340* | pUC19 harboring *BCO26_2340* from *B. coagulans 2-6* | This study |
| pUC19-*BCO26_yflT* | pUC19 harboring *BCO26_yflT* from *B. coagulans 2-6* | This study |
| pUC19-*BCO26_argI* | pUC19 harboring *BCO26_argI* from *B. coagulans 2-6* | This study |
| pUC19-*BCO26_sucC* | pUC19 harboring *BCO26_sucC* from *B. coagulans 2-6* | This study |
| pUC19-*BCO26_2370* | pUC19 harboring *BCO26_2370* from *B. coagulans 2-6* | This study |
| pUC19-*BCO26_citZ* | pUC19 harboring *BCO26_citZ* from *B. coagulans 2-6* | This study |
| pUC19-*BCO26_rsbV* | pUC19 harboring *BCO26_rsbV* from *B. coagulans 2-6* | This study |
| pUC19-*BCO26_hag* | pUC19 harboring *BCO26_hag* from *B. coagulans 2-6* | This study |
| pUC19-*BCO26_yhgD* | pUC19 harboring *BCO26_yhgD* from *B. coagulans 2-6* | This study |
| pUC19-*BCO26_1679* | pUC19 harboring *BCO26_1679* from *B. coagulans 2-6* | This study |
| pUC19-*BCO26_rsbW* | pUC19 harboring *BCO26_rsbW* from *B. coagulans 2-6* | This study |
| pUC19-*BCO26_yteA* | pUC19 harboring *BCO26_yteA* from *B. coagulans 2-6* | This study |
| pUC19-*BCO26_2825* | pUC19 harboring *BCO26_2825* from *B. coagulans 2-6* | This study |
| pUC19-*BCO26_levE* | pUC19 harboring *BCO26_levE* from *B. coagulans 2-6* | This study |
| pUC19-*BCO26_sigB* | pUC19 harboring *BCO26_sigB* from *B. coagulans 2-6* | This study |
| pUC19-*BCO26_2573* | pUC19 harboring *BCO26_2573* from *B. coagulans 2-6* | This study |

| | | |
|---|---|---|
| pUC19-*BCO26_1484* | pUC19 harboring *BCO26_1484* from *B. coagulans 2-6* | This study |
| pET28a-*cspL* | pET28a harboring *BCO26_cspL* from *B. coagulans* 2-6 | This study |
| pET28a-*cspL*-M11 | pET-28a harboring *BCO26_cspL* from *B. coagulans 2-6* with mutated G14 Y15 G16 F17 I18 E19 R20 V26 F27 V28 H29 | This study |
| pET28a-*cspL*-M7 | pET-28a harboring *BCO26_cspL* from *B. coagulans 2-6* with mutated G14 Y15 G16 F17 I18 E19 R20 | This study |
| pYES2-*cspL* | pYES2 harboring *BCO26_cspL* from *B. coagulans 2-6* | This study |
| pME6032-*cspL* | pME6032 harboring *BCO26_cspL* from *B. coagulans 2-6* | Novagen |
| pUC19-*cspA* | pUC19 harboring *cspA* from *E. coli* | This study |
| pLQ856 | pDR3 derivative | Ref. 31 |
| pLQ856-*cspL* | pLQ856 derivative with inserted *cspL* under the control of kasOp* | This study |

[a]$Amp^R$ and $Kan^R$ resistance to ampicillin and kanamycin, respectively.

**Table S9. Sequences of primers used in this study.**

| Primer | Sequence (5' > 3') |
|---|---|
| CspL-F | CGCGGATCC atggaacatggtacagtaaa |
| CspL-R | CCGGAATTC ttagtcttctttttgaacat |
| BCO26_CspD-F | CGCGGATCC atgcaaaacggtaaagtaaa |
| BCO26_CspD-R | CCGGAATTC ttatgaaagttttgttacat |
| BCO26_YkuS-F | CGCGGATCC atggctgtaatcggtgtaga |
| BCO26_YkuS-R | CCGGAATTC ttacattctgctcgccactt |
| BCO26_2925-F | CCCAAGCTT atgaaaaagcgggcaattgt |
| BCO26_2925-R | CGCGGATCC tcaggaaaccactgcctttt |
| BCO26_Dps-F | CGCGGATCC atggcagaaaacgaacaatt |
| BCO26_Dps-R | CCGGAATTC ttaccgctttccaagaaagg |
| BCO26_GsiB-F | CCCAAGCTT atggcagacaaagataaaaa |
| BCO26_GsiB-R | CGCGGATCC ttaatcttcaccgttgtttt |
| BCO26_YbfB-F | CGCGGATCC atgttagaacgaaaagcaaa |
| BCO26_YbfB-R | CCGGAATTC ttaatgcgaatgctgggcac |
| BCO26_MntH-F | CCCAAGCTT atgagtgaaaaaatgatgag |
| BCO26_MntH-R | CCGGAATTC ttatataaacgtatcaatca |
| BCO26_2932-F | CCCAAGCTT atgagcatcagacagggaaa |
| BCO26_2932-R | CGCGGATCC ttaccggtaggccggttctt |
| BCO26_GabD-F | CGCGGATCC atggaagactatttgatgta |
| BCO26_GabD-R | CCGGAATTC ttataaaccgagggaaatat |
| BCO26_GroEL-F | CGCGGATCC atggcaaaagaaattaaatt |
| BCO26_GroEL-R | CCGGAATTC ttacatcatgccgcccatgc |
| BCO26_HrcA-F | CGCGGATCC atggcggatcttgaggaact |
| BCO26_HrcA-R | CCGGAATTC ctatctgtcatacaatttcg |
| BCO26_ClpE-F | CGCGGATCC atgttatgtgacaaatgcca |
| BCO26_ClpE-R | CCGGAATTC ttatttttccgcgatggcaa |
| BCO26_1771-F | CCCAAGCTT atgtttgatttaatgccatt |
| BCO26_1771-R | CGCGGATCC ttattgaatttcaatccttt |
| BCO26_DnaK-F | CGCGGATCC atgagcaaaattatcggcat |
| BCO26_DnaK-R | CCGGAATTC ttattttttattatcatcga |
| BCO26_GroES-F | CGCGGATCC atgtttcacgtgttaaaacc |
| BCO26_GroES-R | CCGGAATTC ttattccacaaccgccagaa |
| BCO26_0541-F | CGCGGATCC atggtaggaattatcattgc |
| BCO26_0541-R | CCGGAATTC ttatttttgtttgttcagct |
| BCO26_YkzI-F | CGCGGATCC atgaaacaagtaatcccttc |
| BCO26_YkzI-R | CCGGAATTC ttacatggctttcattttca |
| BCO26_KatE-F | CTAGTCTAGA atgagtagtgaacggaaact |
| BCO26_KatE-R | CGAGCTC tcatatcaaacgcctgtccc |
| BCO26_0399-F | CCCAAGCTT atgccattggaactggtaat |
| BCO26_0399-R | CTAGTCTAGA tcaatgctttcccccctcta |
| BCO26_2461-F | CGCGGATCC atgatgaaaaaatcaatggc |
| BCO26_2461-R | CCGGAATTC tcactttttcgcgcctgcaa |

| | |
|---|---|
| BCO26_LevG-F | CCCAAGCTT atggcacaagaactaaaatt |
| BCO26_LevG-R | CTAGTCTAGA ttacattaagtgaattaaat |
| BCO26_YqiG-F | CGCGGATCC atgagcaaatacgataaact |
| BCO26_YqiG-R | CCGGAATTC tcactctgcaaacgggaacc |
| BCO26_2340-F | CCCAAGCTT atgaaccgaaatttttgtaa |
| BCO26_2340-R | CGCGGATCC ttatctttcggcactccgca |
| BCO26_YflT-F | CGCGGATCC atgcataaagtagaagtggt |
| BCO26_YflT-R | CCGGAATTC ttatagcaggtgttcaggcc |
| BCO26_ArgI-F | CGCGGATCC atggagaaacatattgcaat |
| BCO26_ArgI-R | CCGGAATTC ttaaagaagtttttcccga |
| BCO26_SucC-F | CGCGGATCC atgaatattcacgagtatca |
| BCO26_SucC-R | CCGGAATTC ttagctgaccagttcgacaa |
| BCO26_2370-F | CGCGGATCC atgagatcgattgtaaatga |
| BCO26_2370-R | CCGGAATTC ttaatacagcgtaatttcct |
| BCO26_CitZ-F | CGCGGATCC atgacagcaacaagaggtct |
| BCO26_CitZ-R | CCGGAATTC ttaccggtcttcgagcggaa |
| BCO26_RsbV-F | CGCGGATCC atggactttgaagtagatgt |
| BCO26_RsbV-R | CCGGAATTC tcacactccaccttctattt |
| BCO26_Hag-F | CCCAAGCTT atgattatcaatcacaacat |
| BCO26_Hag-R | CGCGGATCC ttaacgcaacaattgcaata |
| BCO26_YhgD-F | ACATGCATGC atggaaacggaccgcaggct |
| BCO26_YhgD-R | CTAGTCTAGA tcagttctgcagccctttta |
| BCO26_1679-F | CTAGTCTAGA atgtctgttgcttcgacaga |
| BCO26_1679-R | TCCCCCGGG ttacaagccgcattttaatt |
| BCO26_RsbW-F | CGCGGATCC atggaggagtttgatcatat |
| BCO26_RsbW-R | CCGGAATTC tcaggttgaggcagttttga |
| BCO26_YteA-F | CCCAAGCTT atgctgacaaaagaacaact |
| BCO26_YteA-R | CGCGGATCC tcatttcttttttttctcgt |
| BCO26_2825-F | CGCGGATCC atgggggcgattcagatcat |
| BCO26_2825-R | CCGGAATTC tcatgcccggcccccccctt |
| BCO26_LevE-F | CGCGGATCC atggcattggatatacggct |
| BCO26_LevE-R | CCGGAATTC tcatggatgaagcagtttat |
| BCO26_SigB-F | CGCGGATCC atgtcaaaactgcctcaacc |
| BCO26_SigB-R | CCGGAATTC ctaattctccacatgttgca |
| BCO26_2573-F | ACATGCATGC atggaaaacattaaaatgct |
| BCO26_2573-R | CTAGTCTAGA tcaatgcttttccccctcta |
| BCO26_1484-F | CGCGGATCC atggaacaaggtaaagtaaa |
| BCO26_1484-R | CCGGAATTC ttataatttcgaaacgttcg |
| pET28a-CspL-F | CATGCCATGGca atggaacatggtacagtaaa |
| pET28a-CspL-R | CCGCTCGAG ttagtcttctttttgaacat |
| CspA-F | CGCGGATCC atgtccggtaaaatgactgg |
| CspA-R | CCGGAATTC ttacaggctggttacgttac |
| pYES2-CspL-F | CGCGGATCC atggaacatggtacagtaaa |
| pYES2-CspL-R | CCGGAATTC ttagtcttctttttgaacat |

| | |
|---|---|
| pME6032-CspL-F | CCGGAATTC atggaacatggtacagtaaa |
| pME6032-CspL-R | CATGCCATGG ttagtcttctttttgaacat |
| 77-cspL-F | cggttggtaggatccacatatggagcacggcaccgtgaagt |
| 77-cspL-R | tatgacatgattacgaattcagtcctccttctgcacgttggcg |

**Table S10. Sequences of oligonucleotide used in this study.**

| RNA | Sequence (5’ > 3’) |
|---|---|
| RNA 1 | CGGGAGAGGCGGUUUGCGUAUUGU |
| RNA 2 | GGAGAGGCGGUUUGCGUAUUGU |
| RNA 3 | GGGAGAGGCGGUUUGCGUAU |
| RNA 4 | GGAGAGGCGGUUUGCGU |
| RNA 5 | GCAUUAAUGAAU |
| RNA 6 | GCAUUAAUGAA |
| RNA 7 | CAUUAAUGAA |
| RNA 8 | AUUAAUGAAU |
| RNA 9 | AUUAAUGA |
| RNA 10 | UAAUGA |
| RNA 11 | UAAUG |
| RNA 12 | AAUG |
| RNA 13 | GAAUG |
| RNA 14 | TAAUG |
| RNA 15 | AAUGC |
| ssDNA 1 | AGGTACCCGGGGATCCTCTAGAGTCGTC |
| ssDNA 2 | TGTACCCGGGGATCCTCTAGAGTCGC |
| ssDNA 3 | GTACCCGGGGATCCTCTAGAGTCG |
| ssDNA 4 | GGGAGACCGGAATTCGAGCTCG |
| ssDNA 5 | GAGACCGGAATTCGAGCT |
| ssDNA 6 | GACCGGAATTCGAG |
| ssDNA 7 | CCGGAATTCG |
| ssDNA 8 | GGAATT |
| ssDNA 9 | GAAT |
| ssDNA 10 | GAATC |
| ssDNA 11 | CGAAT |
| ssDNA 12 | TGAAT |

**Table S11.** Synthetic gene sequences used in this study.

| Name | Sequence (5′ > 3′) |
|---|---|
| *cspL*-M11 | ccatggca gaa cat ggt aca gta aaa tgg ttt aac agt gaa aaa gca gca gca gca gca gca gca gaa ggc gga gac gac gca gca gca gca ttc tcg gcc atc cag ggt gaa ggc tat aaa acg ctt gaa gaa ggc cag aaa gta tca ttt gat gtg gaa gaa gga tca cgc ggc ccg cag gcg gca aat gtt caa aaa gaa gac ctcgag |
| *cspL*-M7 | ccatggca gaa cat ggt aca gta aaa tgg ttt aac agt gaa aaa gca gca gca gca gca gca gca gaa ggc gga gac gac gtg ttt gtc cat ttc tcg gcc atc cag ggt gaa ggc tat aaa acg ctt gaa gaa ggc cag aaa gta tca ttt gat gtg gaa gaa gga tca cgc ggc ccg cag gcg gca aat gtt caa aaa gaa gac ctcgag |
